# Dynamic regulation of gonadal transposon control across the lifespan of the naturally short-lived African turquoise killifish

**DOI:** 10.1101/2022.09.02.506427

**Authors:** Bryan B. Teefy, Ari Adler, Alan Xu, Katelyn Hsu, Param Priya Singh, Bérénice A. Benayoun

## Abstract

Although germline cells are considered to be functionally “immortal”, both the germline and supporting somatic cells in the gonad within an organism will experience aging. With increased age at parenthood, the age-related decline in reproductive success has become an important biological issue for an aging population. However, molecular mechanisms underlying reproductive aging across sexes in vertebrates remain poorly understood. To decipher molecular drivers of vertebrate gonadal aging across sexes, we perform longitudinal characterization of the gonadal transcriptome throughout lifespan in the naturally short-lived African turquoise killifish *(Nothobranchius furzeri)*. By combining mRNA-seq and small RNA-seq from 26 individuals, we characterize the aging gonads of young adult, middle-aged, and old female and male fish. We analyze changes in transcriptional patterns of genes, transposable elements (TEs), and piRNAs. We find that testes seem to undergo only marginal changes during aging. In contrast, in middle-aged ovaries, the timepoint associated with peak female fertility in this strain, PIWI pathway components are transiently downregulated, TE transcription is elevated, and piRNA levels generally decrease, suggesting that egg quality may already be declining at middle-age. Furthermore, we show that piRNA ping-pong biogenesis declines steadily with age in ovaries, while it is maintained in aging testes. To our knowledge, this dataset represents the most comprehensive transcriptomic dataset for vertebrate gonadal aging. This resource also highlights important pathways that are regulated during reproductive aging in either ovaries or testes, which could ultimately be leveraged to help restore aspects of youthful reproductive function.

## Introduction

In much of the industrialized world, parental age is increasing making fertility-related challenges associated with later-life childbearing increasingly relevant (Kenny et al. 2013; Bertoldo et al. 2020). In humans, oocyte quality begins to rapidly degrade around 30 years old while menopause, the irreversible loss of female fertility, occurs around 50 years old (Alberts et al. 2013; Finch 2014). In contrast, while male fertility gradually declines with age, complete loss of testicular function is not a feature of male reproductive aging (Gunes et al. 2016). Mammals, as well as birds, are exceptional among vertebrates as these groups produce a finite supply of oocytes before or shortly after birth ensuring that any species that lives long enough should eventually deplete its oocytes, although the starting supply of oocyte greatly outnumbers the final number of oocytes that will be ovulated (Mira 1998). However, there are many other mechanisms that influence reproductive decline, such as neuroendocrine aging, that occur in both sexes and are widely conserved among vertebrates (Perheentupa and Huhtaniemi 2009). Thus, leveraging a tractable vertebrate model organism to understand molecular mechanisms underlying reproductive aging will provide important insights into lifelong regulation of reproductive function.

To study reproductive aging, we examined the impact of aging on testes and ovaries in the African Turquoise Killifish *Nothobranchius furzeri*, the shortest-lived vertebrate that can be bred in captivity, with a lifespan of about 4-6 months (Hu and Brunet 2018). The turquoise killifish evolved this short lifespan in response to their unique lifecycle, which revolves around the formation of ephemeral ponds of water (Kim et al. 2016). Depending on the specific species, killifish can become sexually mature as early as two weeks post-hatching, although most killifish become sexually mature around 4-5 weeks of age, the fastest known time to sexual maturation of any vertebrate (Naumann and Englert 2018; Vrtilek et al. 2018b). Killifish are asynchronous breeders, and upon sexual maturity, females continuously undergo oogenesis and lay eggs, typically on a daily basis (Terzibasi Tozzini and Cellerino 2020). While male reproductive senescence may be negligible, females do experience an age-related decline in fecundity, though the molecular mechanisms contributing to reproductive aging in either sex have not been elucidated (Vrtilek et al. 2018a; Zak and Reichard 2021). Importantly, we and others have helped establish a key functional genomics toolkit for this species, making it a uniquely tractable experimental model to decipher molecular mechanisms driving aspects of aging (Valenzano et al. 2011; Reichwald et al. 2015; Valenzano et al. 2015; Harel et al. 2016; Willemsen et al. 2020).

An emerging facet of vertebrate aging is the reactivation of transposable elements (TEs) with aging in somatic tissues (De Cecco et al. 2013; Chen et al. 2016; Simon et al. 2019; LaRocca et al. 2020; Yang et al. 2022). However, whether TEs also transcriptionally activate within the aging gonad, which is uniquely protected from spurious TE mobilization by the PIWI-piRNA pathway, has been largely unexplored in a vertebrate system. The PIWI pathway is an Argonaute-based small RNA pathway that maintains germline genomic integrity by antagonizing the mobilization of transposable elements (TEs) through complementary RNA base-pairing and TE RNA target degradation during gametogenesis and embryogenesis. PIWI family proteins bind 24-35 nucleotide (nt) <u>P</u>IWI-interacting <u>RNA</u>s, or piRNAs, which are complementary to or derived from TEs. piRNAs are expressed primarily in the gonad and are distinctly larger than miRNAs (Mani and Juliano 2013; Czech and Hannon 2016). Once bound to an RNA target, PIWI proteins cleave the RNA target precisely 10 nts from the 5’ end of the complementary piRNA (Mani and Juliano 2013; Czech and Hannon 2016). To date, most studies examining the impact of aging on the PIWI pathway have focused on invertebrate model organisms, such as the fly *Drosophila melanogaster* (Sousa-Victor et al. 2017; Yamashiro and Siomi 2018; Lin et al. 2020). For instance, Piwi expression is required to suppress TE expression, reduce DNA damage, and maintain intestinal stem cell lineages in the aging Drosophila midgut (Sousa-Victor et al. 2017). In the fly ovary, age-related decline in *Piwi* expression in the cells of the germline stem cell niche results in a transposon de-repression and a loss of germline stem cells (GSCs) (Lin et al. 2020). In contrast to the germline stem cell niche, the expression of PIWI pathway components in whole *Drosophila* egg chambers increases with age, although this is not associated with any global changes in TE expression (Erwin and Blumenstiel 2019).

Whether TEs are derepressed with age in the vertebrate gonad and whether TE expression is modulated by age-related PIWI dynamics remains unknown. Thus, it will be important to determine whether PIWI regulation (and concomitant TE control) in the aging gonad may influence reproductive aging.

In this study, we leverage mRNA and small RNA sequencing to capture the transcriptional trajectory of gonadal aging in a naturally short-lived vertebrate. To do so, we characterize the transcriptome of African turquoise killifish ovaries and testes in young adults (5 weeks), middle-age (10 weeks) and old (15 weeks) animals in the GRZ inbred lab strain. In particular, we focus on the interplay of TEs and piRNAs, and assess the effect of age on the transcriptional status of these classes of RNAs.

## Results

### Lifespan, reproductive, and transcriptomic landscapes of the aging gonad in the African turquoise killifish

To understand how gonads are affected during aging in a vertebrate model organism, we performed a study of ovarian and testicular landscapes throughout aging in African turquoise killifish from the GRZ strain (**Fig. 1A**). In our housing conditions (see Methods), 15 weeks correspond to ∼90% survival for both female and male animals (**Fig. 1B; Supplemental Table S1A**), although GRZ females lived significantly longer than males in our husbandry conditions (p = 0.002, log Rank test; see Methods) (**Fig. 1B**). To determine the effect of aging on the vertebrate gonadal transcriptomic landscape, we collected ovaries and testes from N = 5 GRZ female and male African turquoise killifish at three different time points post-hatching: young adulthood (5 weeks; onset of fertility), middle-age (10 weeks), and old age (15 weeks; before any dramatic population survival decrease) (**Fig. 1A**). The choice of these specific time points is consistent with broadly accepted guidelines in aging research, to avoid measuring development or survivorship bias (Flurkey et al. 2007) (see methods). In the highly inbred GRZ strain of African turquoise killifish, fecundity was reported to peak around 10-12 weeks of age (Zak and Reichard 2021), roughly corresponding to the middle-age time point that we selected. Only samples with intact RNA (RIN >7) were processed further to eliminate biases linked to RNA degradation, yielding 14 female samples and 12 male samples, leaving at least 4 samples per biological group (**Supplemental Fig. S1A**). We performed (i) mRNA transcriptome characterization (*i.e.* mRNA-seq using polyA selection), analyzing both transcriptional patterns of genes and transposable elements [TEs], and (ii) small-RNA transcriptome characterization (*i.e.* small RNA-seq), focusing on piRNAs, the germline-specific class of small RNAs of the PIWI pathway (**Fig. 1A** and **Supplemental Fig. S1A**).

**Figure 1.**
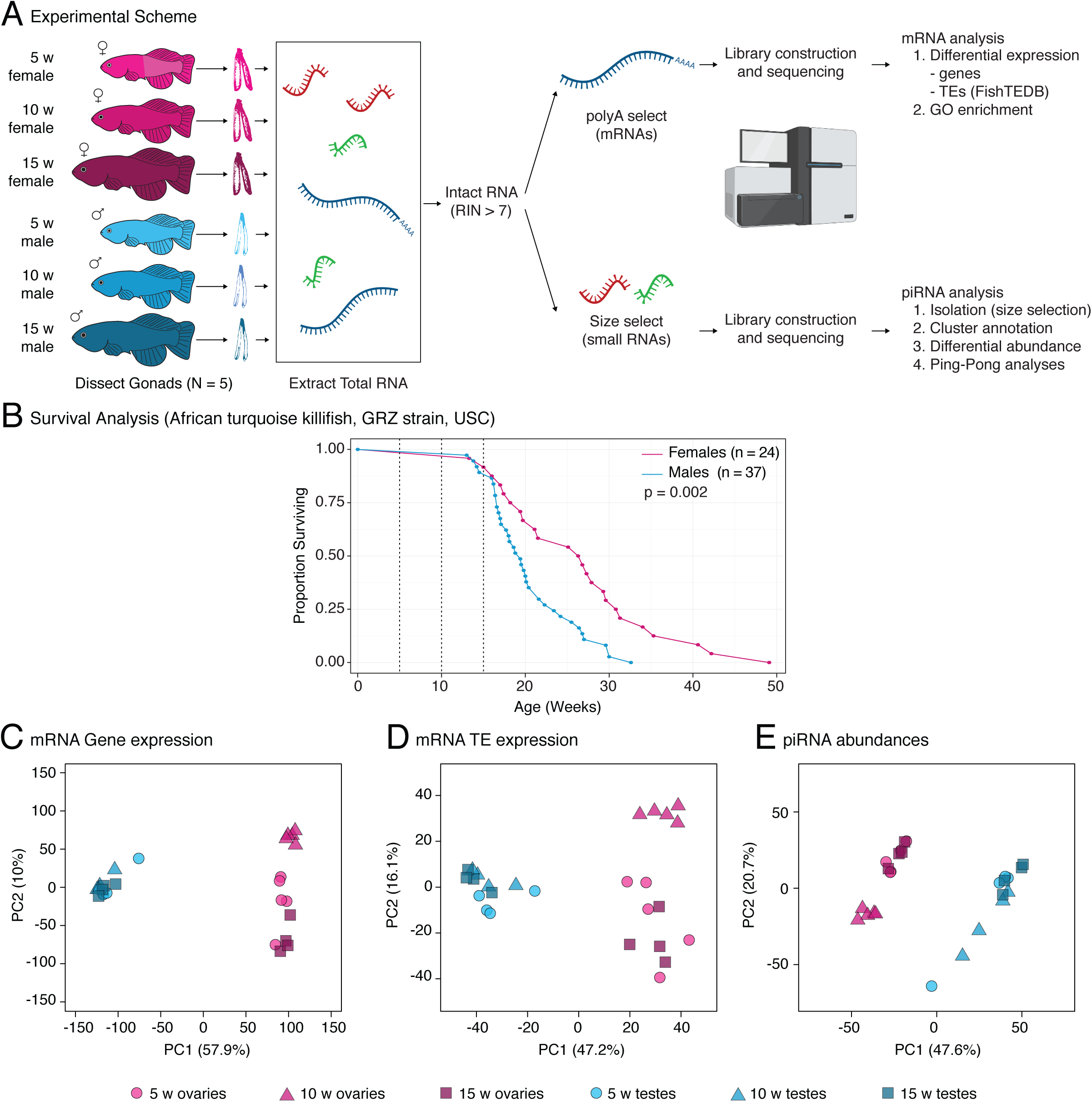
A study of gonadal aging in the naturally short-lived African turquoise killifish. **(A)** Experimental scheme for killifish gonad sequencing. Killifish gonads (N = 5) were dissected and total RNA was extracted. Only high quality RNA samples (RIN>7) were included for further processing (see **Supplemental Fig. S1A**). Small RNA and mRNA RNA-seq libraries were derived from each total RNA sample and sequenced for downstream analyses. **(B)** Lifespan curve for female and male GRZ strain killifish at the USC Benayoun lab facility. GRZ females lived significantly longer than males (p = 0.002; log Rank test). Median lifespan of females: 26.55 weeks, males: 19.4 weeks. Vertical dotted lines: ages sampled in this study (5, 10, 15 weeks), chosen to represent aging before substantially decreased population survival. **(C-E)** Principal component analyses [PCAs] for **(C)** mRNA gene expression **(D)** TE expression and **(E)** TE-mapping piRNA abundance (see methods).

To globally visualize the similarity of the transcriptomic landscapes of aging ovaries and testes at multiple levels of regulation, we used principal component analysis [PCA] (**Fig. 1C-E**). PCA analysis of genic mRNA, TE mRNA, and TE-targeting piRNA expression revealed that the main source of variation [PC1] corresponded to gonadal identity (*i.e.* ovary vs. testis), although animal age also weakly separated samples for mRNA and piRNA cluster (but not TEs) transcription on PC2 (**Fig. 1C-E**). Notably, despite genomic TE content being a fixed feature for the species, we observed clear sex-dimorphism in the transcriptional landscape of TEs and associated piRNAs in the turquoise killifish gonads. These observations are consistent with deeply sex-dimorphic transcriptional programs in the gonad at the level of mRNAs, TEs and small RNAs, and are consistent with a recent study in young *Drosophila* ovaries and testes (Chen et al. 2021). Separately, we analyzed the ovarian and testicular transcriptomes to uncover potential sex-specific transcriptional patterns with aging (**Supplemental Fig. S1B-S**; see supplementary methods). Using PCA, we noted that middle-aged female expression data clustered distinctly for all analyzed RNA species in PC1 (**Supplemental Fig. S1B-D**). This observation may reflect the outsized contribution of a transcriptional program associated with oogenesis, or fertility more broadly, to the expression profiles of all RNA species assayed at this timepoint. Intriguingly, female piRNA expression data segregated most strongly by age, suggesting that the small RNA program in the ovary is most sensitive to aging (**Supplemental Fig. S1D**). In contrast, the male transcriptomic data did not visually separate strongly by age (**Supplemental Fig. S1E-G**).

Consistent with our PCA results, we noted stronger clustering of samples by age for ovaries compared to testes for gene, TE and piRNA expression using hierarchical clustering with bootstrap resampling by ‘pvclust’ (Suzuki and Shimodaira 2006) (**Supplemental Fig. S1H-M**; see supplementary methods). Finally, we also quantified the proportion of gene, TE and piRNA expression variance explained by age in the ovaries and testes using the ‘variancePartition’ framework (Hoffman and Schadt 2016) (**Supplemental Fig. S1N-S**; see supplementary methods). Importantly, the median percentage of variance explained by age was systematically higher in ovaries than testes (genes: ∼25% *vs.* ∼6%; TEs: ∼15% *vs.* ∼6; piRNAs: ∼49% *vs.* ∼2%; **Supplemental Fig. S1N-S**). Together, our observations suggest that testicular identity and function are more stable in response to organismal aging relative to the ovary. Interestingly, our results are reminiscent of a microarray analysis of gonadal aging in mice, where ovaries underwent substantially more transcriptional changes with aging than testes (Sharov et al. 2008). Our transcriptomic data is also consistent with observations of negligible male reproductive aging in this species (Vrtilek et al. 2018a; Zak and Reichard 2021). Together, our data suggests unique changes in the transcriptional landscapes of ovaries versus testes throughout aging.

### Aging differentially impacts gene expression in turquoise killifish ovaries and testes

Since aging is often a non-linear process, at least in somatic tissues (Baumgart et al. 2016; Rosenberg et al. 2021), we decided to determine differential gene expression with aging within the DEseq2 framework using a likelihood ratio testing [LRT] approach, so as to unbiasedly identify <u>patterns</u> of differential gene expression in ovary and testis aging (Love et al. 2014). To note, since LRT can be overly lenient (Core 2021), we used a stringent False Discovery Rate [FDR] threshold of FDR <1e-6 to identify differentially expressed transcripts. Significantly regulated transcripts were then assigned to patterns using ‘degPatterns’ from the ‘DEGreport’ R package (Pantano 2022), which uses hierarchical clustering of gene expression to identify shared significant patterns of expression among differentially expressed genes in an unsupervised fashion. This unbiased approach detected four major differential patterns corresponding to transcripts whose expression is (a) transiently down at middle-age, (b) transiently up at middle-age, (c) monotonously down with age, or (d) monotonously up with age (**Fig. 2A-B**, **3A**, and **Supplemental Fig. S2A**, **S3A**; **Supplemental Table S2A-B,D**). For reading convenience, we label these age-related patterns of differential expression hereafter as (a), (b), (c) and (d) whenever they arise from the unsupervised ‘degPatterns’ clustering. Intriguingly, we noted the majority of differentially expressed genes showed transient changes at middle-age in both ovaries and testes (respectively 1931 out of 2041, and 578 out of 603 differentially expressed genes), rather than linear trends with age. This trend suggests that most of the transcriptional variation in gonadal gene expression occurs at middle-age when fertility, at least in females, is at its peak. This observation is interesting, because it suggests that gonadal biology is already substantially impacted in middle-age healthy animals.

**Figure 2.**
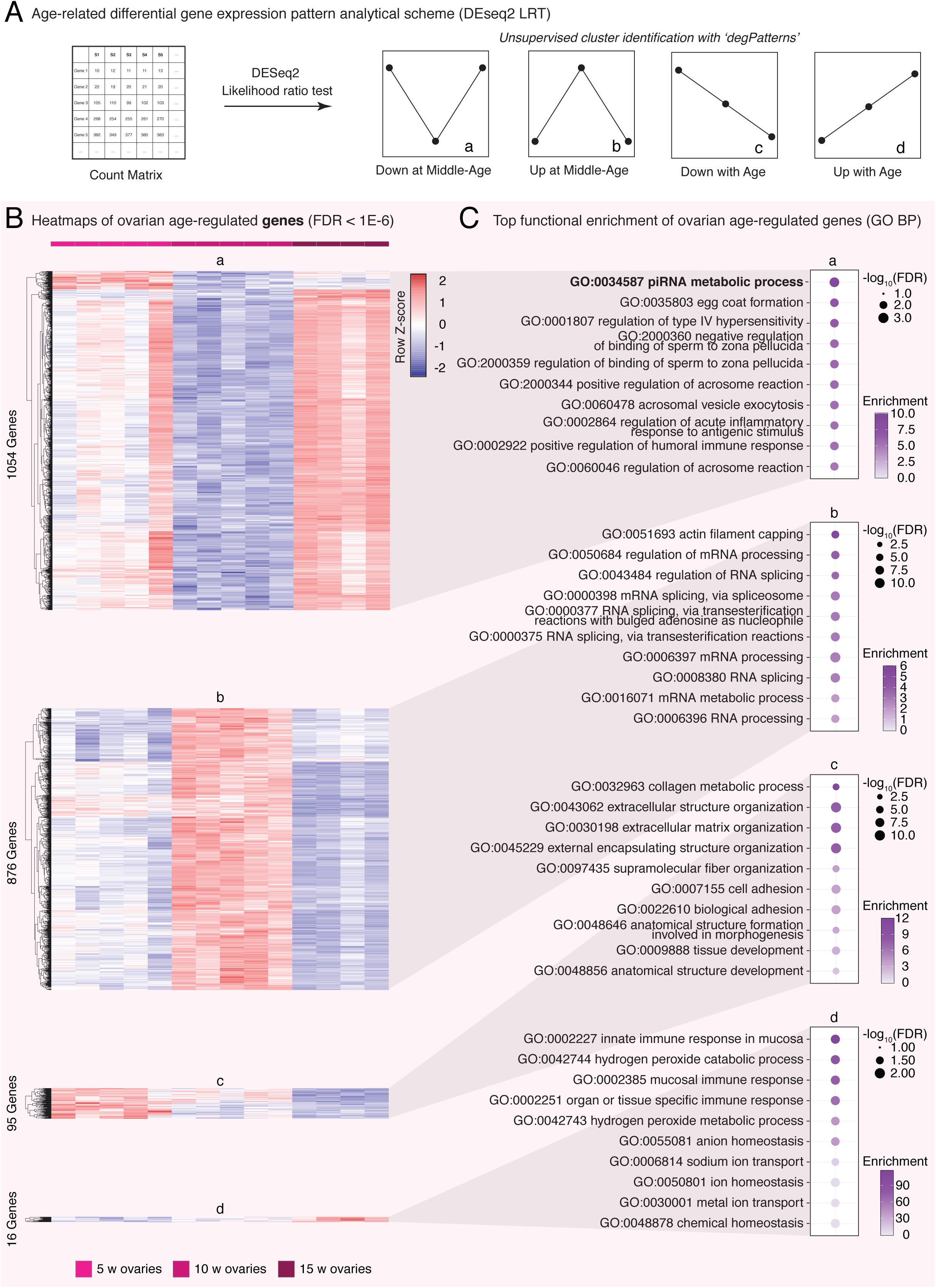
Differential gene expression analysis and GO Biological Process functional enrichment analysis with ovarian aging. **(A)** Scheme used for differential gene expression [DGE] analysis used in this study. DGE was performed with DESeq2’s likelihood ratio testing [LRT], which revealed 4 approximate groups of differentially expressed genes corresponding to expression (a) Down at Middle-Age, (b) Up at Middle-Age, (c) Down with Age and (d) Up with Age. **(B)** Heatmaps of gene expression for genes significantly regulated with age in aging killifish ovaries by DEseq2 LRT (FDR < 1e-6). Most differentially expressed genes transiently changed at middle-age. **(C)** Functional enrichment analysis for each gene cluster in Fig. 2B, showing the top 10 most significant GO “Biological Process” terms by ‘GOstats’ (FDR < 5%; see **Supplemental Table S3A** for complete list of enriched terms). The most significant term downregulated in middle-aged ovaries is “piRNA metabolic process” (bolded). FDR: False discovery rate. Enrichment: fold enrichment over background.

Next, we asked which biological functions were associated with each of these gene expression patterns with gonadal aging. For this purpose, we performed Gene Ontology [GO] Enrichment Analysis for each age-regulated pattern in ovaries and testes (Falcon and Gentleman 2007) (**Fig. 2C**, **3B**, and **Supplemental Fig. S2B, S3B**; **Supplemental Table S3A-F**). First, we examined enrichment for terms in the “Biological Process” GO category (**Fig. 2C**, **3B**; **Supplemental Table S3A,D**). Interestingly, genes downregulated in middle-aged ovaries were enriched most strongly for “piRNA metabolic process” (*i.e.* pattern a; GO:0034587; **Fig. 2C**; **Supplemental Table S3A**). Specifically, genes transcriptionally downregulated in middle-age ovaries include the genes encoding both effector PIWI proteins (*i.e.* the turquoise killifish homologs to human *PIWIL1* and *PIWIL2*), is consistent with overall decreased TE processing capacity. To note, as seen in **Supplemental Table S1C**, genes belonging to the “piRNA metabolic process” GO terms are, as expected, robustly expressed in killifish gonads throughout life.

**Figure 3.**
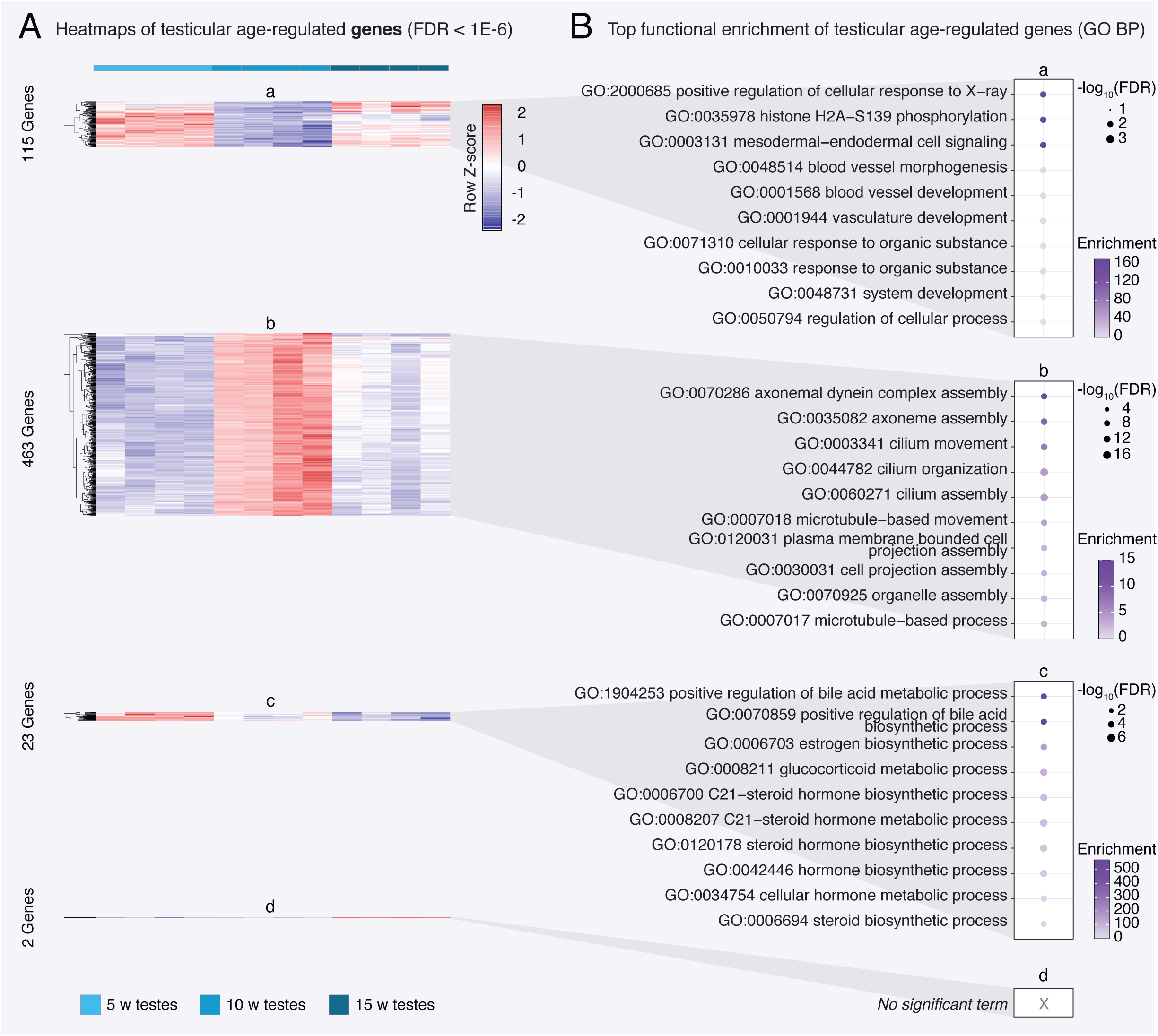
Differential gene expression and GO Biological Process functional enrichment analysis with testicular aging. **(A)** Heatmaps of gene expression for genes significantly regulated with age in aging killifish testes by DEseq2 LRT (FDR < 1e-6). Groups correspond to those defined in Fig. 2A. Like ovaries, most differential gene expression in killifish testes occurs transiently at middle-age. **(B)** Functional enrichment analysis for each gene cluster shown in Fig. 3A showing the top 10 most significant GO “Biological Process” terms by ‘GOstats’ (FDR < 5%; see **Supplemental Table S3D** for complete list of enriched terms). Genes that are upregulated in middle-age testis are enriched for terms related to spermatogenesis. Note that cluster (d) (genes up with age) had too few genes to generate meaningful enrichment results. FDR: False discovery rate. Enrichment: fold enrichment over background.

Intriguingly, in zebrafish, the gene encoding PIWIL1, the primary catalytic protein of the PIWI pathway, is expressed most highly in developing germline stem cells before expression decreases in more mature oocytes (Liu et al. 2022), which could indicate a higher ratio of mature oocytes to developing germline stem cells in middle-aged killifish ovaries compared to young and old ovaries. Thus, downregulation of PIWI-related genes at mid-age may be related to differences in mature oocytes content. To test this hypothesis, we performed (i) a small-scale histological analysis of oocyte diameter in ovaries from fish at different ages (since oocyte size is directly linked to maturation) (Api et al. 2018) (**Supplemental Fig. S4A-B**; see supplementary methods), and (ii) used a bulk transcriptome deconvolution approach using a zebrafish single-cell RNA-seq dataset of young adult ovaries as a reference dataset (**Supplemental Fig. S4C-G**; see supplementary methods). First, histological analysis revealed that the diameter distribution of oocytes in aging African turquoise killifish ovaries, measured by 4 independent blinded observers, did not vary significantly with age (**Supplemental Fig. S4B**). Second, whereas genes encoding PIWI-related are sharply downregulated at middle-age, deconvolution analysis did not detect substantial changes at middle-age in the proportion of immature germ cells in young *vs.* middle-aged turquoise killifish ovaries (**Supplemental Fig. S4F,G**). Thus, it is unlikely that observed bulk transcriptome changes in the expression of genes involved in piRNA metabolism in the middle-aged killifish ovary reflect substantial changes in the underlying ovarian composition of oocyte maturation stages. Alternatively, this may also suggest that there is a relaxation of piRNA-mediated TE expression control in ovaries at middle-age in the African turquoise killifish (see below). Other terms enriched for genes downregulated in middle-age ovaries were largely related to egg maturation and egg-specific processes (*e.g.* zona pellucida and regulation of acrosome reaction; **Fig. 2C**), consistent with the notion that ovarian function (and oocyte quality) may have already started to decline at middle-age in the African turquoise killifish.

Genes that were upregulated in middle-aged ovaries were enriched for terms related to RNA metabolism, consistent with conserved importance of maternally deposited RNAs in oocytes (*i.e.* pattern b; **Fig. 2C**). Genes that were linearly downregulated with age in the ovaries were enriched for terms relating to extracellular matrix organization (*i.e.* pattern c; **Fig. 2C**). Finally, genes that were upregulated with age in the ovaries included those involved in immune responses and ion homeostasis (*i.e.* pattern d; **Fig. 2C**), which may relate to overall increases in inflammation with aging that has been observed in somatic tissues (Benayoun et al. 2019). Increased expression of inflammatory genes has also been observed in the transcriptome of aging mouse ovaries (Sharov et al. 2008). We also performed a similar functional enrichment analysis using terms from the “Cellular Component” (**Supplemental Fig. S2B; Supplemental Table S3B**). and “Molecular Function” (**Supplemental Table S3C**).GO categories, with largely consistent trends.

In aging testes, we observed the emergence of a similar trend, with most differential genes being regulated at middle-age (*i.e.* patterns a and b), and few genes linearly regulated with aging (*i.e.* patterns c and d; **Fig. 3A**; **Supplemental Table S3D**). Genes transiently downregulated at middle-age were enriched for terms relating to vascular development (pattern a; **Fig. 3B**). This observation suggests that the vasculature of the testis may be actively remodeled, consistent with known unique vascular requirements for testicular function (Sargent et al. 2015). Genes transiently upregulated at middle-age included those involved in spermatogenesis, *e.g.* “axoneme assembly” and “cilium movement”, consistent with peak spermatogenesis potentially occurring at this stage (pattern b; **Fig. 3B**). Lastly, genes that were downregulated linearly with age were heavily enriched for steroid biosynthesis related terms, suggestive of a steady decline in steroid biosynthesis in the aging gonad (pattern c; **Fig. 3B**). Interestingly, this transcriptional downregulation of steroid biosynthesis genes is consistent with a recent mass-spectrometry study that detected significantly decreased levels of sex-steroids in the testes of aging turquoise killifish (Dabrowski et al. 2020). “Cellular component” analysis again corroborated the GO “Biological Process” analysis, especially with regards to sperm-related terms being upregulated at middle-age (**Supplemental Fig. S3B**; **Supplemental Table S3E**). Terms downregulated at middle-age included those associated with heterochromatin and extracellular vesicles, while terms downregulated linearly with age are enriched for mitochondrial and collagen related terms (**Supplemental Fig. S3B**; **Supplemental Table S3E**).

Together, our results show that the African turquoise killifish gonadal transcriptional landscapes are broadly remodelled at middle-age, highlighting the need to include middle-age time points in such analyses to get a full view of age-related trajectories. Notably, the largest changes in gene expression seem to align with the window of peak fertility and egg production in this species, suggesting that egg quality may start declining at middle-age despite high egg output.

### Transposon expression is age- and sex-dependent in the turquoise killifish gonad

Next, we asked whether control of TE transcription may be impacted during gonadal aging in the turquoise killifish. Indeed, accumulating evidence has shown that TE expression tends to increase with age in somatic tissues across species (De Cecco et al. 2013; Chen et al. 2016; Benayoun et al. 2019; Simon et al. 2019; Bravo et al. 2020; LaRocca et al. 2020; Yang et al. 2022). Although TE control is especially important in the germline to maintain genome stability throughout generations, whether TE expression escapes control in gonadal tissues with vertebrate aging has not yet been investigated. Interestingly, a recent genomic analysis of several species of African killifishes suggests that the shorter-lived killifish species have been accumulating TEs in their genomes, suggesting that short-lived/annual killifish species, such as the African turquoise killifish, may not efficiently repress germline TE activity throughout life (Cui et al. 2020).

To determine whether global changes in TE expression can indeed occur in aging killifish gonads, we first analyzed what proportion of mapped reads in our mRNA-seq dataset were derived from TE sequences (**Fig. 4A-B**). TE-derived sequences represented approximately 7-11% of reads in turquoise killifish ovaries and testes (**Fig. 4B**). Interestingly, global levels of TE-derived mRNA sequences were stable between young and middle-aged ovaries, but significantly dipped in old ovaries, whereas they tended to proportionally increase with age in testes (**Fig. 4B**). Such a pattern may reflect the process of gametogenesis, wherein the epigenome is repatterned, resulting in TE reactivation (Ben Maamar et al. 2021), despite the expected antagonizing effects of the PIWI pathway on TE expression in the germline.

**Figure 4.**
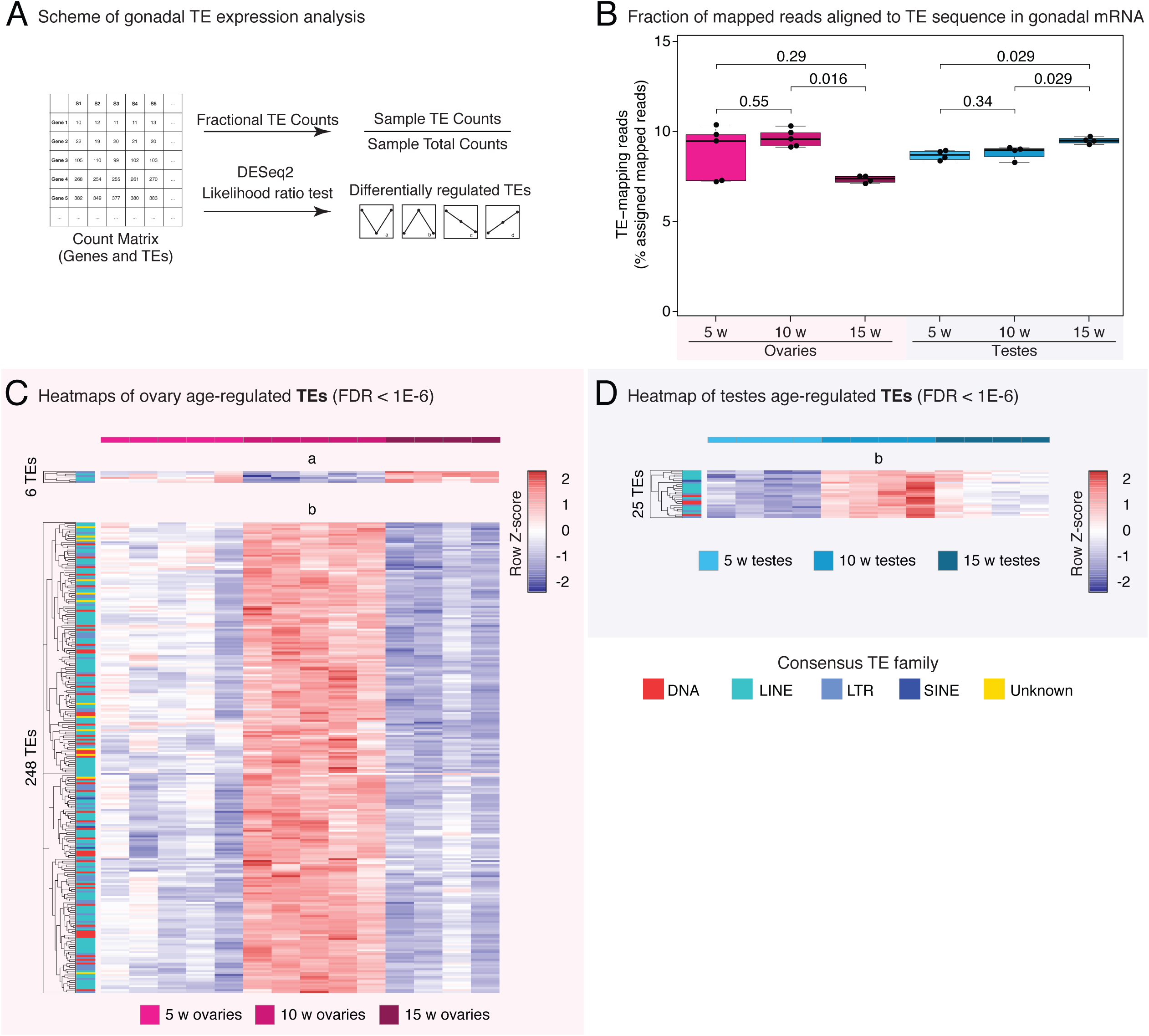
TE expression dynamics in the aging turquoise killifish gonad. **(A)** Experimental scheme for TE transcriptional analysis in aging gonads. Fractional TE counts were computed by counting the number of TE counts over total assigned mRNA-seq counts (genes and TEs). TE DGE was conducted in the same manner as the genic DGE analysis (see methods), and were labelled with the cluster nomenclature from Fig. 2A. **(B)** Proportion of assigned reads allocated to TEs from each mRNA-seq library for ovaries and testes. Significance in non-parametric Wilcoxon Rank Sum test. Note that old ovaries have significantly fewer proportional TE counts relative to middle-aged ovaries, and that testes seem to show a marginal increase in TE transcription with aging. **(C)** Heatmaps of mRNA expression for TEs significantly regulated with age in aging killifish ovaries by DEseq2 LRT (FDR < 1e-6). Each TE family is color-coded by row, revealing no obvious bias or enrichment for a specific TE type. **(D)** Heatmaps of mRNA expression for TEs significantly regulated with age in aging killifish testes by DEseq2 LRT (FDR < 1e-6). The only age differentially regulated TEs in the turquoise killifish testis are transiently upregulated at middle-age.

Next, we asked whether TE transcription was changed with aging in ovaries and testes at the level of each subfamily. We identified differentially expressed TE transcripts using our mRNA-seq analysis pipeline (FDR < 1e-6, as above; **Fig. 4A,C,D** and **Supplemental Fig. S5A-B; Supplemental Table S4A-B**). Similar to our differential gene expression analysis, we observed that most significant changes in TE expression levels occurred in middle-aged gonads (*i.e.* patterns a and b; **Fig. 4C,D** and **Supplemental Fig. S5A,B**). In the ovary, 6 TEs were downregulated only at middle-age (pattern a), and 248 TEs were specifically upregulated at middle-age (pattern b; **Fig. 4C** and **Supplemental Fig. S5A**). These age-regulated TEs seem to belong to families broadly representative of the overall turquoise killifish genomic TE content, with a predominance of LINE sequences, suggestive of general relaxation of TE control (**Fig. 4C,D**). Interestingly, this timepoint coincides with the functional enrichment for “piRNA metabolic process” for genes downregulated in middle-age ovaries (**Fig. 2C**). Thus, the transcriptional upregulation of TEs in the middle age ovary may reflect an unfavorable reactivation of certain TE species in response to the transient decline in PIWI pathway activity. Such an increase in TE transcription, even in the context of mature oocytes, is expected to be deleterious to oocyte health (Malki et al. 2014). No significant TEs were linearly downregulated or upregulated with age in the ovaries (*i.e.* patterns c and d, respectively; **Fig. 4C** and **Supplemental Fig. S5A**). In the aging testes, only 25 TEs passed our differential expression threshold (FDR < 1e-6), and they were all upregulated at middle-age (pattern b; **Fig. 4D** and **Supplemental Fig. S5B**). No significant TEs were downregulated at middle-age or linearly with age in the testes (*i.e.* patterns a, c, and d, respectively; **Fig. 4D** and **Supplemental Fig. S5B**). A burst of TE transcription in middle-age gonads could reflect peak fertility and gamete output, where a higher proportion of germ cells are undergoing epigenetic repatterning to yield increased numbers of mature gametes. Alternatively, increased TE transcription may reflect a relaxation of genome surveillance mechanisms, ultimately leading to decreased gamete quality and survivability.

### piRNAs and the PIWI pathway activity are regulated throughout life in turquoise killifish gonads

Thus far, we have found that: (i) genes encoding the key components of the PIWI pathways are robustly expressed throughout life in turquoise killifish ovaries and testes (**Supplemental Table S1C**; **Fig. 5A**), (ii) PIWI pathway genes are downregulated in ovaries at middle-age (**Fig. 5A** and **2C**), and (iii) transcription of TEs is globally regulated with gonadal aging (**Fig. 4B**), with several TE subfamilies transcriptionally specifically upregulated in middle-age ovaries and, to a lesser extent, middle-age testes (**Fig. 4C,D**).

**Figure 5.**
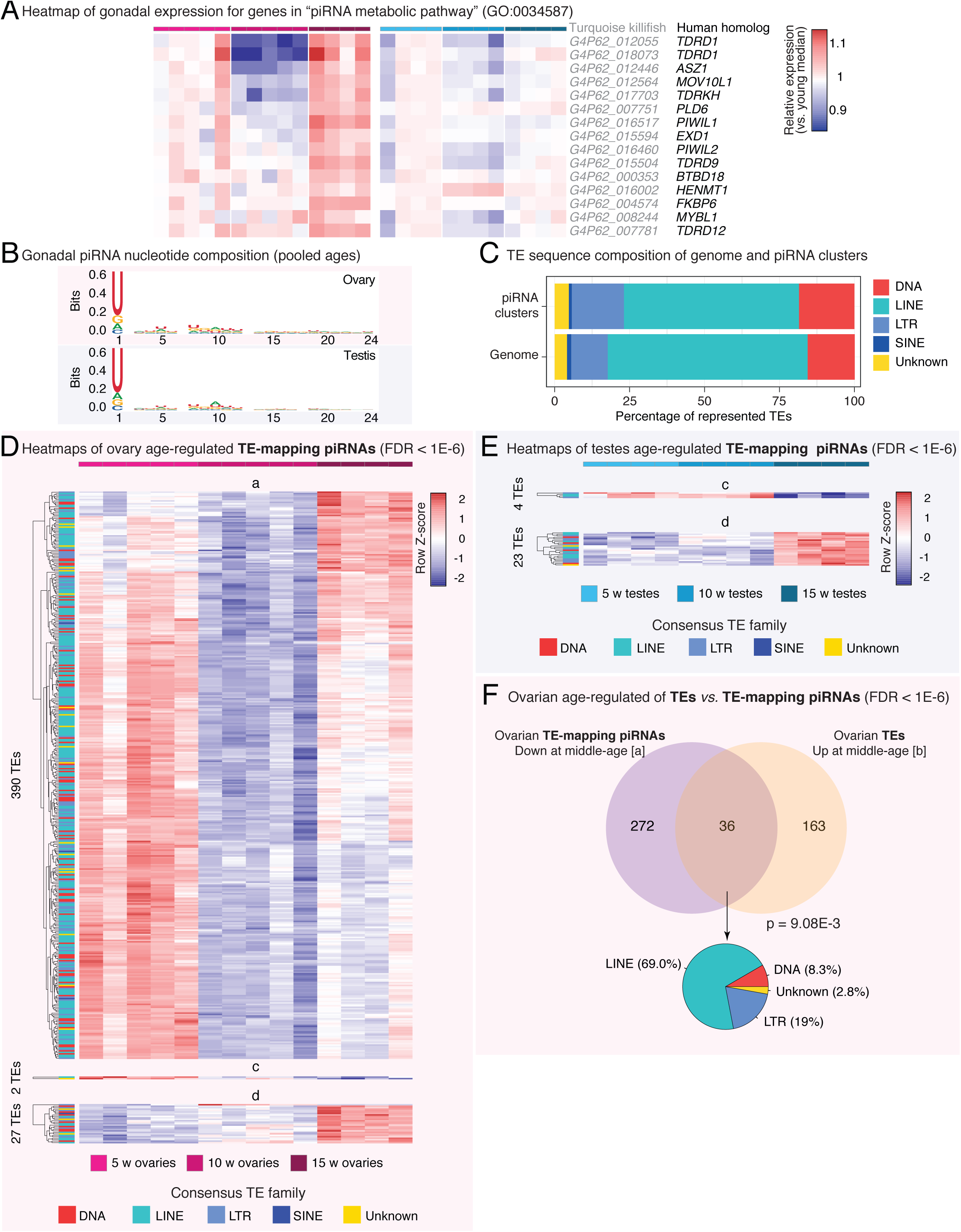
PIWI pathway activity and piRNA abundance characterization in aging turquoise killifish gonads. **(A)** Heatmaps of normalized gene expression from ovaries and testes with aging in the GO “piRNA metabolic pathway” term. Killifish gene names and their best human homologs are indicated for each row. Rows were normalized by the median expression level in the cognate young gonad, so as to facilitate visualization of age-related changes. Genes involved in this pathway tend to be significantly downregulated in middle-aged ovaries (see Fig. 2C). **(B)** Sequence logo plots of piRNA nucleotide composition from ovaries and testes, using data for each gonad type pooled across ages and replicates. As expected for piRNAs, killifish piRNAs show a strong position 1-U bias. There is also a slight bias for Adenine at position 10, consistent with active ping-pong biogenesis in killifish gonads. **(C)** Stacked barplots depicting the TE family composition of piRNA clusters and the killifish genome, as a percentage of identified elements. The piRNA cluster TE composition is broadly reflective of the TE composition of the genome. **(D)** Heatmaps of TE-targeted piRNA abundance for TEs significantly regulated with age in aging killifish ovaries by DEseq2 LRT (FDR < 1e-6). TE family is color-coded for each row. In ovaries, most differentially piRNA-targeted TEs show a decrease of piRNA counts at middle-age. **(E)** Heatmaps of TE-targeted piRNA abundance for TEs significantly regulated with age in aging killifish testes by DEseq2 LRT (FDR < 1e-6). **(F)** Venn diagram of the overlap between TE_mapping piRNAs significantly <u>up</u>regulated in middle-age ovaries (see Fig. 5D; pattern a) and TEs that are significantly <u>down</u>regulated in middle-age ovaries (see Fig. 4C; pattern b). Significance for the overlap being larger than expected by chance was calculated using Fisher’s exact test, compared to TEs detected in both analyses. A pie chart of the composition of “consistent” TEs is also reported.

Thus, we reasoned that underlying changes in piRNA abundances throughout aging may drive changes in TE transcriptional levels.

To analyze PIWI pathway activity in ovaries and testes throughout turquoise killifish life, we generated small RNA sequencing libraries from the same samples used for mRNA-seq (**Fig. 1A**, **Supplemental Fig. S1A**). We leveraged these libraries to annotate piRNA clusters and evaluate piRNA expression throughout life in the African turquoise killifish gonads. First, to computationally isolate piRNAs from our small RNA-seq library, we used a size filter of 24-35 nucleotides, which was previously shown to efficiently capture piRNAs and separate them from miRNAs (Gong et al. 2018; Huang et al. 2019) (**Supplemental Fig. S6A**). Consistent with the resulting reads deriving from piRNAs, the resultant nucleotide composition showed the expected strong 1U bias in both types of gonads (Thomson and Lin 2009) (**Fig. 5B**). In addition, these piRNA-derived reads seemed to arise from 222 unique piRNA clusters in the turquoise killifish genome, as identified by Protrac (Rosenkranz and Zischler 2012). Importantly, the predicted TE content of the identified piRNA clusters is remarkably reflective of the annotated genomic TE content (**Fig. 5C**), consistent with the co-evolution between the TE invasion of the genome and the genomic defense against TE activity. In contrast to mice, in which ovarian piRNAs are rare and the PIWI pathway is broadly believed to be non-essential (Watanabe et al. 2008), turquoise killifish ovaries seem to express the components of PIWI pathway throughout life and are replete with piRNAs. This may be the result of continuous oocyte production in the killifish ovary since PIWI is required during oogenesis (Thomson and Lin 2009; Ketting 2011; Roovers et al. 2015).

Since piRNAs will align almost exclusively to TE sequences, we modelled piRNA abundance dynamics by performing differential expression analysis on TEs using counts from the piRNAs mapping to TE sequences embedded within the genome. This approach enables us to distinguish which TE species are differentially targeted by the PIWI pathway in turquoise killifish gonads as a function of age. As piRNA biogenesis is expected to at least partially be a response to TE activity, TE-specific piRNA counts should positively correlate with expressed TE species. Indeed, we observed a strong positive correlation between the mRNA levels of TEs and the abundance of cognate piRNAs in each biological group (Spearman Rank correlation Rho ≥ 0.66; **Supplemental Fig. S6B,C**). Similar to transcriptional patterns observed in our genic and TE data, the bulk of differential piRNA expression occurred at middle-age in ovarian tissue (pattern a; **Fig. 5D** and **Supplemental Fig. S6D**; **Supplemental Table S5A**). However, mirroring the TE transcriptional changes where many TE species were <u>upregulated</u> in the middle-age ovaries (**Fig. 4C**), the majority of differentially abundant piRNAs were <u>downregulated</u> at middle-age in the ovaries (**Fig. 5D**), consistent with downregulation of the PIWI pathway genes (**Fig. 2C** and **5A**). Importantly, there was significant overlap between TEs that were specifically upregulated in middle-age ovaries and TEs targeted by piRNAs with decreased piRNA abundance in middle age ovaries (p = 9.08E-3 in Fisher’s exact test; **Fig. 5F**), consistent with the notion of a partial relaxation of piRNA-mediated TE repression in middle-age ovaries. Testis piRNA expression was not as strongly regulated with age as compared to that of ovaries, with few age-regulated piRNA species, most of which showed increased expression with age (pattern d; **Fig. 5E** and **Supplemental Fig. S6E**).

### Ping-Pong activity declines with age in oocytes, remains stable in testes

In the presence of a particular TE mRNA, the PIWI pathway initializes catalytic degradation of the TE transcript guided by TE-complementary “primary” piRNA sequences, which triggers the production of additional “secondary” piRNA sequences that specifically target the active TE sequence (Czech and Hannon 2016). This mechanism of piRNA synthesis through amplification of piRNA populations corresponding to actively transcribed TEs is known as “ping-pong” biogenesis (Czech and Hannon 2016) (**Fig. 6A**). Effectively, ping-pong biogenesis is an adaptive mechanism by which the PIWI pathway defends the germline against active TE threats (Czech and Hannon 2016). Due to the annealing requirements between primary piRNA sequences, target TE mRNAs, and TE-derived secondary piRNAs, a signature of ping-pong biogenesis is a 10 bp overlap between the 5’ ends of opposite orientation piRNAs (Wang et al. 2015) (**Fig. 6A** and **Supplemental Fig. S7A-B**). This ping-pong biogenesis mechanism can be measured globally or at the level of a specific TE sequence.

**Figure 6.**
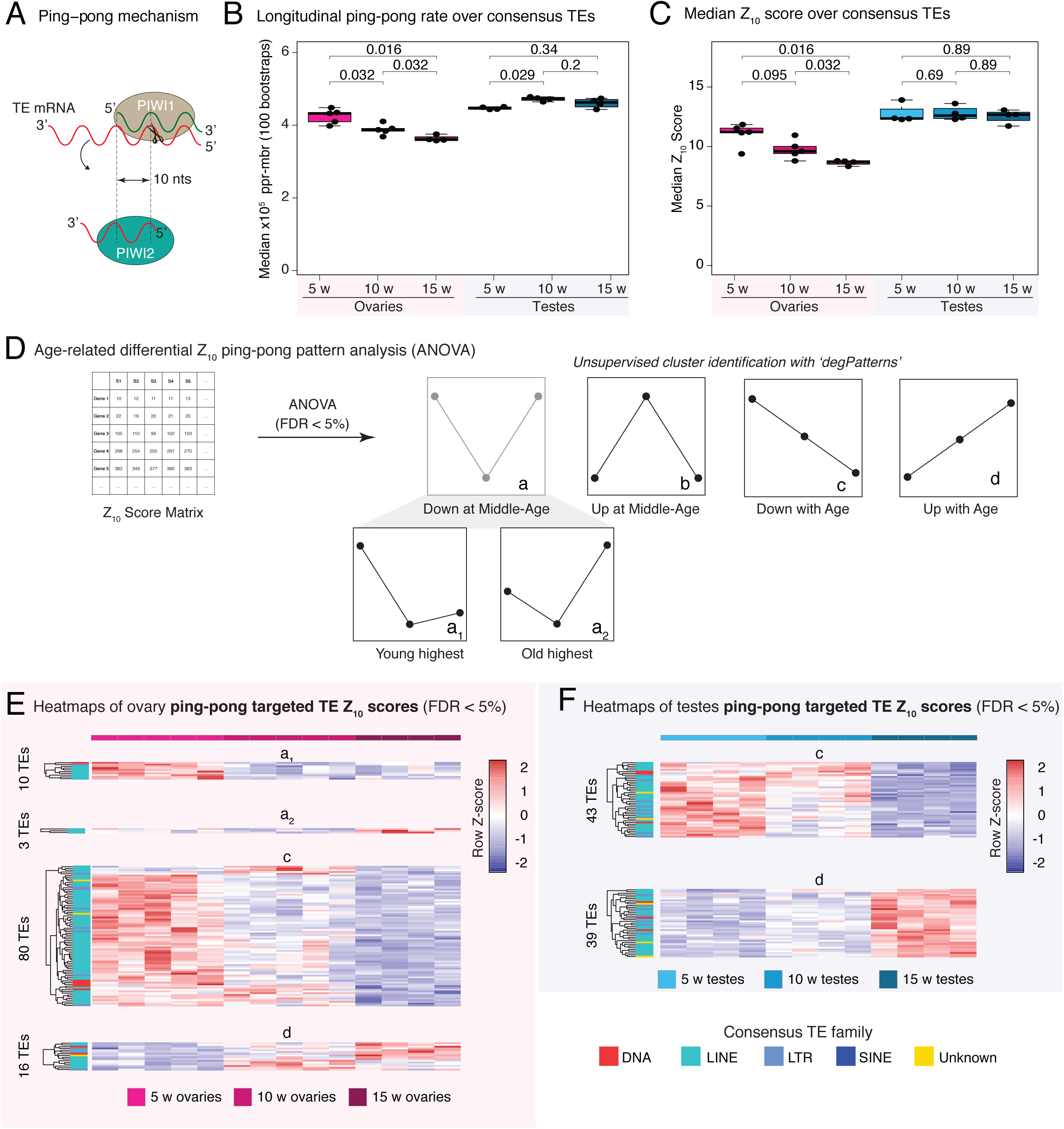
Analysis of ping-pong biogenesis in aging turquoise killifish gonads. **(A)** Explanatory diagram of ping-pong biogenesis. PIWI1 (the PIWI protein loaded with ‘primary’ piRNAs complementary to TEs) binds to and cleaves a TE mRNA species. The cleaved TE mRNA is loaded into PIWI2 (the PIWI protein loaded with cleaved TE fragments, or ‘TE-derived secondary’ piRNAs). PIWI proteins always cleave their targets 10 basepairs from the 5’ end of the piRNA. This mechanism, known as ping-pong biogenesis, will generate a preponderance of complementary piRNAs with a 10 basepair overlap, the detection of which is the basis of the assays shown in Fig. 6B**-F**. **(B)** Boxplot showing the median frequency of 10 basepair overlaps for 1 million piRNA reads over 100 bootstraps per biological sample (ppr-mbr:<u>P</u>ing-<u>P</u>ong <u>R</u>eads per <u>M</u>illion <u>B</u>ootstrapped <u>R</u>eads,) as generated by PPmeter. Age groups within each sex were compared using the non-parametric Wilcoxon rank sum test. **(C)** Boxplot showing median Z_10_ scores over consensus TEs for each biological sample. Z10 scores are an alternative measure of ping-pong. Non-parametric Wilcoxon Rank Sum tests were used to test for significant differences between age groups. **(D)** Schematic for analysis of differential Z_10_ score patterns with gonadal aging. Z_10_ scores were generated for each consensus TE sequence in FishTEDB in each biological sample. An ANOVA test was run to detect any significant differences in Z_10_ score between age groups within each sex (FDR < 5%). Significant TEs were assigned to groups based on expression dynamics that align to the broad groups defined in Fig. 2A, with the detection of 2 new subgroups showing minimal expression at middle-age (a_1_ and a_2_), with highest Z_10_ scores in the young samples and old samples, respectively. **(E)** Heatmaps of differential Z_10_ scores with age in aging killifish ovaries by ANOVA (FDR < 5%). **(F)** Heatmaps of differential Z_10_ scores with age in aging killifish testes by ANOVA (FDR < 5%). A roughly equivalent amount of consensus TE sequences show linear up- and downregulation with age. The cognate TE family is color-coded by row, and does not reveal any specific bias.

We first used the PPmeter tool from the Protrac software suite to globally measure ping-pong biogenesis rates in each biological group (Jehn et al. 2018) (**Fig. 6B**). This tool calculates the instances of 10 bp overlaps out of every million overlap pairs assayed to report the relative fraction of reads engaged in active ping-pong within a given piRNA sequencing library. We used this tool for each biological replicate with 100 bootstraps and report the median values in each biological sample over bootstrap samples (**Fig. 6B**). In parallel, we also used an independent method in which, for each consensus TE sequence, we computed Z_10_ scores, a standard measure of ping-pong activity defined as the Z-score of the 10 bp overlap frequencies between opposite orientation piRNAs within a 20bp window (Han et al. 2015; Vandewege et al. 2022). For each replicate, we took the median Z_10_ score over all detected TE sequences to assess ping-pong at a global scale (**Fig. 6C**). Convincingly, both orthogonal methods revealed a steady decrease of the piRNA population participating in ping-pong biogenesis with aging in the ovaries, whereas this fraction remained relatively stable in the testes (**Fig. 6C**). The age-related decline in ovarian ping-pong activity may be related to a decrease in piRNA processing component expression starting at middle age (**Fig. 5A**). This trend could result in a decrease in the ability of the PIWI pathway to control TE transcription levels with ovarian aging. Alternatively, this steady decrease with aging in ovarian ping-pong biogenesis may reflect the interplay of (i) a decreased efficiency of the PIWI pathway at middle-age (**Fig. 2C**, **5A**), and (ii) an overall decrease of TE transcription levels in old ovaries (**Fig. 4B**). Meanwhile, the observed steady global ping-pong rates in testes are consistent with relatively stable TE and piRNA expression dynamics relative to ovaries that we observed (**Fig. 4D, 5E**).

We were curious whether ping-pong biogenesis may be differentially regulated for each consensus TE sequence. For this purpose, we further examined the consensus TE Z_10_ data at the consensus TE transcript level. We used an ANOVA-based analysis to determine which TEs have differential levels of ping-pong activity as measured by Z_10_ scores with aging in ovaries and testes (**Fig. 6D-F** and **Supplemental Fig. S7C-D**). Then, TEs with significant age-related changes in ping-pong biogenesis were clustered into patterns, similar as before using the ‘degPatterns’ paradigm (**Fig. 2A**, **6D**). This analysis identified TEs with ping-pong biogenesis levels corresponding to previously described patterns b, c and d. In addition, it identified 2 significant patterns for TEs with decreased ping-pong biogenesis at middle-age (the pattern previously designated as (a)): (a_1_) with maximal ping-pong biogenesis in the young gonad, or (a2) maximal ping-pong biogenesis in the old gonad. Interestingly, most of the TEs showing differential ping-pong levels in the ovaries showed significantly decreased ping-pong with aging (pattern c; **Fig. 6E** and **Supplemental Fig. S7C**), consistent with the global ping-pong trends (**Fig. 6B,C**). In the testes, 82 TEs show differential ping-pong scores but without a clear global age-related trend, possibly reflecting a more robust PIWI activity throughout life (**Fig. 6F** and **Supplemental Fig. S7D**). In summary, multiple methods of analysis show ping-pong activity progressively wanes in the aging ovary, but remains steady in the testes which may be reflective of respective gametogenesis rates in the female versus male gonads throughout life.

## Discussion

### A resource for the study of reproductive aging in vertebrates

The germline is considered an immortal cell lineage but the somatic cells that comprise and support the germline are affected by aging. In this study, we examined how aging affects the female and male gonads of a naturally short-lived vertebrate species, the African turquoise killifish. To our knowledge, this dataset is the largest ‘omic’ dataset evaluating vertebrate ovarian and testicular aging and will represent a unique resource for reproductive aging research. To note, we performed this study using only fish from the inbred GRZ laboratory strain, a single genetic background of African turquoise killifish, continuously inbred since originally established in 1973. Thus, it is possible that other strains of this species may show different gonadal aging trajectories. For instance, other laboratory strains (*e.g.* MZM-0703, MZM-0410) are longer lived than the GRZ strain (Kim et al. 2016), and could thus exhibit differences in their reproductive aging patterns. Further, laboratory strains represent only a fraction of the genetic diversity available in the wild for this species. Indeed, GRZ and wild-derived turquoise killifish reared under the same laboratory conditions experience different fecundity and fertility rates during aging (Zak and Reichard 2021), although the same overall reproductive aging trends (albeit at different timescales) were observed across populations. Finally, African turquoise killifish may not experience significant age-related decline in reproductive function over their short lifespan in the wild, although a decline in relative fecundity (*i.e.* egg generation controlled for total body mass) has been reported in wild females (Vrtilek et al. 2018a). Thus, we note that future studies incorporating more genetically diverse strains (and in their natural habitat) may reveal variation in the patterns of molecular regulation of reproductive aging in this species.

With our inclusion of a middle-aged time point, in addition to young adult and old ages, this dataset enables the discovery of dynamic and non-linear patterns of regulation, thus helping understand gonadal aging trajectories in addition to just the endpoint of gonadal aging. Indeed, we found that many key events, including those relating to TE transcriptional regulation and piRNA pathway regulation, are most salient at middle-age in ovaries, rather than undergoing linear changes with age. Our observations differ from reports in *Drosophila* egg chambers, in which PIWI-related genes were upregulated during aging though global TE expression levels remained largely stable (Erwin and Blumenstiel 2019). Importantly, our observations that the aging ovarian transcriptome is more impacted than the testicular transcriptome are consistent with previous microarray studies of mouse gonadal aging (Sharov et al. 2008). To note, in long-lived species (*e.g.* humans), male germ cells can accumulate mutations from repeated divisions during aging, which can impact offspring fitness and disease risk – the “father’s age effect” (Kong et al. 2012). Due to the relatively short lifespan of vertebrate models used in aging research (*i.e.* African turquoise killifish, mouse), it is unlikely that the impact of repeated male germ cell divisions over male lifespan can be appropriately modeled in these species, thus potentially explaining the relative paucity of changes observed in the molecular landscapes of aging testes.

### A unique crosstalk between piRNA biogenesis and TE activity in the aging vertebrate gonad

We characterized TE expression and PIWI pathway behavior throughout life in turquoise killifish ovaries and testes. Importantly, we found that the PIWI pathway controls TE expression in the gonads throughout the turquoise killifish lifespan. Age-related fertility decline is common across many metazoans (Jones et al. 2014). The turquoise killifish is no exception to this rule, with fertility generally declining after middle-age, especially in females (Zak and Reichard 2021). For both sexes, fertility peaks around the middle-age timepoint used in this study (Zak and Reichard 2021). Interestingly, this timepoint coincided with the largest changes in TE- and piRNA related events in both sexes, albeit much more pronounced in ovaries compared to testes. This may correspond to a period of increasing gametogenesis in which epigenetic modifications are globally removed from germ cells prior to repatterning. Such epigenetic erasure would provide a means of “escape” for TEs in the form of transcription. Since the females are asynchronous spawners and continuously produce oocytes, we might expect that, like in males, TE transcription may reflect gamete production output. To note, our observations may capture differential amplitudes of reproductive aging between the sexes - with more dramatic increases in TE transcription in ovaries, which undergo a clearly measurable reproductive senescence, compared to more modest changes in testes, which are thought to undergo near negligible reproductive senescence (Vrtilek et al. 2018a; Zak and Reichard 2021).

Overall, even in this short-lived species, the PIWI pathway seems to broadly keep gonadal TE transcription stable at the global transcriptomic level (**Fig. 4B**), although there is clear regulatory variation depending on specific TE sequences. This relatively controlled transcriptional output for TE loci seems to contrast with reports of global transcriptional derepression of TE sequences in aging somatic tissues (De Cecco et al. 2013; Bravo et al. 2020). When expressed, TEs can cause DNA damage and/or initiate innate immune responses due to their intrinsic similarity to viruses (Simon et al. 2019). This contrast between TE activity in germline/somatic tissues is especially interesting considering that non-aging taxa (*e.g. Hydra*, *Planaria*) can express PIWI in somatic stem cells (Schaible et al. 2015; Sahu et al. 2017; Teefy et al. 2020). This suggests that looser TE control may drive aspects of aging, especially in the soma, whereas tighter TE control is crucial to allow the emergence of non-aging states, such as those found in the germline.

### Dynamic TE control in the aging African turquoise killifish ovary

If piRNA metabolism dynamics in the turquoise killifish ovary are similar to those of zebrafish (with *PIWIL1* expression being largely confined to immature germ cells) (Liu et al. 2022), then downregulation of PIWI-related genes in bulk middle-age ovaries could merely reflect a lower underlying proportion of immature oocytes. However, our histological and deconvolution analyses suggest that large scale changes in oocytes maturation are unlikely to underlie observed changes in PIWI pathway activity during ovarian aging (**Supplemental Fig. S4**).

Alternatively, increased TE expression in middle-aged ovaries (and thus decreased TE control) may be a symptom of decreased oocyte quality before fertility starts to decline, suggesting that reproductive aging may precede organismal decline. Specifically, decreased TE transcription and increased expression of PIWI pathway components in old ovaries may constitute a delayed response to increased TE transcription and decreased expression of PIWI pathway components at middle-age.

Finally, it is possible that such a transient relaxation of TE control in middle-age ovaries for the turquoise killifish may have adaptative value. Indeed, parental age in the turquoise killifish has been proposed to convey environmental information for evolutionary benefit. Embryos of old breeders enter are more likely to state of suspended animation known as diapause, which killifish utilize to persist under dry conditions between rainy seasons, than the embryos of young breeders (Api et al. 2018). Within a single wet season, the offspring of older mothers are more likely to immediately encounter a dry season, while those of younger mothers are more likely to propagate and yield multiple generations (Api et al. 2018).

Likewise, allowing some level of age-associated TE mobilization in the germline through tunable activity of the PIWI pathway could help ensure adequate levels of genetic diversity in a species with naturally small populations due to its unique ecology. Consistently, annual killifish, like *Nothobranchius furzeri*, have much higher genomic TE content than non-annual killifish species that do not experience such extreme population bottlenecks from seasonal aquatic habitat desiccation (Cui et al. 2020). TEs have long been appreciated in plant biology for their contribution to genetic diversity and are now appreciated sources of genetic diversity and drivers of speciation in animals (McClintock 1984; Gonzalez et al. 2010; Akagi et al. 2013; Belyayev 2014; Bariah et al. 2020; Chen et al. 2020; Catlin and Josephs 2022). In line with the hypothesis that relaxed germline TE restriction may enhance population-level genetic diversity is our observation that ping-pong rate in killifish ovaries decreases steadily with age. This age-related PIWI behavior could support a population facing severe bottlenecks by first generating many healthier (but less genetically diverse) progeny during young adulthood. Subsequently, with females still breeding at a more advanced age, reduced PIWI pathway activity could help generate a more genetically diverse pool of progeny that could improve genetic diversity for the long-term adaptability of the local population.

In summary, our data supports the notion that even a short-lived species such as the African turquoise killifish can maintain adequate TE silencing in germline-containing tissues throughout life, likely due to the activity of the PIWI pathway. The impact of fluctuations in TE transcription and PIWI pathway activity, especially in ovarian tissues, remains an open question. Understanding disruptions to gonadal genomic regulatory networks during aging will be crucial to define strategies to preserve or prolong reproductive fitness with aging.

## Material and methods

### African turquoise killifish husbandry

African turquoise killifish were raised according to gold standard procedures (Dodzian et al. 2018). To decrease risks of aggression, adult fish were single housed in 2.8L tanks on a recirculating aquatics system manufactured by Aquaneering Inc. System water parameters were as follows: temperature: 29°C; pH: 7.3; conductance: 670-750 S; Μ ammonia and nitrite: 0ppm and Nitrate: ∼15 ppm. Adult fish were fed twice per day with Hikari Freeze Dried Bloodworms 4-6 hours apart, and live *Artemia* once per day. Fry were reared in system water incubated at 28°C, then placed on the recirculating system starting at 2 weeks post hatch. Fry were fed live *Artemia* exclusively until 4 weeks post hatch. The fish facility was kept on a light/dark cycle of 13/11 hours (lights on 9am-10pm). The fish were euthanized by immersion in 1.5 g/L of Tricaine MS-222 dissolved in system water followed by decapitation. All fish were euthanized between 2 and 4 pm to minimize circadian effects.

All husbandry conditions and experimental procedures were approved by the University of Southern California (USC) IACUC. Animal care and animal experimentation were performed in accordance with IACUC approved protocols for *Nothobranchius furzeri at* the University of Southern California (approved protocols *20879* and *21023*).

### Lifespan data collection

Lifespan data was derived from historical data in our colony collected 2019-2022. Time from hatching to humane endpoint/euthanasia or natural death was recorded as killifish lifespan. All lifespan data reported here is derived from single-housed fish to limit death or injuries linked with frequent fighting in this species.

### RNA extraction and sequencing

Ovaries and testes were dissected from 5-week-old, 10-week-old, and 15-week-old turquoise killifish (N = 5 per group) and flash-frozen on dry ice. Our chosen number of biological replicates for each group is in the range shown to be robust for differential gene expression analysis by RNA-seq (Lamarre et al. 2018). Importantly, we defined the 3 age groups based on generally accepted guidelines in the field of aging research: (i) the chosen “young” adult time point at 5 weeks represents sexual maturity (*i.e.* the age at which individuals are able to reproduce), to avoid measuring developmental signals, (ii) the chosen “old” time point at 15 weeks represents ∼90% population survival in our GRZ colony regardless of sex (**Fig. 1B**), allowing sample of aged animals before any risk of survivorship bias (Flurkey et al. 2007), and (iii) the chosen “middle-aged” time point at 10 weeks is the midpoint between these extremes. Total RNA was extracted from flash-frozen gonads by homogenizing using Trizol (Thermo Fisher, 15596018) and Lysing Matrix D tubes (MP Biomedicals, 11422420) on a BeadBug 6 microtube homogenizer (Millipore Sigma, Z742683) for 2 rounds of 30 seconds at 3500 rpms. RNA was assayed for quality at the USC Genome Core using the Agilent Bioanalyzer total RNA assay, and only high-quality RNA samples (RIN > 7) were selected for further processing to avoid RNA degradation related biases. Due to poor RNA quality, samples for three males (one from each age group) and one 15-week female were eliminated at this stage. RNA samples were sent to Novogene Corporation (USA) for library construction and sequencing, where mRNA and small RNA sequencing libraries were generated from the same total RNA samples in parallel. mRNA libraries were prepared by enrichment with oligo(dT) beads and sequenced on an Illumina NovaSeq 6000 generating 150 bp paired-end libraries. Small RNA seq libraries were generated by 3’ adapter ligation, 5’ adapter ligation, reverse transcription, PCR amplification, and gel purification and size selection. Small RNA samples were sequenced on an Illumina Novaseq 6000 generating 50 nucleotide single-end read libraries. Raw sequencing data has been deposited to SRA under PRJNA854614.

### mRNA-seq data pre-processing and genomic alignment

FASTQ reads from mRNA libraries were first hard-trimmed using fastx_trimmer (fastx_toolkit 0.0.13) (Gordon A. 2010), to remove 5’ adapter fragments and low quality bases at the 3’ end of the reads using parameters “-f 16”, “-l 100”, and “-Q33”. Next, Illumina adapters were removed using TrimGalore 0.6.7 (cutadapt 3.3) (Felix Krueger 2021). To maximize our capture of relevant reads, we used the most contiguous genome reference currently available for the African turquoise killifish, Genbank accession GCA_014300015.1, described in Williamsen *et al.,* 2020 (Willemsen et al. 2020), as well as its corresponding gene annotation file. Since analysis of repetitive sequences was desired, we performed soft-masking with Repeatmasker 4.1.2-p1 (Smit 2013-2015) and a turquoise killifish specific consensus TE library downloaded from FishTEDB (downloaded: February 5, 2021) (Shao et al. 2018). A TE-specific GTF file for downstream analysis was generated using a custom R script for compatibility with the TETranscripts counting program (Jin et al. 2015). RNA-seq reads were aligned to the soft-masked turquoise killifish genome using STAR 2.7.0e, allowing 200 multimappers to accommodate TE sequence alignment (Dobin et al. 2013; Jin et al. 2015). Gene and TE counts were generated using TETranscripts 2.2.1 with default settings using gene annotations from GenBank and the RepeatMasker derived custom TE-specific GTF file.

### Differential gene and TE expression analysis

The TETranscripts count matrices for genes and TEs were analyzed together in R 4.1.2 using DESeq2 v1.34.0 (Love et al. 2014). To identify differentially expressed transcripts and their patterns with aging in ovaries and testes, we used likelihood ratio testing [LRT] separately for each sex with each age as its own group. Significant genes were defined as those with an adjusted p-value of < 1E-6 with the DESeq2 LRT test. Genes and TEs were then analyzed separately. Once significantly differentially expressed genes and TEs were identified, clusters of differentially expressed genes and TEs were unbiasedly identified using the ‘degPatterns’ function from the ‘DEGreport’ v1.30.3 package (Pantano 2022).

### Fractional TE analysis

To compare fractional TE expression in killifish gonads, the TEtranscripts count matrices used for differential gene and TE expression analysis were imported into R. For each biological replicate, assigned counts were classified as genic or TE-derived and the fractional TE representation for each replicate library was calculated as TE read counts divided by total read counts. Fractional counts were grouped by age and sex and tested for significance by the non-parametric Wilcoxon rank sum test between groups within each sex.

### Functional enrichment analysis

Data was prepared for functional enrichment analysis by running blastp (ncbi-blast v2.13.0) with African turquoise killifish protein sequences from the (Willemsen, 2020) GCA_014300015.1 genome version, against Ensembl release 104 human protein sequences. For each query killifish sequence, only top hits with E-value < 1e-3 were retained, and these human homologs were used for functional enrichment analyses. Gene ontology (GO) enrichment analysis was performed using ‘GOstats’ 2.60.0 with GO terms downloaded using ‘biomaRt’ 2.50.3 (Durinck et al. 2009) corresponding to Ensembl release 104. GO analysis was performed using the “Biological Process”, “Cellular Component”, and “Molecular Function” categories with a significance threshold equal to a false discovery rate [FDR] < 5%. GO analysis was performed on each of the gene patterns identified in the LRT-based DGE analysis described above.

### piRNA selection, annotation, and piRNA cluster analyses

FASTQ reads from small RNA libraries were trimmed of their adapters while keeping a minial length of at least 18 nucleotides using TrimGalore 0.6.7 (cutadapt 3.3) (Felix Krueger 2021) with the commands “--length 18”, “-a GTTCAGAGTTCTACAGTCCGACGATC”, and “-a GTTCAGAGTTCTACAGTCCGACGATC”. piRNAs were selected by size-selecting sequences between 24-35 nucleotides in length from the trimmed small RNA libraries (Gong et al. 2018; Huang et al. 2019). piRNA length distributions (**Supplemental Fig. S6A**) were generated by combining piRNA length histogram data by sex and plotting the resultant length distribution in R. piRNA nucleotide distribution for each sex was computed by converting piRNA reads to fasta format files and concatenating by sex. Position weight matrices were generated from these fasta files in R using the package ‘Biostrings’ 2.58.0 (**Fig. 5B**). Sequence logos were generated from position weight matrices using the R package ‘ggseqlogo’ 0.1.

To identify piRNA clusters, we used the Protrac 2.42 software suite (Rosenkranz and Zischler 2012). Briefly, piRNAs were collapsed to unique reads, while retaining count numbers using the functions TBr2_collapse.pl and TBr2_duster.pl. These reads were then mapped to the soft-masked reference genome described previously using the function sRNAmapper.pl and clusters were identified by a preponderance of piRNA mapping within a particular window using the function proTRAC_2.4.3.pl. Clusters were identified for each biological replicate before being combined such that only unique piRNA clusters remained using the Protrac function merge.pl. To assess the TE composition of piRNA clusters relative to the genome, the intersectbed command (bedtools 2.27.1) was used to extract piRNA cluster fasta sequences from the genome (Quinlan and Hall 2010). Then, Repeatmasker 4.1.2-p1 was run as above on the isolated piRNA cluster fasta sequences to obtain TE family information, which was then contrasted with the genome-wide data in R.

### piRNA differential expression and Ping-Pong analysis

piRNAs were mapped to the soft-masked genome reference using bowtie 1.2.3 (Langmead et al. 2009) with three allowed mismatches since piRNAs are known to bind with imperfect base-pairing (Juliano et al. 2014; George et al. 2015; Zhang et al. 2015; Teefy et al. 2020) using the parameters “-v 3 -a --best --strata -S” before being converted to bam format files with samtools 1.10 (Danecek et al. 2021). piRNA counts were then assigned using ‘featureCounts’ from the Subread 2.0.2 package, using fractional count attribution for multimapping reads (Liao et al. 2014). For featureCounts based read counting, a custom GTF file consisting of genes, TEs, piRNA clusters, and rRNA repeats as features was used to account for piRNA mapping to all relevant features to minimize biases in piRNA counts.

The resultant count matrix was imported into R, and rRNA-mapping reads were discarded prior to normalization. Then piRNA counts that mapped to genes, TEs, and piRNA clusters were normalized together using DEseq2. LRT-based clustering and DGE analysis was then performed for piRNA counts that mapped to TE sequences as described above.

To measure ping-pong biogenesis, we only considered piRNAs mapping to consensus TE sequences, since they constitute the most biologically relevant target of gonadal piRNAs. To analyze ping-pong biogenesis at a global level, we used the PPmeter 0.4 program (Jehn et al. 2018). Briefly, PPmeter reports the rate of 10 bp overlaps between piRNA reads, a proxy for ping-pong biogenesis, by generating pseudo-replicates through boostrapping one million piRNA reads (100 bootstraps). Using this approach, piRNA libraries can be directly compared to each other by measuring the number of ping-pong events (*i.e.* 10 bp overlaps) per one million piRNA reads. The resultant metric, ping-pong rate per million bootstrapped reads (ppr-mbr) was calculated for each bootstrap in each piRNA library. Bootstrap values were combined per biological replicate and the median value was taken. Median ppr-mbr values were then grouped by age and sex and tested for statistical difference by age group within each sex by the non-parametric Wilcoxon rank sum test.

For TE-specific level ping-pong measurement, we used custom scripts that calculated Z_10_ scores for each consensus TE sequence. Z_10_ scores, as defined in (Han et al. 2015; Vandewege et al. 2022) is the difference of 10 bp overlap occurrences and the mean of all other overlap occurrences within a 20 bp window divided by the standard deviation of all other overlap occurrences. This is shown in the following equation:

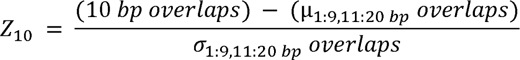

To generate Z_10_ scores, the distances between piRNA 5’ ends that mapped to opposite strands of consensus TE sequences were tabulated into a histogram using a custom bash script. The resultant histograms were imported into R and processed in a custom script that reported the Z_10_ score for each consensus TE sequence. For some TE sequences, coverage was incomplete such that we could not derive Z_10_-score information; these sequences were discarded from consideration. To measure the changes in Z_10_ scores with age, TE Z_10_ scores were separated by sex and grouped by age. Group-level Z_10_ scores were tested for significant differences by 1-way ANOVA in R, and only TEs with an adjusted p-value <0.05 were considered significantly regulated with aging in either gonad type. Significant TEs were then classified into clusters using the ‘degPatterns’ function from ‘DEGreport’ 1.30.3 package as described previously. For biological group-level analysis of Z_10_ scores, median Z_10_ values over all detected TE sequences were taken for each biological replicate and plotted as a function of age and sex. Difference in Z_10_ scores per group were tested between age groups within sexes by a non-parametric Wilcoxon rank sum test.

### Scripts and code availability

All scripts used in this study are available as an archive accompanying this manuscript (Supplemental Code), and on the Benayoun laboratory Github (https://github.com/BenayounLaboratory/Killifish_reproductive_aging_resource). All R scripts were run using R version 4.1.2.

## Data access

All raw sequencing data generated in this study have been submitted to the NCBI BioProject database (https://www.ncbi.nlm.nih.gov/bioproject/) under accession number PRJNA854614.

## Acknowledgments

Some panels were made with BioRender.com. We thank Suchi Patel of the USC Genome Core for running RNA bioanalyzer assays. We thank Jomille Jerez, Isabel Ollerton, Rajyk Bhala and Katelyn Hsu for assistance with killifish husbandry. We thank Dr. Carolyn Phillips, Dr. Minhoo Kim, Dr. Itamar Harel, Justin Gilmore, Juan Bravo, Casandra McGill, and Rajyk Bhala for insights and feedback on our manuscript. We thank Drs. Ryo Sanabria and Gilberto Garcia for advice on histological image analysis using ImageJ. This work was supported by an NIA T32 AG052374 Postdoctoral Training Grant fellowship to B.B.T, NIA R21 AG063739, NIGMS R35 GM142395, a pilot grant from the NAVIGAGE Foundation, and a Hanson-Thorell Family award to B.A.B.

The authors acknowledge the Center for Advanced Research Computing (CARC) at the University of Southern California for providing computing resources that have contributed to the research results reported within this publication (https://carc.usc.edu). Ovarian histological analysis was performed by the Translational Pathology Core at the USC Norris Comprehensive Cancer Center (supported by NCI P30 CA014089).

## Author contributions

B.B.T. and B.A.B. designed the study. B.B.T. and A.A. performed fish husbandry. A.A. dissected killifish gonadal tissues. B.B.T. performed RNA extraction of samples and computational analyses, with assistance from A.X., P.P.S. and B.A.B. B.B.T. and B.A.B. wrote the manuscript with input from all authors. K.H. captured microscope images of aging ovaries, B.B.T., A.X., K.H. and B.A.B. conducted ImageJ analysis of ovarian histology. All authors edited and commented on the manuscript.

