## Supplemental methods and supplemental figure legends for "Dynamic regulation of gonadal transposon control across the lifespan of the naturally short-lived African turquoise killifish"

**Supplementary Methods**

*Principal Component Analysis [PCA], Hierarchical Clustering and Variance Partition Analysis*

The TETranscripts count matrices for genes and TEs were analyzed together in R 4.1.2 using DESeq2 v1.34.0 (Love et al. 2014). For global analyses of transcriptional aging patterns across testes and ovaries, all samples were analyzed and normalized together, using sex and age as model covariates (**Fig. 1D-F**; **Supplemental Fig. S1B-S**). The DEseq2 VST-normalized expression log2 count matrices were used as input for analyses of sample relationships: (i) Principal Component Analysis (PCA), (ii) hierarchical clustering with bootstrap resampling with ‘pvclust’, and (iii) analysis of explained variance as a function of age using ‘variancePartition’. For hierarchical clustering, we used ‘pvclust’ 2.2-0 (Suzuki and Shimodaira 2006), using correlation as a distance metric (*i.e.* ‘cor’), average linkage and 1000 bootstrap samples. For the analysis of explained variance by age in ovaries and testes, we leveraged a linear mixed model approach optimized for gene expression studies, as implemented in R package ‘variancePartition’ 1.24.1 (Hoffman and Schadt 2016).

These analyses for piRNA expression were performed similarly, using the count matrix obtained as described in the methods.

*Histological analysis of aging ovarian tissue*

Ovaries were dissected from 5-week-old, 10-week-old, and 15-week-old female turquoise killifish (N = 2-3 per biological group). Ovaries were fixed in Bouin's solution (Sigma HT10132) at room temperature for 24 h, then washed off using 70% ethanol. Fixed tissue was processed with the Translational Pathology Core at the University of Southern California Norris Comprehensive Care Center for paraffin embedding, sectioning and histological stains. Samples were sectioned longitudinally and for each sample, slides were prepared and stained with hematoxylin and eosin (H&E). Each slide contained 2 tissue sections per slide for technical robustness. Samples were imaged on a Keyence BZ-X710 microscope at 4X magnification.

One section/microphotograph per animal was used to characterize oocyte diameter by four blinded observers using ImageJ/FIJI 2.9.0/1.53t. The distribution of Oocyte diameters at each age group, for each independent observer, is reported in **Supplemental Fig. S4B**, as well as comparison by consecutive age groups (using a goodness-of-fit Kolmogorov-Smirnov test to compare empirical distributions), suggesting similar distribution of oocyte maturation stages irrespective of age.

Although this is a small cohort of fish (N = 8), we believe that large differences in oocyte maturation patterns (like those required to observe dramatic changes at the transcriptional level of bulk ovaries) should still be detectable even with these small numbers. Quantified histology pictures have been submitted to Figshare (doi:10.6084/m9.figshare.21572727).

*Deconvolution analysis of bulk African turquoise killifish ovarian transcriptomes*

To determine whether there were large changes in the cellular composition of killifish ovaries during aging which may underlie transcriptional changes in components of the PIWI pathway, we leveraged a recent single-cell RNA-seq dataset generated on ovaries from young adult zebrafish (Liu et al. 2022). To maximize our ability to obtain accurate deconvolution results, we used the R ‘granulator’ 1.2.0 pipeline (Pfister et al. 2021), which allows benchmarking and use of 7 state-of-the-art transcriptome deconvolution methods.

To perform the deconvolution process, granulator requires an input matrix of reference gene expression signatures on pure cell types to estimate the relative proportions of each cell type of interest in a target bulk dataset. Thus, we took advantage of the annotations of the single cell RNA-seq zebrafish ovarian dataset that had been deposited to the Broad Institute Single Cell Portal (accession SCP928). Importantly, best deconvolution results are obtained from cell types that are well defined (Pfister et al. 2021), so we collapsed cell types with annotated subsets to their parent cell types (*i.e.* meiotic oocytes, mature oocytes, follicular cells). UMI counts were collapsed by cell type to generate signature expression profiles. In addition, to test deconvolution accuracy process and benchmark deconvolution methods, we also generated five ovarian *in silico* pseudobulk samples with known proportions of each cell type, randomized from the “true” observed proportions in the zebrafish ovarian single cell dataset.

Granulator requires “transcript per million” (tpm) normalized input RNA-seq data. Thus, for the turquoise killifish input matrix, tpm were obtained by normalizing gene counts from TEtranscripts (see above) relative to transcript lengths, since mRNA-seq covers the entire length of transcripts. For the single cell data, the 10xGenomics protocol is based on sequencing only the sequence upstream polyA stretches of transcripts, thus typically covering only ~100bp of the 3’UTR of transcripts. Thus, UMIs were normalized relative to a 100bp length. To “translate” gene expression profiles from the zebrafish context to the turquoise killifish context, we used sequence homology with blastp (ncbi-blast 2.10.0+) with African turquoise killifish protein sequences from the (Willemsen, 2020) GCA_014300015.1 genome version, against Ensembl release 108 zebrafish protein sequences. For each query zebrafish sequence, only the top hit with E-value < 1e-5 was retained as its turquoise killifish homolog for deconvolution analysis.

To assess the accuracy of our analyses, we observed that germ cell expression signature profiles clearly cluster apart from that of somatic cell types (**Supplemental Fig. S4C**). Importantly, expression of PIWIL1 is mostly observed in the germline compartment (*i.e.* germline stem and progenitor cells [GSPCs], meiotic oocytes, and maturing oocytes), with maximal expression in more immature germ cells (*i.e.* GSPCs and meiotic oocytes; **Supplemental Fig. S4D**), consistent with the analysis reported in Liu et al., 2022. We benchmarked deconvolution algorithms with ‘granulator’ to determine whether they were sensitive enough to accurately detect cell types within *in silico* pseudobulk ovarian transcriptional profiles. All benchmarked algorithms performed well (mean Pearson correlation coefficient metric of known *vs.* predicted ovarian cell proportions > 0.88), with support vector regression [SVR] and non-negative least squares [NNLS] as the top 2 performing algorithms (**Supplemental Fig. S4E**).

After verifying performance on ovarian *in silico* pseudobulk mixtures, we applied the signature matrix together with SVR and NNLS to the turquoise killifish ovarian transcriptome. Importantly, we did not detect substantial changes in the proportion of immature germ cells between young and middle-aged ovaries by SVR nor NNLS (*i.e.* GSPCs and meiotic cells, which typically show highest expression of PIWI-pathway component genes in young ovaries; **Supplemental Fig. S4D**). Changes in immature germ cells proportion were detected between middle-aged and old ovaries, albeit in opposite directions, and thus not supporting a loss or gain of these cells (**Supplemental Fig. S4F-G**). Thus, our deconvolution analysis suggests that PIWI pathway transcriptional downregulation at middle-age is not driven by broad changes in the ovarian content of immature germ cells.

***Legends to Supplemental Figures:***

**Supplemental Figure S1. Global characterization of aging turquoise killifish gonadal dataset.**

**(A)**  Description of datasets generated in this study. A total of 26 RNA-seq and 26 small RNA-seq datasets were generated from ovary and testes from ages 5, 10, and 15 weeks old. **(B-G)** PCA plots of (B) ovarian gene expression, (C) ovarian TE expression, (D) ovarian TE-targeted piRNA abundance, (E) testicular gene expression, (F) testicular TE expression, (G) testicular TE-targeted piRNA abundance. **(H-M)** pvclust hierarchical clustering of (H) ovarian gene expression, (I) ovarian TE expression, (J) ovarian TE-targeted piRNA abundance, (K) testicular gene expression, (L) testicular TE expression, (M) testicular TE-targeted piRNA abundance. **(N-S)** VariancePartition analysis of variation by age (vs. residual variance) of (N) ovarian gene expression, (O) ovarian TE expression, (P) ovarian TE-targeted piRNA abundance, (Q) testicular gene expression, (R) testicular TE expression, (S) testicular TE-targeted piRNA abundance.

**Supplemental Figure S2. Differential gene expression analysis and GO ‘cellular component’ enrichment analysis in aging turquoise killifish ovaries.**

**(A)** Boxplots of differentially expressed gene expression groups in ovaries by DEseq2 LRT (FDR < 1e-6). Each dot represents the expression level of a significantly differentially expressed gene in each group normalized by Z-score for ease of group-to-group comparison. **(B)** Top 10 GO "Cellular Component" terms enriched in each associated cluster according to ‘GOstats’ (FDR < 5%; see **Supplemental Table S3B** for the complete list of enriched terms). Terms down at middle-age related to the PIWI pathway include "pi-body" and "P granule" terms. Patterns are labelled according to **Fig. 2A**. FDR: False discovery rate. Enrichment: fold enrichment over background.

**Supplemental Figure S3. Differential gene expression analysis and GO ‘cellular component’ enrichment analysis in aging turquoise killifish testes.**

**(A)** Boxplots of differentially expressed gene expression groups in testes by DEseq2 LRT (FDR < 1e-6). Each dot represents the expression level of a significantly differentially expressed gene in each group normalized by Z-score for ease of group-to-group comparison. **(B)** Top 10 GO "Cellular Component" terms enriched in each associated cluster according to ‘GOstats’ (FDR < 5%; see **Supplemental Table S3E** for the complete list of enriched terms). Most terms enriched at middle-age are related to spermatogenesis. Patterns are labelled according to **Fig. 2A**. FDR: False discovery rate. Enrichment: fold enrichment over background.

**Supplemental Figure S4. Longitudinal histological and deconvolution analyses of turquoise killifish ovaries.**

**(A)** Representative images of 5-, 10-, and 15-week old H&E stained turquoise killifish ovaries. Scale bar: 1mm. **(B)** Boxplots of oocyte diameters grouped by age measured in micrometers by four blinded observers. Distributions of oocyte diameters were compared for young vs. middle-aged ovaries, and middle-aged vs. old ovaries using a Kolmogorov-Smirnov goodness-of-fit test. All comparisons were non-significant (p-value > 0.05), suggesting that the composition of oocytes along maturation stages is largely unchanged throughout turquoise killifish gonadal aging. **(C)** Expression correlation matrix of signature expression profiles from pure ovarian cell types, derived from Zebrafish scRNA-seq dataset from (Liu et al. 2022), using Spearman rank correlation on TPM expression of killifish homologs. Note the clear separation of germline and somatic cells clearly. **(D)** Expression levels (in tpm) of *PIWIL1* in signature expression profiles from pure ovarian cell types show the highest expression in immature germ cells (*i.e.* GSPCs and meiotic oocytes). **(E)** Benchmarking of deconvolution algorithms performance by ‘granulator’, on *in silico* mixed pseudobulk expression of ovarian gene expression derived from the Zebrafish ovary reference dataset. The mean Pearson correlation coefficient over all cell types for each algorithms are used to reflect the overall performance of each algorithm on the in silico mixes of known proportion. Note that SVR and NNLS show the best performance. **(F-G)** Deconvolution results of the aging African turquoise killifish ovarian bulk transcriptomes using SVR (F) or NNLS (G). The predicted proportion of immature germ cells (*PIWIL1*-high) from young, middle-aged, and old deconvoluted turquoise killlifish ovarian bulk transcriptomes. No significant difference was detected between the young and middle-aged samples. Although there were significant changes in the proportions between middle-aged and old ovaries, they are not consistently in the same direction according to both algorithms, Significance in non parametric Wilcoxon tests. ns: non-significant. *: p < 0.05.

**Supplemental Figure S5. TE expression dynamics in the gonads of aging turquoise killifish.**

**(A)** Boxplots of significantly differentially expressed TE clusters in ovaries by DEseq2 LRT (FDR < 1e-6). **(B)** Boxplot of the differentially expressed TE clusters in testes by DEseq2 LRT (FDR < 1e-6). Each dot represents the expression level of a significantly differentially expressed gene in each group normalized by Z-score for ease of group-to-group comparison. Patterns are labelled according to **Fig. 2A**.

**Supplemental Figure S6. piRNA characterization and regulation in aging turquoise killifish gonads.**

**(A)** piRNA length distribution shown by sex, after computational filtering of small RNAs 24-35 bp. Both ovaries and testes have similar piRNA length profiles, with the majority of detected species at ~27bp. **(B)** Scatterplots of young, middle-aged, and old killifish ovaries normalized TE expression *vs.* piRNA counts assigned to the same cognate consensus TE sequence. There is a strong Spearman rank correlation Rho in all ovarian age groups (Rho = 0.66; significance of correlation test: p < 2.2e-16), indicating that piRNAs are produced sustainably throughout life in response to expressed TEs. **(C)** Scatterplots of young, middle-aged, and old killifish testes normalized TE expression *vs.* piRNA counts assigned to the same cognate consensus TE sequence . There is a slightly higher Spearman rank correlation in testes than ovaries (Rho = 0.74-0.75; significance of correlation test: p < 2.2e-16), likely reflecting a larger proportion of germ cells in testicular tissue. Rho: Spearman rank correlation value. **(D)** Boxplots of differentially piRNA-mapped groups of TEs in ovaries, by DEseq2 LRT (FDR < 1e-6), using the grouping arrangement defined in **Fig. 2A**. **(E)** Boxplots of differentially piRNA-mapped groups of TEs in testes by DEseq2 LRT (FDR < 1e-6).

**Supplemental Figure S7. Analysis of ping-pong biogenesis signatures in aging turquoise killifish gonads (continued).**

**(A)** Median Z_i_-scores in ovaries grouped by age measured for each position within a 20 basepair window as summarized for position 10 in **Fig. 6C**. The highest Z_i_-score occurs at position 10 (corresponding to the Z_10_ value), indicating that ping-pong is occurring at all ages. **(B)** Median Z_i_-scores in testes grouped by age measured for each position within a 20 basepair window. Like the ovarian samples, the highest Z_10_-score occurs at position 10 indicating ping-pong activity. **(C)** Boxplots of differential Z_10_ scores of TEs grouped into patterns in ovaries (patterns defined in **Fig. 6D**). Each dot is the Z-normalized Z_10_ score per age group. **(D)** Boxplots of differential Z_10_ scores of TEs grouped into clusters in testes. Each dot is the Z-normalized Z_10_ score per age group.

***Inventory of Supplemental Tables:***

**Supplemental Table S1:** **Lifespan and fecundity data from GRZ African turquoise killifish reported in this study.**

(**A**) GRZ Lifespan Data (expressed in weeks) in the Benayoun laboratory colony at USC.

**Supplemental Table S2:** **List of genes with differential expression significance with age by DESeq2 LRT (FDR < 1e-6), and expression of “piRNA metabolic pathway” GO term associated genes.**

(**A**) Complete Genic Ovarian DESeq2 LRT Analysis with clusters. (**B**) Complete Genic Testicular DESeq2 LRT Analysis with clusters. (**C**) DEseq2 VST-Normalized log_2_(count) gene expression for "piRNA metabolic process" GO term (ovaries and testes). (**D**) Best BLAST hit with E-value < 1E^-3^ for turquoise killifish protein sequences aligning to Human Ensembl 104 protein sequences.

**Supplemental Table S3:** **Enriched GO terms associated to genes with significant age-regulation in ovaries or testes (FDR < 5%).**

(**A**) Ovarian Significantly enriched GO Biological Process Terms by clusters as defined in **Fig. 2A** (GOStats FDR <5%). (**B**) Ovarian Significantly enriched GO Cellular Component Terms by clusters as defined in **Fig. 2A** (GOStats FDR <5%). (**C**) Ovarian Significantly enriched GO Molecular Function Terms by clusters as defined in **Fig. 2A** (GOStats FDR <5%). (**D**) Testicular Significantly enriched GO Biological Process Terms by clusters as defined in **Fig. 2A** (GOStats FDR <5%). (**E**) Testicular Significantly enriched GO Cellular Component Terms by clusters as defined in **Fig. 2A** (GOStats FDR <5%). (**F**) Testicular Significantly enriched GO Molecular Function Terms by clusters as defined in **Fig. 2A** (GOStats FDR <5%).

**Supplemental Table S4:** **List of TEs with differential expression significance with age by DESeq2 LRT (FDR < 1e-6).**

(**A**) Complete TE Ovarian DESeq2 LRT Analysis with clusters as defined in **Fig. 2A**. (**B**) Complete TE Testicular DESeq2 LRT Analysis with clusters as defined in **Fig. 2A**.

**Supplemental Table S5:** **List of piRNA targeted TE sequences with differential abundance significance with age by DESeq2 LRT (FDR < 1e-6).**

(**A**) Complete piRNA mapping to TE Ovarian DESeq2 LRT Analysis with clusters as defined in **Fig. 2A**. (**B**) Complete piRNA mapping to TE Testicular DESeq2 LRT Analysis with clusters as defined in **Fig. 2A**.

**Supplemental Table S6:** **List of piRNA targeted TE sequences with significant differential Z_10_ ping-pong scores with age by ANOVA (FDR < 0.05).**

(**A**) TEs with significantly differential ping-pong Z_10_ scores in ovaries organized by clusters as defined in **Fig. 6D** (ANOVA FDR < 5%). (**B**) TEs with significantly differential ping-pong Z_10_ scores in testes organized by clusters as defined in **Fig. 6D** (ANOVA FDR < 5%).

**Supplementary Files**

**Supplemental Code**: Code used in this study.
