## Supplemental Code for "Dynamic regulation of gonadal transposon control across the lifespan of the naturally short-lived African turquoise killifish": 1_mRNA_hard_trim.sh

```
# hard_trim_mRNAs.sh
#Hard Trimming with fastx_trimmer (fastx_toolkit v0.0.13) to remove
# i)  adapter fragments at the 5' end (remove first 16 nucleotides)
# ii) remove low quality bases (> 100 nts)

#hardtrim young females
gzcat YF1/YF1_1.fq.gz | /Users/bryanteefy/Downloads/bin/fastx_trimmer -f 16 -l 100 -z -i - -Q33 -o ./hard_trimmed_reads/YF1_1_HT.fq.gz
gzcat YF1/YF1_2.fq.gz | /Users/bryanteefy/Downloads/bin/fastx_trimmer -f 16 -l 100 -z -i - -Q33 -o ./hard_trimmed_reads/YF1_2_HT.fq.gz
gzcat YF2/YF2_1.fq.gz | /Users/bryanteefy/Downloads/bin/fastx_trimmer -f 16 -l 100 -z -i - -Q33 -o ./hard_trimmed_reads/YF2_1_HT.fq.gz
gzcat YF2/YF2_2.fq.gz | /Users/bryanteefy/Downloads/bin/fastx_trimmer -f 16 -l 100 -z -i - -Q33 -o ./hard_trimmed_reads/YF2_2_HT.fq.gz
gzcat YF3/YF3_1.fq.gz | /Users/bryanteefy/Downloads/bin/fastx_trimmer -f 16 -l 100 -z -i - -Q33 -o ./hard_trimmed_reads/YF3_1_HT.fq.gz
gzcat YF3/YF3_2.fq.gz | /Users/bryanteefy/Downloads/bin/fastx_trimmer -f 16 -l 100 -z -i - -Q33 -o ./hard_trimmed_reads/YF3_2_HT.fq.gz
gzcat YF4/YF4_1.fq.gz | /Users/bryanteefy/Downloads/bin/fastx_trimmer -f 16 -l 100 -z -i - -Q33 -o ./hard_trimmed_reads/YF4_1_HT.fq.gz
gzcat YF4/YF4_2.fq.gz | /Users/bryanteefy/Downloads/bin/fastx_trimmer -f 16 -l 100 -z -i - -Q33 -o ./hard_trimmed_reads/YF4_2_HT.fq.gz
gzcat YF5/YF5_1.fq.gz | /Users/bryanteefy/Downloads/bin/fastx_trimmer -f 16 -l 100 -z -i - -Q33 -o ./hard_trimmed_reads/YF5_1_HT.fq.gz
gzcat YF5/YF5_2.fq.gz | /Users/bryanteefy/Downloads/bin/fastx_trimmer -f 16 -l 100 -z -i - -Q33 -o ./hard_trimmed_reads/YF5_2_HT.fq.gz

#hardtrim middle-aged females
gzcat MF1/MF1_1.fq.gz | /Users/bryanteefy/Downloads/bin/fastx_trimmer -f 16 -l 100 -z -i - -Q33 -o ./hard_trimmed_reads/MF1_1_HT.fq.gz
gzcat MF1/MF1_2.fq.gz | /Users/bryanteefy/Downloads/bin/fastx_trimmer -f 16 -l 100 -z -i - -Q33 -o ./hard_trimmed_reads/MF1_2_HT.fq.gz
gzcat MF2/MF2_1.fq.gz | /Users/bryanteefy/Downloads/bin/fastx_trimmer -f 16 -l 100 -z -i - -Q33 -o ./hard_trimmed_reads/MF2_1_HT.fq.gz
gzcat MF2/MF2_2.fq.gz | /Users/bryanteefy/Downloads/bin/fastx_trimmer -f 16 -l 100 -z -i - -Q33 -o ./hard_trimmed_reads/MF2_2_HT.fq.gz
gzcat MF3/MF3_1.fq.gz | /Users/bryanteefy/Downloads/bin/fastx_trimmer -f 16 -l 100 -z -i - -Q33 -o ./hard_trimmed_reads/MF3_1_HT.fq.gz
gzcat MF3/MF3_2.fq.gz | /Users/bryanteefy/Downloads/bin/fastx_trimmer -f 16 -l 100 -z -i - -Q33 -o ./hard_trimmed_reads/MF3_2_HT.fq.gz
gzcat MF4/MF4_1.fq.gz | /Users/bryanteefy/Downloads/bin/fastx_trimmer -f 16 -l 100 -z -i - -Q33 -o ./hard_trimmed_reads/MF4_1_HT.fq.gz
gzcat MF4/MF4_2.fq.gz | /Users/bryanteefy/Downloads/bin/fastx_trimmer -f 16 -l 100 -z -i - -Q33 -o ./hard_trimmed_reads/MF4_2_HT.fq.gz
gzcat MF5/MF5_1.fq.gz | /Users/bryanteefy/Downloads/bin/fastx_trimmer -f 16 -l 100 -z -i - -Q33 -o ./hard_trimmed_reads/MF5_1_HT.fq.gz
gzcat MF5/MF5_2.fq.gz | /Users/bryanteefy/Downloads/bin/fastx_trimmer -f 16 -l 100 -z -i - -Q33 -o ./hard_trimmed_reads/MF5_2_HT.fq.gz

#hardtrim old females
gzcat OF1/OF1_1.fq.gz | /Users/bryanteefy/Downloads/bin/fastx_trimmer -f 16 -l 100 -z -i - -Q33 -o ./hard_trimmed_reads/OF1_1_HT.fq.gz
gzcat OF1/OF1_2.fq.gz | /Users/bryanteefy/Downloads/bin/fastx_trimmer -f 16 -l 100 -z -i - -Q33 -o ./hard_trimmed_reads/OF1_2_HT.fq.gz
gzcat OF3/OF3_1.fq.gz | /Users/bryanteefy/Downloads/bin/fastx_trimmer -f 16 -l 100 -z -i - -Q33 -o ./hard_trimmed_reads/OF3_1_HT.fq.gz
gzcat OF3/OF3_2.fq.gz | /Users/bryanteefy/Downloads/bin/fastx_trimmer -f 16 -l 100 -z -i - -Q33 -o ./hard_trimmed_reads/OF3_2_HT.fq.gz
gzcat OF4/OF4_1.fq.gz | /Users/bryanteefy/Downloads/bin/fastx_trimmer -f 16 -l 100 -z -i - -Q33 -o ./hard_trimmed_reads/OF4_1_HT.fq.gz
gzcat OF4/OF4_2.fq.gz | /Users/bryanteefy/Downloads/bin/fastx_trimmer -f 16 -l 100 -z -i - -Q33 -o ./hard_trimmed_reads/OF4_2_HT.fq.gz
gzcat OF5/OF5_1.fq.gz | /Users/bryanteefy/Downloads/bin/fastx_trimmer -f 16 -l 100 -z -i - -Q33 -o ./hard_trimmed_reads/OF5_1_HT.fq.gz
gzcat OF5/OF5_2.fq.gz | /Users/bryanteefy/Downloads/bin/fastx_trimmer -f 16 -l 100 -z -i - -Q33 -o ./hard_trimmed_reads/OF5_2_HT.fq.gz

#hardtrim young males
gzcat YM1/YM1_1.fq.gz | /Users/bryanteefy/Downloads/bin/fastx_trimmer -f 16 -l 100 -z -i - -Q33 -o ./hard_trimmed_reads/YM1_1_HT.fq.gz
gzcat YM1/YM1_2.fq.gz | /Users/bryanteefy/Downloads/bin/fastx_trimmer -f 16 -l 100 -z -i - -Q33 -o ./hard_trimmed_reads/YM1_2_HT.fq.gz
gzcat YM2/YM2_1.fq.gz | /Users/bryanteefy/Downloads/bin/fastx_trimmer -f 16 -l 100 -z -i - -Q33 -o ./hard_trimmed_reads/YM2_1_HT.fq.gz
gzcat YM2/YM2_2.fq.gz | /Users/bryanteefy/Downloads/bin/fastx_trimmer -f 16 -l 100 -z -i - -Q33 -o ./hard_trimmed_reads/YM2_2_HT.fq.gz
gzcat YM3/YM3_1.fq.gz | /Users/bryanteefy/Downloads/bin/fastx_trimmer -f 16 -l 100 -z -i - -Q33 -o ./hard_trimmed_reads/YM3_1_HT.fq.gz
gzcat YM3/YM3_2.fq.gz | /Users/bryanteefy/Downloads/bin/fastx_trimmer -f 16 -l 100 -z -i - -Q33 -o ./hard_trimmed_reads/YM3_2_HT.fq.gz
gzcat YM5/YM5_1.fq.gz | /Users/bryanteefy/Downloads/bin/fastx_trimmer -f 16 -l 100 -z -i - -Q33 -o ./hard_trimmed_reads/YM5_1_HT.fq.gz
gzcat YM5/YM5_2.fq.gz | /Users/bryanteefy/Downloads/bin/fastx_trimmer -f 16 -l 100 -z -i - -Q33 -o ./hard_trimmed_reads/YM5_2_HT.fq.gz

#hardtrim middle-aged males
gzcat MM1/MM1_1.fq.gz | /Users/bryanteefy/Downloads/bin/fastx_trimmer -f 16 -l 100 -z -i - -Q33 -o ./hard_trimmed_reads/MM1_1_HT.fq.gz
gzcat MM1/MM1_2.fq.gz | /Users/bryanteefy/Downloads/bin/fastx_trimmer -f 16 -l 100 -z -i - -Q33 -o ./hard_trimmed_reads/MM1_2_HT.fq.gz
gzcat MM2/MM2_1.fq.gz | /Users/bryanteefy/Downloads/bin/fastx_trimmer -f 16 -l 100 -z -i - -Q33 -o ./hard_trimmed_reads/MM2_1_HT.fq.gz
gzcat MM2/MM2_2.fq.gz | /Users/bryanteefy/Downloads/bin/fastx_trimmer -f 16 -l 100 -z -i - -Q33 -o ./hard_trimmed_reads/MM2_2_HT.fq.gz
gzcat MM3/MM3_1.fq.gz | /Users/bryanteefy/Downloads/bin/fastx_trimmer -f 16 -l 100 -z -i - -Q33 -o ./hard_trimmed_reads/MM3_1_HT.fq.gz
gzcat MM3/MM3_2.fq.gz | /Users/bryanteefy/Downloads/bin/fastx_trimmer -f 16 -l 100 -z -i - -Q33 -o ./hard_trimmed_reads/MM3_2_HT.fq.gz
gzcat MM5/MM5_1.fq.gz | /Users/bryanteefy/Downloads/bin/fastx_trimmer -f 16 -l 100 -z -i - -Q33 -o ./hard_trimmed_reads/MM5_1_HT.fq.gz
gzcat MM5/MM5_2.fq.gz | /Users/bryanteefy/Downloads/bin/fastx_trimmer -f 16 -l 100 -z -i - -Q33 -o ./hard_trimmed_reads/MM5_2_HT.fq.gz

#hardtrim old males
gzcat OM1/OM1_1.fq.gz | /Users/bryanteefy/Downloads/bin/fastx_trimmer -f 16 -l 100 -z -i - -Q33 -o ./hard_trimmed_reads/OM1_1_HT.fq.gz
gzcat OM1/OM1_2.fq.gz | /Users/bryanteefy/Downloads/bin/fastx_trimmer -f 16 -l 100 -z -i - -Q33 -o ./hard_trimmed_reads/OM1_2_HT.fq.gz
gzcat OM3/OM3_1.fq.gz | /Users/bryanteefy/Downloads/bin/fastx_trimmer -f 16 -l 100 -z -i - -Q33 -o ./hard_trimmed_reads/OM3_1_HT.fq.gz
gzcat OM3/OM3_2.fq.gz | /Users/bryanteefy/Downloads/bin/fastx_trimmer -f 16 -l 100 -z -i - -Q33 -o ./hard_trimmed_reads/OM3_2_HT.fq.gz
gzcat OM4/OM4_1.fq.gz | /Users/bryanteefy/Downloads/bin/fastx_trimmer -f 16 -l 100 -z -i - -Q33 -o ./hard_trimmed_reads/OM4_1_HT.fq.gz
gzcat OM4/OM4_2.fq.gz | /Users/bryanteefy/Downloads/bin/fastx_trimmer -f 16 -l 100 -z -i - -Q33 -o ./hard_trimmed_reads/OM4_2_HT.fq.gz
gzcat OM5/OM5_1.fq.gz | /Users/bryanteefy/Downloads/bin/fastx_trimmer -f 16 -l 100 -z -i - -Q33 -o ./hard_trimmed_reads/OM5_1_HT.fq.gz
gzcat OM5/OM5_2.fq.gz | /Users/bryanteefy/Downloads/bin/fastx_trimmer -f 16 -l 100 -z -i - -Q33 -o ./hard_trimmed_reads/OM5_2_HT.fq.gz
```
