## Supplemental Code for "Dynamic regulation of gonadal transposon control across the lifespan of the naturally short-lived African turquoise killifish": 2_mRNA_trim_galore.sh

```
#trim_galore_mRNA.sh

#this script uses trim_galore v0.6.7 to
#remove any remaining adapters and low quality reads

for f in $(find "." -name '*1_HT.fq.gz')
do
    f1=$(basename "${f}");
    f2=$(basename "${f}" | sed 's/1_HT\.fq\.gz/2_HT\.fq\.gz/g');
	trim_galore --paired $f1 $f2 -o trim_galore_out 
done
```
