## Supplemental Code for "Dynamic regulation of gonadal transposon control across the lifespan of the naturally short-lived African turquoise killifish": 3_repeatmasker_genome.sh

```
#repeatmasker_genome.sh

#this script uses RepeatMasker v4.1.2-p1 to
#mask the genome using TE sequences from FishTEDB (Shao et al., 2018)

RepeatMasker -pa 20 -lib Nothobranchius_furzeri_TEs.fa  GCA_014300015.1_MPIA_NFZ_2.0_genomic.fa -xsmall
```
