## Supplemental Code for "Dynamic regulation of gonadal transposon control across the lifespan of the naturally short-lived African turquoise killifish": 4_GTF_generation_TETranscripts.R

```
#This script prepares TE gtf files for TETransripts
#This script uses Repeatmasker .out files that have been pre-processed

#To pre-process Repeatmasker .out files, run the following regular expressions in a text editor:

#Input File: "GCA_014300015.1_MPIA_NFZ_2.0_genomic.fa.out"

#remove * symbol in some final columns to make each row have equal columns
# "\*" to ""

#remove first blank column
# "^\t" to ""

#remove blank final column
# "\t\r" to "\r"

#Save file as: "preprocessed_gtf.fa.out"

###############################################
#set working directory; this will be the same as the output directory
setwd("/Users/bteefy/Dropbox/PIWI/CODE/1_Gene_and_TE_preprocessing")

# R version 4.1.2 (2021-11-01)

#load libraries
library(gdata) #gdata_2.18.0
library(tidyr) #tidyr_1.1.4 
library(dplyr) #dplyr_1.0.7

#define outprefix
my.outprefix <- paste(Sys.Date(),"GTF_generation",sep="_")

#read in modulated Repeatmasker .out file; Code-checkers: unzip first (gzip file)
outfile <- read.table("./Input/preprocessed_gtf.fa.out", sep = "\t", header = F)

#remove simple repeats and low complexity regions
outfile <- outfile[!grepl("Simple", outfile$V11),]
outfile <- outfile[grepl("NotFur", outfile$V10),]

#change complementary ("C") to "-"
outfile$V9 <- gsub("C", "-",outfile$V9)

#keep only positinal info and names; strand
outfile <- outfile[,c(5:7,9:11)]

#separate into family types with unknowns/unclears being filled in when they throw up NAs
outfile_sep <- outfile %>% separate(V11, c("A", "B"),sep = "/")
outfile_sep$B <- ifelse(is.na(outfile_sep$B), outfile_sep$A, outfile_sep$B)

#remove bulky outfile for processing speed/memory concerns
rm(outfile)

#populate columns
outfile_sep$transcript_id <- outfile_sep$V10
outfile_sep$TE <- "TE"
outfile_sep$exon <- "exon"
outfile_sep$period1 <- "."
outfile_sep$period2 <- "."
colnames(outfile_sep) <- c("chr", "start", "stop", "strand", "gene_id", "class_id", "family_id", "transcript_id", "TE", "exon", "period1", "period2")
outfile_sep <- outfile_sep[,c("chr", "TE", "exon", "start", "stop", "period1", "strand", "period2", "gene_id", "transcript_id","family_id", "class_id")]

# combine new metrics into a single final column; use quotes to help track everything clearly
outfile_sep$name <- paste0('gene_id "',outfile_sep$gene_id,'"; transcript_id "',
                           outfile_sep$transcript_id,'"; family_id "',outfile_sep$family_id,
                           '"; class_id "',outfile_sep$class_id,'";')

outfile_sep <- outfile_sep[,c("chr", "TE", "exon", "start", "stop", "period1", "strand", "period2", "name")]

#now remove quotes for use in TETranscripts
outfile_sep$name <- gsub('"', '', outfile_sep$name)

#export gtf
write.table(outfile_sep, file = "./Results/TE_gtf_for_TETranscripts.gtf",
            row.names = F, col.names = F, quote = F, sep = "\t")

#######################
sink(file = paste(my.outprefix,"_session_Info.txt", sep =""))
sessionInfo()
sink()
```
