## Supplemental Code for "Dynamic regulation of gonadal transposon control across the lifespan of the naturally short-lived African turquoise killifish": 5_star_genome_index.sh

```
#star_genome_index.sh

#this script uses star v2.7.0e to index the soft-masked genome

STAR --runThreadN 2 --runMode genomeGenerate --genomeDir ./star_genomic_index --limitGenomeGenerateRAM 60000000000 --genomeFastaFiles Soft_Masked_Genome.fa --sjdbGTFfile GCA_014300015.1_MPIA_NFZ_2.0_genomic.gtf
```
