## Supplemental Code for "Dynamic regulation of gonadal transposon control across the lifespan of the naturally short-lived African turquoise killifish": 6_map_mRNAs_to_genome.sh

```
#map_mRNAs_to_genome.sh

#this script uses star v2.7.0e to map mRNA fastq files to the softmasked genome

for f in $(find "." -name '*_1_HT_val_1.fq.gz')
do
    f2=$(basename "${f}" | sed 's/_1_HT_val_1\.fq\.gz/_2_HT_val_2\.fq\.gz/g');
    of=$(basename "${f}" | sed 's/_1_HT_val_1\.fq\.gz//g');
    
    STAR --genomeDir star_genomic_index --readFilesIn $f $f2 --readFilesCommand zcat --runThreadN 6 --outFilterMultimapNmax 200 --outFilterIntronMotifs RemoveNoncanonicalUnannotated --alignEndsProtrude 10 ConcordantPair --limitGenomeGenerateRAM 60000000000 --outSAMtype BAM SortedByCoordinate --outFileNamePrefix $of
done
```
