## Supplemental Code for "Dynamic regulation of gonadal transposon control across the lifespan of the naturally short-lived African turquoise killifish": 7_mRNA_counting_TETranscripts.sh

```
#mRNA_counting_TETranscripts.sh

#this script uses TEtranscripts v2.2.1 to count gene and TE reads from star mapping
#the -t and -c options are necessary but irrelevant since there are no proper control or treatment groups
#outputs a count matrix for all replicates

TEtranscripts -t MF1.bam MF2.bam MF3.bam MF4.bam MF5.bam MM1.bam MM2.bam MM3.bam MM5.bam OF1.bam OF3.bam OF4.bam OF5.bam OM1.bam OM3.bam OM4.bam OM5.bam -c YF1.bam YF2.bam YF3.bam YF4.bam YF5.bam YM1.bam YM2.bam YM3.bam YM5.bam --GTF GCA_014300015.1_MPIA_NFZ_2.0_genomic.gtf --TE TE_gtf_for_TETranscripts.gtf --sortByPos --project Gene_TE_Counts
```
