## Supplemental Code for "Dynamic regulation of gonadal transposon control across the lifespan of the naturally short-lived African turquoise killifish": README_Gene_and_TE_Preprocessing.txt

########################################################
		README -- 1_Gene_and_TE_Preprocessing
########################################################

# Software versions are indicated in each cognate script as comments

	1. mRNA reads were hard-trimmed with fastx_trimmer
		- 1_mRNA_hard_trim.sh
	
	2. mRNA reads were trimmed of remaining adapters with trim_galore 
		- 2_mRNA_trim_galore.sh
	
	3. Soft-mask reference genome with Repeatmasker
		- 3_repeatmasker_genome.sh
	
	4. Custom TE-specific generation for counting with TETranscripts 
		- 4_GTF_generation_TETranscripts.R
	
	5. Genome indexing for STAR mapping of mRNA sequences to the genome 
		- 5_star_genome_index.sh
	
	6. Map mRNAs to reference genome with STAR (200 multimappers allowed for TEtranscript step)
		- 6_map_mRNAs_to_genome.sh
	
	7. Quantify mRNA and TE counts with TETranscripts 
		- 7_mRNA_counting_TETranscripts.sh
