## Supplemental Code for "Dynamic regulation of gonadal transposon control across the lifespan of the naturally short-lived African turquoise killifish": 1_small_RNA_trimming.sh

```
#small_RNA_trimming.sh

# this script performs small RNA trimming with trim_galore v0.6.7 to remove specific adapters and low quality reads
# discard reads that become shorter than 18bp
# each file is trimmed twice. once for the forward and reverse adapters

RAWDIR="/Volumes/easystore/small_RNA/usftp21.novogene.com/raw_data"
QUALDIR="/Volumes/easystore/small_RNA/usftp21.novogene.com/raw_data/Trim_Galore" 

#young females
trim_galore --length 18 -a GTTCAGAGTTCTACAGTCCGACGATC $RAWDIR/YF1/YF1.fq.gz		-o $QUALDIR
trim_galore --length 18 -a AGATCGGAAGAGCACACGTCT $QUALDIR/YF1_trimmed.fq.gz -o $QUALDIR
trim_galore --length 18 -a GTTCAGAGTTCTACAGTCCGACGATC $RAWDIR/YF2/YF2.fq.gz		-o $QUALDIR
trim_galore --length 18 -a AGATCGGAAGAGCACACGTCT $QUALDIR/YF2_trimmed.fq.gz -o $QUALDIR
trim_galore --length 18 -a GTTCAGAGTTCTACAGTCCGACGATC $RAWDIR/YF3/YF3.fq.gz		-o $QUALDIR
trim_galore --length 18 -a AGATCGGAAGAGCACACGTCT $QUALDIR/YF3_trimmed.fq.gz -o $QUALDIR
trim_galore --length 18 -a GTTCAGAGTTCTACAGTCCGACGATC $RAWDIR/YF4/YF4.fq.gz		-o $QUALDIR
trim_galore --length 18 -a AGATCGGAAGAGCACACGTCT $QUALDIR/YF4_trimmed.fq.gz -o $QUALDIR
trim_galore --length 18 -a GTTCAGAGTTCTACAGTCCGACGATC $RAWDIR/YF5/YF5.fq.gz		-o $QUALDIR
trim_galore --length 18 -a AGATCGGAAGAGCACACGTCT $QUALDIR/YF5_trimmed.fq.gz -o $QUALDIR
 
#middle-aged females
trim_galore --length 18 -a GTTCAGAGTTCTACAGTCCGACGATC $RAWDIR/MF1/MF1.fq.gz		-o $QUALDIR
trim_galore --length 18 -a AGATCGGAAGAGCACACGTCT $QUALDIR/MF1_trimmed.fq.gz -o $QUALDIR
trim_galore --length 18 -a GTTCAGAGTTCTACAGTCCGACGATC $RAWDIR/MF2/MF2.fq.gz		-o $QUALDIR
trim_galore --length 18 -a AGATCGGAAGAGCACACGTCT $QUALDIR/MF2_trimmed.fq.gz -o $QUALDIR
trim_galore --length 18 -a GTTCAGAGTTCTACAGTCCGACGATC $RAWDIR/MF3/MF3.fq.gz		-o $QUALDIR
trim_galore --length 18 -a AGATCGGAAGAGCACACGTCT $QUALDIR/MF3_trimmed.fq.gz -o $QUALDIR
trim_galore --length 18 -a GTTCAGAGTTCTACAGTCCGACGATC $RAWDIR/MF4/MF4.fq.gz		-o $QUALDIR
trim_galore --length 18 -a AGATCGGAAGAGCACACGTCT $QUALDIR/MF4_trimmed.fq.gz -o $QUALDIR
trim_galore --length 18 -a GTTCAGAGTTCTACAGTCCGACGATC $RAWDIR/MF5/MF5.fq.gz		-o $QUALDIR
trim_galore --length 18 -a AGATCGGAAGAGCACACGTCT $QUALDIR/MF5_trimmed.fq.gz -o $QUALDIR
 
#old females
trim_galore --length 18 -a GTTCAGAGTTCTACAGTCCGACGATC $RAWDIR/OF1/OF1.fq.gz		-o $QUALDIR
trim_galore --length 18 -a AGATCGGAAGAGCACACGTCT $QUALDIR/OF1_trimmed.fq.gz -o $QUALDIR
trim_galore --length 18 -a GTTCAGAGTTCTACAGTCCGACGATC $RAWDIR/OF2/OF2.fq.gz		-o $QUALDIR
trim_galore --length 18 -a AGATCGGAAGAGCACACGTCT $QUALDIR/OF2_trimmed.fq.gz -o $QUALDIR
trim_galore --length 18 -a GTTCAGAGTTCTACAGTCCGACGATC $RAWDIR/OF3/OF3.fq.gz		-o $QUALDIR
trim_galore --length 18 -a AGATCGGAAGAGCACACGTCT $QUALDIR/OF3_trimmed.fq.gz -o $QUALDIR
trim_galore --length 18 -a GTTCAGAGTTCTACAGTCCGACGATC $RAWDIR/OF4/OF4.fq.gz		-o $QUALDIR
trim_galore --length 18 -a AGATCGGAAGAGCACACGTCT $QUALDIR/OF4_trimmed.fq.gz -o $QUALDIR
trim_galore --length 18 -a GTTCAGAGTTCTACAGTCCGACGATC $RAWDIR/OF5/OF5.fq.gz		-o $QUALDIR
trim_galore --length 18 -a AGATCGGAAGAGCACACGTCT $QUALDIR/OF5_trimmed.fq.gz -o $QUALDIR

#young males
trim_galore --length 18 -a GTTCAGAGTTCTACAGTCCGACGATC $RAWDIR/YM1/YM1.fq.gz		-o $QUALDIR
trim_galore --length 18 -a AGATCGGAAGAGCACACGTCT $QUALDIR/YM1_trimmed.fq.gz -o $QUALDIR
trim_galore --length 18 -a GTTCAGAGTTCTACAGTCCGACGATC $RAWDIR/YM2/YM2.fq.gz		-o $QUALDIR
trim_galore --length 18 -a AGATCGGAAGAGCACACGTCT $QUALDIR/YM2_trimmed.fq.gz -o $QUALDIR
trim_galore --length 18 -a GTTCAGAGTTCTACAGTCCGACGATC $RAWDIR/YM3/YM3.fq.gz		-o $QUALDIR
trim_galore --length 18 -a AGATCGGAAGAGCACACGTCT $QUALDIR/YM3_trimmed.fq.gz -o $QUALDIR
trim_galore --length 18 -a GTTCAGAGTTCTACAGTCCGACGATC $RAWDIR/YM4/YM4.fq.gz		-o $QUALDIR
trim_galore --length 18 -a AGATCGGAAGAGCACACGTCT $QUALDIR/YM4_trimmed.fq.gz -o $QUALDIR
trim_galore --length 18 -a GTTCAGAGTTCTACAGTCCGACGATC $RAWDIR/YM5/YM5.fq.gz		-o $QUALDIR
trim_galore --length 18 -a AGATCGGAAGAGCACACGTCT $QUALDIR/YM5_trimmed.fq.gz -o $QUALDIR

#middle-aged males
trim_galore --length 18 -a GTTCAGAGTTCTACAGTCCGACGATC $RAWDIR/MM1/MM1.fq.gz		-o $QUALDIR
trim_galore --length 18 -a AGATCGGAAGAGCACACGTCT $QUALDIR/MM1_trimmed.fq.gz -o $QUALDIR
trim_galore --length 18 -a GTTCAGAGTTCTACAGTCCGACGATC $RAWDIR/MM2/MM2.fq.gz		-o $QUALDIR
trim_galore --length 18 -a AGATCGGAAGAGCACACGTCT $QUALDIR/MM2_trimmed.fq.gz -o $QUALDIR
trim_galore --length 18 -a GTTCAGAGTTCTACAGTCCGACGATC $RAWDIR/MM3/MM3.fq.gz		-o $QUALDIR
trim_galore --length 18 -a AGATCGGAAGAGCACACGTCT $QUALDIR/MM3_trimmed.fq.gz -o $QUALDIR
trim_galore --length 18 -a GTTCAGAGTTCTACAGTCCGACGATC $RAWDIR/MM4/MM4.fq.gz		-o $QUALDIR
trim_galore --length 18 -a AGATCGGAAGAGCACACGTCT $QUALDIR/MM4_trimmed.fq.gz -o $QUALDIR
trim_galore --length 18 -a GTTCAGAGTTCTACAGTCCGACGATC $RAWDIR/MM5/MM5.fq.gz		-o $QUALDIR
trim_galore --length 18 -a AGATCGGAAGAGCACACGTCT $QUALDIR/MM5_trimmed.fq.gz -o $QUALDIR

#old males
trim_galore --length 18 -a GTTCAGAGTTCTACAGTCCGACGATC $RAWDIR/OM1/OM1.fq.gz		-o $QUALDIR
trim_galore --length 18 -a AGATCGGAAGAGCACACGTCT $QUALDIR/OM1_trimmed.fq.gz -o $QUALDIR
trim_galore --length 18 -a GTTCAGAGTTCTACAGTCCGACGATC $RAWDIR/OM2/OM2.fq.gz		-o $QUALDIR
trim_galore --length 18 -a AGATCGGAAGAGCACACGTCT $QUALDIR/OM2_trimmed.fq.gz -o $QUALDIR
trim_galore --length 18 -a GTTCAGAGTTCTACAGTCCGACGATC $RAWDIR/OM3/OM3.fq.gz		-o $QUALDIR
trim_galore --length 18 -a AGATCGGAAGAGCACACGTCT $QUALDIR/OM3_trimmed.fq.gz -o $QUALDIR
trim_galore --length 18 -a GTTCAGAGTTCTACAGTCCGACGATC $RAWDIR/OM4/OM4.fq.gz		-o $QUALDIR
trim_galore --length 18 -a AGATCGGAAGAGCACACGTCT $QUALDIR/OM4_trimmed.fq.gz -o $QUALDIR
trim_galore --length 18 -a GTTCAGAGTTCTACAGTCCGACGATC $RAWDIR/OM5/OM5.fq.gz		-o $QUALDIR
trim_galore --length 18 -a AGATCGGAAGAGCACACGTCT $QUALDIR/OM5_trimmed.fq.gz -o $QUALDIR
```
