## Supplemental Code for "Dynamic regulation of gonadal transposon control across the lifespan of the naturally short-lived African turquoise killifish": 2_size_split_sRNAs.sh

```
#size_split_sRNAs.sh

for f in $(find "." -name '*_trimmed_trimmed.fq.gz')
do

f2=$(basename "${f}")

of_miRNA=$(basename "${f}" | sed 's/_trimmed_trimmed\.fq\.gz/_miRNA\.fq\.gz/g');
of_piRNA=$(basename "${f}" | sed 's/_trimmed_trimmed\.fq\.gz/_piRNA\.fq\.gz/g');

zcat $f2 | awk 'BEGIN {FS = "\t" ; OFS = "\n"} {header = $0 ; getline seq ; getline qheader ; getline qseq ; if (length(seq) >= 20 && length(seq) <= 23) {print header, seq, qheader, qseq}}' | gzip > $of_miRNA
zcat $f2 | awk 'BEGIN {FS = "\t" ; OFS = "\n"} {header = $0 ; getline seq ; getline qheader ; getline qseq ; if (length(seq) >= 24 && length(seq) <= 35) {print header, seq, qheader, qseq}}' | gzip > $of_piRNA

done
```
