## Supplemental Code for "Dynamic regulation of gonadal transposon control across the lifespan of the naturally short-lived African turquoise killifish": 3_piRNA_length_generation.sh

```
#piRNA_length_generation.sh

#Obtain piRNA length data

for f in $(find "." -name '*_piRNA.fq.gz')
        do
        output=$(basename "${f}" | sed 's/_piRNA\.fq\.gz/_piRNA_length\.txt/g');

gunzip -c $f | awk '{if(NR%4==1) {printf(">%s\n",substr($0,2));} else if(NR%4==2) print;}'| awk '$0 ~ ">" {print c; c=0;printf substr($0,2,10) "\t"; } $0 !~ ">" {c+=length($0);} END { print c; }' | sed '/^$/d' | awk '{print $2}' | sort | uniq -c | awk '{print $1,$2}' > $output

done
```
