## Supplemental Code for "Dynamic regulation of gonadal transposon control across the lifespan of the naturally short-lived African turquoise killifish": 4_preprocess_logo_plots.sh

```
#preprocess_logo_plots.sh
# pre-process piRNA fastq files for logo plot generation
# creates a fasta file with headers removed

# uses bedtools v2.27.1

#females

for f in $(find "." -name '*F*_piRNA.fq.gz')
        do
        output=$(basename "${f}" | sed 's/_piRNA\.fq\.gz/_piRNA_reduced_females\.fa/g');


gunzip -c $f | awk '{if(NR%4==1) {printf(">%s\n",substr($0,2));} else if(NR%4==2) print;}' | awk '/^>/{ seqlen=0; print; next; } seqlen < 24 { if (seqlen + length($0) > 24) $0 = substr($0, 1, 24-seqlen); seqlen += length($0); print }' | sed '/^>/d' > $output

done

#replace T with U

cat *_piRNA_reduced_females.fa > total_fasta_females.fasta

sed 's/T/U/g' total_fasta_females.fasta > total_fasta_females_replaced.fasta

#males

for f in $(find "." -name '*M*_piRNA.fq.gz')
        do
        output=$(basename "${f}" | sed 's/_piRNA\.fq\.gz/_piRNA_reduced_males\.fa/g');


gunzip -c $f | awk '{if(NR%4==1) {printf(">%s\n",substr($0,2));} else if(NR%4==2) print;}' | awk '/^>/{ seqlen=0; print; next; } seqlen < 24 { if (seqlen + length($0) > 24) $0 = substr($0, 1, 24-seqlen); seqlen += length($0); print }' | sed '/^>/d' > $output

done

cat *_piRNA_reduced_males.fa > total_fasta_males.fasta

sed 's/T/U/g' total_fasta_males.fasta > total_fasta_males_replaced.fasta
```
