## Supplemental Code for "Dynamic regulation of gonadal transposon control across the lifespan of the naturally short-lived African turquoise killifish": 5_piRNA_fq_to_fa.sh

```
#piRNA_fq_to_fa.sh

#converts piRNA fastq files into fasta format files for use in Protrac

for f in $(find "." -name '*_piRNA.fq.gz')
do
    f1=$(basename "${f}");
    f2=$(basename "${f}" | sed 's/_piRNA\.fq\.gz/_piRNA\.fa/g');

    gunzip -c $f1 | awk '{if(NR%4==1) {printf(">%s\n",substr($0,2));} else if(NR%4==2) print;}' > $f2

done
```
