## Supplemental Code for "Dynamic regulation of gonadal transposon control across the lifespan of the naturally short-lived African turquoise killifish": 6_piRNA_cluster_identification_protrac.sh

```
#piRNA_cluster_identification_protrac.sh

#This script 1. Collapses piRNAs to unique reads with a counter
		   # 2. Maps them
		   # 3. Identifies clusters

#uses scripts from proTRAC v2.42 (Rosenkranz, 2012, BMC Bioinformatics)
#uses perl v5.30.1

tb_coll_path="/path/to/TBr2_collapse.pl"
tb_dust_path="/path/to/TBr2_duster.pl"
map_path="/path/to/sRNAmapper.pl"
genome_path="/path/to/Masked_Genome.fa"
protrac_path="/path/to/proTRAC_2.4.3.pl"

for f in $(find "." -name '*_piRNA.fa')
do
        f1=$(basename "${f}");
        of1=$(basename "${f}" | sed 's/\.fa/\.collapsed/g');
        of2=$(basename "${f}" | sed 's/\.fa/\.collapsed\.no-dust/g');
        map=$(basename "${f}" | sed 's/\.fa/\.collapsed\.no-dust\.map/g');
        
perl $tb_coll_path -i $f1 -o $of1

perl $tb_dust_path -i $of1

perl $map_path -input $of2 -genome $genome_path -alignments best

perl $protrac_path -map $map -genome $genome
        
done
```
