## Supplemental Code for "Dynamic regulation of gonadal transposon control across the lifespan of the naturally short-lived African turquoise killifish": 7_modified_merge.pl

```
#modified_merge.pl

#This script merges clusters identified in piRNA_cluster_identification_protrac.sh
#This merges any clusters within 10 kb into a single cluster. This was done for all (males and females included)
#from proTRAC v2.42


#protrac will generate directories using system time. note that the names "proTRAC*collapsed.no-dust.map*" will generate differently named directories depending on when it is run

# The following array must contain a list of proTRAC output folders.
# Listed proTRAC runs must base on the same genome. Otherwise,
# merging cluster coordinates makes no sense!

@proTRAC_folders=
        (
"proTRAC_MF1_piRNA.collapsed.no-dust.map_2021y7m29d18h56m39s","proTRAC_MF2_piRNA.collapsed.no-dust.map_2021y7m30d3h52m32s","proTRAC_MF3_piRNA.collapsed.no-dust.map_2021y7m29d18h16m56s","proTRAC_MF4_piRNA.collapsed.no-dust.map_2021y7m30d3h6m6s","proTRAC_MF5_piRNA.collapsed.no-dust.map_2021y7m29d20h50m52s","proTRAC_MM1_piRNA.collapsed.no-dust.map_2021y7m30d11h40m54s","proTRAC_MM2_piRNA.collapsed.no-dust.map_2021y7m29d14h23m53s","proTRAC_MM3_piRNA.collapsed.no-dust.map_2021y7m30d9h51m5s","proTRAC_MM5_piRNA.collapsed.no-dust.map_2021y7m30d8h43m9s","proTRAC_OF1_piRNA.collapsed.no-dust.map_2021y7m30d7h28m20s","proTRAC_OF3_piRNA.collapsed.no-dust.map_2021y7m30d8h3m33s","proTRAC_OF4_piRNA.collapsed.no-dust.map_2021y7m29d13h41m22s","proTRAC_OF5_piRNA.collapsed.no-dust.map_2021y7m30d11h8m47s","proTRAC_OM1_piRNA.collapsed.no-dust.map_2021y7m29d19h38m47s","proTRAC_OM3_piRNA.collapsed.no-dust.map_2021y7m29d21h24m50s","proTRAC_OM4_piRNA.collapsed.no-dust.map_2021y7m30d4h36m37s","proTRAC_OM5_piRNA.collapsed.no-dust.map_2021y7m29d17h18m18s","proTRAC_YF1_piRNA.collapsed.no-dust.map_2021y7m29d12h1m10s","proTRAC_YF2_piRNA.collapsed.no-dust.map_2021y7m30d6h37m23s","proTRAC_YF3_piRNA.collapsed.no-dust.map_2021y7m29d12h45m32s","proTRAC_YF4_piRNA.collapsed.no-dust.map_2021y7m30d5h44m19s","proTRAC_YF5_piRNA.collapsed.no-dust.map_2021y7m30d14h13m56s","proTRAC_YM1_piRNA.collapsed.no-dust.map_2021y7m30d0h20m32s","proTRAC_YM2_piRNA.collapsed.no-dust.map_2021y7m29d15h43m2s","proTRAC_YM3_piRNA.collapsed.no-dust.map_2021y7m30d1h33m54s","proTRAC_YM5_piRNA.collapsed.no-dust.map_2021y7m29d22h43m45s"
);

$min_dist=10000; # This is the minimum distance between two independent piRNA clusters. Clusters closer to each other will be merged.
$output_table="merged_cluster_coordinates.txt";
$|=1;

%clusters=();
foreach$folder(@proTRAC_folders)
        {
        open(TABLE,"$folder/results.table")||print"\nCannot open $folder/results.table.\n$!\n\n";
        while(<TABLE>)
                {
                if($_=~/^Cluster/)
                        {
                        $_=~s/Location: [^\t]+//;
                        $loc=$&;
                        $loc=~s/Location: //;
                        $_=~s/Coordinates: [^\t]+//;
                        $coord=$&;
                        $coord=~s/Coordinates: //;
                        @coord=split('-',$coord);
                        foreach$p($coord[0]..$coord[1])
                                {
                                $clusters{$loc}{$p}++;
                                }
                        print"\n$loc -> $coord[0] .. $coord[1]";
                        }
                }
        close TABLE;
        }

open(MERGE,">$output_table");
$prev_loc="";
$cluster_id=0;
foreach$loc(sort{$a cmp $b}keys%clusters)
        {
        $prev_p=-$min_dist;
        $cluster_id++;
        $start=0;
        foreach$p(sort{$a<=>$b}keys%{$clusters{$loc}})
                {
                if($start==0)
                        {
                        print MERGE"$cluster_id\t$loc\t$p";
                        $start=1;
                        }
                if($p>($prev_p+$min_dist)&&$start==1&&$prev_p!=-$min_dist)
                        {
                        print MERGE"\t$prev_p\n";
                        $cluster_id++;
                        $start=0;
                        }
                $prev_p=$p;
                }
        print MERGE"\t$prev_p\n";
        }
close MERGE;
exit;
```
