## Supplemental Code for "Dynamic regulation of gonadal transposon control across the lifespan of the naturally short-lived African turquoise killifish": 9a_TE_repeat_content_in_clusters.R

```
# setwd("/Users/bryanteefy/Dropbox/PIWI/CODE/5_piRNA_Analysis")
setwd("/Users/berenice/Dropbox/Manuscripts_and_Publications/2022/2022_Bryan_PIWI_manuscript/CODE/5_piRNA_Analysis")

#This script generates stacked barplots comparing TE content in the entire genome to that of piRNA clusters

# R version 4.1.2 (2021-11-01)

#load libraries
library(dplyr)     # dplyr_1.0.9 
library(ggplot2)   # ggplot2_3.3.5

my.outprefix <- paste(Sys.Date(),"Cluster",sep="_")

#read in the outfile from repeatmasker
out_file <- read.table("./Input/GCA_014300015.1_MPIA_NFZ_2.0_genomic.CLEAN.fa.out", comment.char = "", header = F, fill = T)

#remove two-line header (input line number OK)
out_file <- out_file[c(3:3677862),]

#retains only 1)chromosome 2)start 3)stop 4)TE name 5)Family Name
bed_file <- out_file[,c(5:7,10:11)]

#Assess how the families breakdown for the whole genome
#remove empty lines 
bed_file <- bed_file[!apply(bed_file == "", 1, all),]

#output bed file
write.table(bed_file, file = "./Input/repeatmasker_bedfile.bed", sep = "\t", col.names = F, row.names = F, quote = F)

#retain only major family name and combine unclear and Unknown
bed_file$V11 <- gsub("/.*","",bed_file$V11)
bed_file$V11 <- gsub("unclear","Unknown",bed_file$V11)

#make a table of the families of TEs. How frequently do the different TE families appear in the genome?
table_of_TEs <- as.data.frame(table(bed_file$V11))

#remove simple repeats and unify "unknown" and "unclear"
table_of_TEs <- table_of_TEs[(!grepl("Low_complexity", table_of_TEs$Var1)),]
table_of_TEs <- table_of_TEs[(!grepl("Simple_repeat", table_of_TEs$Var1)),]

#create matrix with TE family frequency in the genome
cols.g <- rainbow(nrow(table_of_TEs))
table_of_TEs$percent = round(100*table_of_TEs$Freq/sum(table_of_TEs$Freq), digits = 1)
table_of_TEs$label = paste(table_of_TEs$Var1," (", table_of_TEs$percent,"%)", sep = "")


#Assess how the families breakdown for piRNA clusters

#locally, run:
#bedtools intersect -wo -a merged_cluster_coordinates.CLEAN.bed -b repeatmasker_bedfile.bed > 2022-07-27_piRNA_cluster_TEs_intersect.txt
#this command will output the TE composition of the piRNA clusters

#read in the output of bedtools intersect
cluster_TEs <- read.table("./Input/2022-07-27_piRNA_cluster_TEs_intersect.txt")

#repeat analysis as above with piRNA cluster TEs
cluster_TEs$V9 <- gsub("/.*","",cluster_TEs$V9)
cluster_TEs$V9 <- gsub("unclear","Unknown",cluster_TEs$V9)
table_of_cluster_TEs <- as.data.frame(table(cluster_TEs$V9))
cols <- rainbow(nrow(table_of_cluster_TEs))
table_of_cluster_TEs$percent = round(100*table_of_cluster_TEs$Freq/sum(table_of_cluster_TEs$Freq), digits = 1)
table_of_cluster_TEs$label = paste(table_of_cluster_TEs$Var1," (", table_of_cluster_TEs$percent,"%)", sep = "")

#generate stacked barplot showing TE distribution of clusters vs whole genome
table_of_TEs$origin <- "genome"
table_of_cluster_TEs$origin <- "cluster"
clust <- table_of_cluster_TEs[,c(5,1,3)]
genome <- table_of_TEs[,c(5,1,3)]
all <- rbind(clust, genome)
colnames(all)[2] <- "family"

# Stacked barplot
my.stack.out <- paste0("./Results/3_TE_repeat_content_in_clusters/", my.outprefix,"cluster_stack_plot.pdf")
pdf(my.stack.out)
ggplot(all, aes(fill=family, y=percent, x=origin)) + 
  geom_bar(position="stack", stat="identity") + theme_bw() + 
  scale_fill_manual("legend", values = c(DNA="firebrick1", LINE="darkturquoise", LTR="cornflowerblue", SINE="blue", Unknown="gold"))
dev.off()

#######################
sink(file = paste("./Results/3_TE_repeat_content_in_clusters/", my.outprefix,"_session_Info.txt", sep =""))
sessionInfo()
sink()
```
