## Supplemental Code for "Dynamic regulation of gonadal transposon control across the lifespan of the naturally short-lived African turquoise killifish": 11_piRNA_mapping_to_genome.sh

```
#piRNA_mapping_to_genome.sh

#map piRNAs to the softmasked 2020 Killifish Genome (MPIA_NFZ_2.0, Willemsen, 2020)
#this maps all piRNAs allowing up to 3 mismatches

# use modules:
# bowtie v1.2.3
# samtools v1.10

for f in $(find "." -name '*_piRNA.fq.gz')
do
        f2=$(basename "${f}");
        of=$(basename "${f}" | sed 's/_piRNA\.fq\.gz/_mapped/g');
        of_bam="${of}.bam"
        of_sorted="${of}_sorted"
        of_sorted_bam="${of_sorted}.bam"
        of_sorted_bambai="${of_sorted}.bam.bai"
        of_sorted_bai="${of_sorted}.bai"

        bowtie -v 3 -a --best --strata -S bowtie_combined_index $f2 $of
        samtools view -bS -o $of_bam $of
        rm $of
        samtools sort $of_bam -o $of_sorted_bam
        rm $of_bam
        samtools index $of_sorted_bam
        mv $of_sorted_bambai $of_sorted_bai

done
```
