## Supplemental Code for "Dynamic regulation of gonadal transposon control across the lifespan of the naturally short-lived African turquoise killifish": 12_Prepare_rRNA_for_gtf.txt

#### blat - Standalone BLAT v. 36 fast sequence search command line tool

# Run blat to match NCBI rRNA sequence for killifish (gi|194399202|gb|EU780557.1|) to genome reference
blat GCA_014300015.1_MPIA_NFZ_2.0_genomic.fa Nfur_rRNA_sequence.fasta -out=blast8 rRNA_mapping_to_GCA_014300015.1_MPIA_NFZ_2.0_genomic.blast8

# the blast8 format has the below columns (according to https://gist.github.com/brantfaircloth/944741/695d9f2d633d44c5a797ce3540264a4228d5e722)
text = Query id, Subject id, % identity, alignment length, mismatches, gap openings, q. start, q. end, s. start, s. end, e-value, bit score

# filter blat output for matches of > 500bp, no gaps and >95% sequence identity
# format the output as bed file (start pos < end position)
cat rRNA_mapping_to_GCA_014300015.1_MPIA_NFZ_2.0_genomic.blast8 | perl -lane 'print if (($F[3] > 500) && ($F[5] == 0) && ($F[2] > 95))' | perl -lane 'if ($F[8] < $F[9]) {print "$F[1]\t$F[8]\t$F[9]\trRNA_match_$.\t+"} else {print"$F[1]\t$F[9]\t$F[8]\trRNA_match_$.\t-"}' > rRNA_mapping_to_GCA_014300015.1_MPIA_NFZ_2.0_genomic.filt.bed


# run 13_Make_Featurecounts_gtf.R to output a gtf with gene, transcript and exon entries for rRNA and other features of interest
