## Supplemental Code for "Dynamic regulation of gonadal transposon control across the lifespan of the naturally short-lived African turquoise killifish": 13_Make_Featurecounts_gtf.R

```
# set working directory to directory containing pre-gtf files
 setwd("/Users/bryanteefy/Dropbox/PIWI/CODE/2_piRNA_preprocessing")
#setwd("/Users/berenice/Dropbox/Manuscripts_and_Publications/2022/2022_Bryan_PIWI_manuscript/CODE/2_piRNA_preprocessing")

# Generate featurecounts GTF

# R version 4.1.2 (2021-11-01)

#load libraries
library(gdata) # gdata_2.18.0
library(tidyr) # tidyr_1.1.4
library(dplyr) # dplyr_1.0.7

#1. Prepare rRNA gtf
my.bed <- read.table('./Input/rRNA_mapping_to_GCA_014300015.1_MPIA_NFZ_2.0_genomic.filt.bed', sep = "\t", header = F)

# 1. create a "gene" entry
gene.obj             <- my.bed
colnames(gene.obj)   <- c("chr","start","stop", "name","strand")
gene.obj$type        <- "rRNA"
gene.obj$entry       <- "gene"
gene.obj$placeholder <- "."
gene.obj$desc        <- paste0('gene_id "', gene.obj$name, '"; transcript_id ""; description ""; gbkey "Gene"; gene_biotype "protein_coding"; locus_tag "', gene.obj$name, '"; note "";')

gene.obj.order <- gene.obj[,c("chr","type","entry","start","stop", "placeholder", "strand", "placeholder","desc")]

# 2. create a "transcript" entry
transcript.obj             <- my.bed
colnames(transcript.obj)   <- c("chr","start","stop", "name","strand")
transcript.obj$type        <- "rRNA"
transcript.obj$entry       <- "transcript"
transcript.obj$placeholder <- "."
transcript.obj$desc        <- paste0('gene_id "', transcript.obj$name, '"; transcript_id "', transcript.obj$name, '"; description ""; gbkey "mRNA"; locus_tag "', transcript.obj$name, '"; orig_protein_id "', transcript.obj$name, '"; orig_transcript_id "', transcript.obj$name, '"; product ""; transcript_biotype "mRNA";')

transcript.obj.order <- transcript.obj[,c("chr","type","entry","start","stop", "placeholder", "strand", "placeholder","desc")]


# 3. create a "exon" entry
exon.obj             <- my.bed
colnames(exon.obj)   <- c("chr","start","stop", "name","strand")
exon.obj$type        <- "rRNA"
exon.obj$entry       <- "exon"
exon.obj$placeholder <- "."
exon.obj$desc        <- paste0('gene_id "', exon.obj$name, '"; transcript_id "', exon.obj$name, '"; locus_tag "', exon.obj$name, '"; orig_protein_id "', exon.obj$name, '"; orig_transcript_id "', exon.obj$name, '"; product ""; transcript_biotype "mRNA"; exon_number "1";')

exon.obj.order <- exon.obj[,c("chr","type","entry","start","stop", "placeholder", "strand", "placeholder","desc")]


# now interleave to make a gene, transcript, and exon line for each TE location
# create a rRNA only GTF file
rRNA_gtf <- gdata::interleave(gene.obj.order, transcript.obj.order, exon.obj.order)

# write tables for a TE-specific GTF and a whole genome GTF
write.table(rRNA_gtf, file = paste0("./Results/13_Make_Featurecounts_gtf/", Sys.Date(),"_rRNA_only.gtf"), sep = "\t", col.names = F, row.names = F, quote = F)


#2. Prepare cluster gtf

my.cluster.bed <- read.table('./Input/merged_cluster_coordinates.CLEAN.bed', sep = "\t", header = F)
my.cluster.bed$V5 <- "+"


# 1. create a "gene" entry
gene.obj             <- my.cluster.bed
colnames(gene.obj)   <- c("chr","start","stop", "name","strand")
gene.obj$type        <- "piRNA_cluster"
gene.obj$entry       <- "gene"
gene.obj$placeholder <- "."
gene.obj$desc        <- paste0('gene_id "', gene.obj$name, '"; transcript_id ""; description ""; gbkey "Gene"; gene_biotype "protein_coding"; locus_tag "', gene.obj$name, '"; note "";')

gene.obj.order <- gene.obj[,c("chr","type","entry","start","stop", "placeholder", "strand", "placeholder","desc")]

# 2. create a "transcript" entry
transcript.obj             <- my.cluster.bed
colnames(transcript.obj)   <- c("chr","start","stop", "name","strand")
transcript.obj$type        <- "piRNA_cluster"
transcript.obj$entry       <- "transcript"
transcript.obj$placeholder <- "."
transcript.obj$desc        <- paste0('gene_id "', transcript.obj$name, '"; transcript_id "', transcript.obj$name, '"; description ""; gbkey "mRNA"; locus_tag "', transcript.obj$name, '"; orig_protein_id "', transcript.obj$name, '"; orig_transcript_id "', transcript.obj$name, '"; product ""; transcript_biotype "mRNA";')

transcript.obj.order <- transcript.obj[,c("chr","type","entry","start","stop", "placeholder", "strand", "placeholder","desc")]


# 3. create a "exon" entry
exon.obj             <- my.cluster.bed
colnames(exon.obj)   <- c("chr","start","stop", "name","strand")
exon.obj$type        <- "piRNA_cluster"
exon.obj$entry       <- "exon"
exon.obj$placeholder <- "."
exon.obj$desc        <- paste0('gene_id "', exon.obj$name, '"; transcript_id "', exon.obj$name, '"; locus_tag "', exon.obj$name, '"; orig_protein_id "', exon.obj$name, '"; orig_transcript_id "', exon.obj$name, '"; product ""; transcript_biotype "mRNA"; exon_number "1";')

exon.obj.order <- exon.obj[,c("chr","type","entry","start","stop", "placeholder", "strand", "placeholder","desc")]

# now interleave to make a gene, transcript, and exon line for each TE location
# create a piRNA_cluster only GTF file
piRNA_cluster_gtf <- gdata::interleave(gene.obj.order, transcript.obj.order, exon.obj.order)

# write tables for a TE-specific GTF and a whole genome GTF
write.table(piRNA_cluster_gtf, file = paste0("./Results/13_Make_Featurecounts_gtf/", Sys.Date(),"_piRNA_cluster_only.gtf"), sep = "\t", col.names = F, row.names = F, quote = F)


#3. Prepare featurecounts-style TE gtf

#read in modulated Repeatmasker .out file; Code-checkers: unzip first (gzip file)
outfile <- read.table("./Input/preprocessed_gtf.fa.out", sep = "\t", header = F)

#remove simple repeats and low complexity regions
outfile <- outfile[!grepl("Simple", outfile$V11),]
outfile <- outfile[grepl("NotFur", outfile$V10),]

#change complementary ("C") to "-"
outfile$V9 <- gsub("C", "-",outfile$V9)

#keep only positinal info and names; strand
outfile <- outfile[,c(5:7,9:11)]

#separate into family types with unknowns/unclears being filled in when they throw up NAs
outfile_sep <- outfile %>% separate(V11, c("A", "B"),sep = "/")
outfile_sep$B <- ifelse(is.na(outfile_sep$B), outfile_sep$A, outfile_sep$B)

#remove bulky outfile for processing speed/memory concerns
rm(outfile)

# 1. create a "gene" entry
gene.obj             <- outfile_sep
colnames(gene.obj)   <- c("chr","start","stop","strand", "name", "group", "family")
gene.obj$type        <- "TE"
gene.obj$entry       <- "gene"
gene.obj$placeholder <- "."
gene.obj$desc        <- paste0('gene_id "', paste(gene.obj$name,  gene.obj$family, gene.obj$group, sep = ":"),
                               '"; transcript_id ""; description ""; gbkey "Gene"; gene_biotype "protein_coding"; locus_tag "',
                               paste(gene.obj$name,  gene.obj$family, gene.obj$group, sep = ":"), '"; note "";')

gene.obj.order <- gene.obj[,c("chr","type","entry","start","stop", "placeholder", "strand", "placeholder","desc")]

# 2. create a "transcript" entry
transcript.obj             <- outfile_sep
colnames(transcript.obj)   <- c("chr","start","stop","strand", "name", "group", "family")
transcript.obj$type        <- "TE"
transcript.obj$entry       <- "transcript"
transcript.obj$placeholder <- "."
transcript.obj$desc        <- paste0('gene_id "', paste(transcript.obj$name,  transcript.obj$family, transcript.obj$group, sep = ":"), '"; transcript_id "', paste(transcript.obj$name,  transcript.obj$family, transcript.obj$group, sep = ":"), '"; description ""; gbkey "mRNA"; locus_tag "', paste(transcript.obj$name,  transcript.obj$family, transcript.obj$group, sep = ":"), '"; orig_protein_id "', paste(transcript.obj$name,  transcript.obj$family, transcript.obj$group, sep = ":"), '"; orig_transcript_id "', paste(transcript.obj$name,  transcript.obj$family, transcript.obj$group, sep = ":"), '"; product ""; transcript_biotype "mRNA";')

transcript.obj.order <- transcript.obj[,c("chr","type","entry","start","stop", "placeholder", "strand", "placeholder","desc")]

# 3. create a "exon" entry
exon.obj             <- outfile_sep
colnames(exon.obj)   <- c("chr","start","stop","strand", "name", "group", "family")
exon.obj$type        <- "TE"
exon.obj$entry       <- "exon"
exon.obj$placeholder <- "."
exon.obj$desc        <- paste0('gene_id "', paste(exon.obj$name,  exon.obj$family, exon.obj$group, sep = ":"), '"; transcript_id "', paste(exon.obj$name,  exon.obj$family, exon.obj$group, sep = ":"), '"; locus_tag "', paste(exon.obj$name,  exon.obj$family, exon.obj$group, sep = ":"), '"; orig_protein_id "', paste(exon.obj$name,  exon.obj$family, exon.obj$group, sep = ":"), '"; orig_transcript_id "', paste(exon.obj$name,  exon.obj$family, exon.obj$group, sep = ":"), '"; product ""; transcript_biotype "mRNA"; exon_number "1";')

exon.obj.order <- exon.obj[,c("chr","type","entry","start","stop", "placeholder", "strand", "placeholder","desc")]

# now interleave to make a gene, transcript, and exon line for each TE location
# create a piRNA_cluster only GTF file
TE_cluster_gtf <- gdata::interleave(gene.obj.order, transcript.obj.order, exon.obj.order)

# write tables for a TE-specific GTF and a whole genome GTF
write.table(TE_cluster_gtf, file = paste0("./Results/13_Make_Featurecounts_gtf/", Sys.Date(),"_TE_only.gtf"), sep = "\t", col.names = F, row.names = F, quote = F)

#cat all of the gtfs together after running this script:
#1) GCA_014300015.1_MPIA_NFZ_2.0_genomic.gtf
#2) TE_cluster_gtf
#3) rRNA gtf
#4) piRNA_cluster_gtf
#save as Featurecounts_gtf.gtf

#######################
sink(file = paste("./Results/13_Make_Featurecounts_gtf/", Sys.Date(),"Generate_featurecounts_GTF","_session_Info.txt", sep =""))
sessionInfo()
sink()
```
