## Supplemental Code for "Dynamic regulation of gonadal transposon control across the lifespan of the naturally short-lived African turquoise killifish": 14_piRNA_counting.sh

```
#count_piRNAs.sh

#this script uses featurecounts in the Subread v2.0.2 package
#this script counts piRNAs fractionally allowing for multimappers

/project/bbenayou_34/teefy/RNA_seq/killifish_RNA_seq/Masked_Genome/subread-2.0.2-source/bin/featureCounts -O -M --fraction -a 2022-07-13_complete_piRNA_mapping_gtf.gtf -o piRNA_ftcts_output_no_MF2.txt  YM1.bam YM2.bam YM3.bam YM5.bam MM1.bam MM2.bam MM3.bam MM5.bam OM1.bam OM3.bam OM4.bam OM5.bam YF1.bam YF2.bam YF3.bam YF4.bam YF5.bam MF1.bam MF2.bam MF3.bam MF4.bam MF5.bam OF1.bam OF3.bam OF4.bam OF5.bam
```
