## Supplemental Code for "Dynamic regulation of gonadal transposon control across the lifespan of the naturally short-lived African turquoise killifish": 15_make_TE_only_bowtie_index.sh

```
#make_TE_only_bowtie_index.sh

# this script generates a TE-only bowtie index that will be used
# for te locus level ping-pong analysis

# this script uses bowtie v1.2.3

bowtie-build Nothobranchius_furzeri_TEs.fa ./killi_TE_bowtie_ref
```
