## Supplemental Code for "Dynamic regulation of gonadal transposon control across the lifespan of the naturally short-lived African turquoise killifish": 17_convert_bam_to_bed_pingpong.sh

```
#convert_bam_to_bed_pingpong.sh

# this script uses bedtools v2.27.1

#Now we need to convert to bed files for subsequent processing steps

for f in $(find "." -name '*.bam')
        do
    f2=$(basename "${f}" | sed 's/\.bam/\.bed/g');

bamToBed -i $f > $f2

done
```
