## Supplemental Code for "Dynamic regulation of gonadal transposon control across the lifespan of the naturally short-lived African turquoise killifish": 18_locus_level_z_score_generation.sh

```
#locus_level_z_score_generation.sh

#This script takes mapped piRNA bedfiles, separates them into two different bed files pertaining 
#to each strand (sense/antisense), then finds all the overlaps between each 5’ ends on opposite strands.

#Then it retains only the TE name and overlap length

#Then it generates a histogram of the overlap lengths for each consensus TE.

#this script uses bedtools v2.27.1

for f in $(find "." -name '*.bed')
        do
        pos_bed=$(basename "${f}" | sed 's/\.bed/positive\.bed/g');
        neg_bed=$(basename "${f}" | sed 's/\.bed/negative\.bed/g');
        pos_bed_sorted=$(basename "${f}" | sed 's/\.bed/positive_sorted\.bed/g');
        neg_bed_sorted=$(basename "${f}" | sed 's/\.bed/negative_sorted\.bed/g');
        outfile=$(basename "${f}" | sed 's/\.bed/_overlaps\.txt/g');
        of_hist=$(basename "${f}" | sed 's/\.bed/_histos\.txt/g');
        of_hist_red=$(basename "${f}" | sed 's/\.bed/_histos_reduced\.txt/g');

awk '$6 == "+" { print $0 }' $f > $pos_bed
awk '$6 == "-" { print $0 }' $f > $neg_bed

sort -k1,1 -k2,2n $pos_bed > $pos_bed_sorted
sort -k1,1 -k2,2n $neg_bed > $neg_bed_sorted

rm $pos_bed
rm $neg_bed

bedtools intersect -a $pos_bed_sorted -b $neg_bed_sorted -sorted | awk 'BEGIN { OFS = "\t" } { $7 = $3 - $2 } 1' | awk '{ print $1, $7 }' > $outfile

rm $pos_bed_sorted
rm $neg_bed_sorted

awk 'NR == FNR { ++ctr[$1,$2]; next } { print $0 "\t" ctr[$1,$2]; }' $outfile $outfile > $of_hist

rm $outfile

awk '!a[$0]++' $of_hist > $of_hist_red

rm $of_hist

done
```
