## Supplemental Code for "Dynamic regulation of gonadal transposon control across the lifespan of the naturally short-lived African turquoise killifish": 19_ppMeter_map_to_TEs.sh

```
#ppMeter_map_to_TEs.sh

#this script uses sRNAmapper.pl from proTRAC v2.4.2
#uses perl v5.30.1

for f in $(find "." -name '/path/to/collaped/reads/*_piRNA.collapsed.no-dust')
do

f1=$(basename "${f}");

perl sRNAmapper.pl -input $f1 -genome Nothobranchius_furzeri_TEs.fa -alignments best

done
```
