## Supplemental Code for "Dynamic regulation of gonadal transposon control across the lifespan of the naturally short-lived African turquoise killifish": 20_looped_pingpongmeter.sh

```
#looped_pingpongmeter.sh

#PPmeter takes the *no-dust.map files generated by proTRAC as input.
#Uses .map files that were mapped to consensus TEs only

#this script uses PPmeter_0.4.pl from the ProTRAC v2.42 software suite
#this script uses perl v5.30.1

for f in $(find "." -name '*_piRNA.collapsed.no-dust.map')
do

    f1=$(basename "${f}");

        perl PPmeter_0.4.pl -i $f1 -f ELAND3

done
```
