## Supplemental Code for "Dynamic regulation of gonadal transposon control across the lifespan of the naturally short-lived African turquoise killifish": README_piRNA_preprocessing.txt

########################################################
			README -- 2_piRNA_preprocessing
########################################################

	General Pre-Processing
	######################
	1.   sRNA reads trimmed with trim_galore (1_small_RNA_trimming.sh)
	2.   piRNA isolation by size selection (2_size_split_sRNAs.sh)
	3.   Obtain piRNA length data for general characterization (3_piRNA_length_generation.sh)
	4.   Prepare piRNA data for logo plot generation (4_preprocess_logo_plots.sh)
	
	piRNA Cluster Pre-Processing
	######################
	5.   Convert piRNAs from fastq to fasta format for proTRAC (5_piRNA_fq_to_fa.sh)
	6.   Identify piRNA cluster with proTRAC (6_piRNA_cluster_identification_protrac.sh)
	7.   Consolidate piRNA clusters (7_modified_merge.pl)
	8.   Extract piRNA clusters from genome (8_get_cluster_fasta.sh)
	9a.  Obtain TE repeat content in clusters (R) (9a_TE_repeat_content_in_clusters.R)
	9b.  Obtain TE repeat content in clusters (bash) (9b_te_clusters_bedtools_intersect.sh) 

	piRNA DGE Pre-Processing
	######################	
	10.  Index softmasked genome with bowtie (10_bowtie_genomic_index.sh)
	11.  Map piRNAs to the softmasked genome using bowtie (11_piRNA_mapping_to_genome.sh)
	12.  Process rRNA sequence into format for gtf creation (12_Prepare_rRNA_for_gtf.txt)
	13.  Generate a gtf for featurecounts with rRNA, piRNA cluster, and TE sequences included (13_Make_Featurecounts_gtf.R)
	14.  Count piRNAs using featurecounts (14_piRNA_counting.sh)
	
	Z-Score Pre-Processing
	######################	
	15.  Generate index for consensus TE sequences with bowtie (15_make_TE_only_bowtie_index.sh)
	16.  Map piRNAs to consensus TE reference with bowtie (16_map_piRNAs_to_TE_index.sh)
	17.  Convert consensus TE-mapped piRNA bam files to bed files for locus-level Z-Score Analysis (17_convert_bam_to_bed_pingpong.sh)
	18.  Generate consensus TE-level Z-scores (18_locus_level_z_score_generation.sh)

	PPmeter Pre-Processing
	######################
	19.  Map piRNAs to consensus TE reference using proTRAC sRNAmapper.pl (19_ppMeter_map_to_TEs.sh)
	20.  Run PPmeter to generate ping-pong matrices (20_looped_pingpongmeter.sh)
