## Supplemental Code for "Dynamic regulation of gonadal transposon control across the lifespan of the naturally short-lived African turquoise killifish": 3a_BLAST_preprocessing.txt

#BLAST Scripts for GO

#This script uses local ncbi-blast v2.13.0

#1) Download Homo sapiens Protein Sequence (FASTA) from Ensembl
# Sequences were downloaded August 9, 2021, corresponding to Ensembl release 104 from

http://uswest.ensembl.org/info/data/ftp/index.html/
# Ensembl 104 now archived at: http://may2021.archive.ensembl.org/biomart/martview/08e3f73098b66635c144993bfd6ab30d

To download human protein sequences, perform the following steps:

	a) Select to "BioMart" from the menu at the top of the webpage
	b) CHOOSE DATABASE: Select "Ensembl Genes [version]"
			-Note: this publication used Ensembl release 104
	c) CHOOSE DATASET: Select "Human genes (GRCh38.p13)"
	d) Select "Attributes" on the side menu
	e) Select the "Sequences" option
	f) Under the newly populated "SEQUENCES" menu, ensure that "Peptide" is selected
	g) Select "Results" from the top left corner menu
	h) Ensure that protein sequences are present in FASTA format 
		ex. >ENSG00000001617|ENSG00000001617.12|ENST00000002829|ENST00000002829.8
			MLVAGLLLWASLL...*
	i) Export file as a "Compressed file (.gz)" in FASTA format
	j) Unzip file locally and save this file as "human_prot.fasta".
	
#2) Generate a local BLAST database of the Homo sapiens Protein Sequences

makeblastdb -in human_prot.fasta -dbtype prot -name human_prot_db 

#3) BLAST Killifish Protein Sequences against the Human Protein Database

blastp -db human_prot_db -query GCA_014300015.1_MPIA_NFZ_2.0_protein.faa -outfmt 7 -max_target_seqs 1 -out prot_hits_human_killi.txt

#4) Use a perl one-liner to parse BLAST output to obtain (i) only hits that have E-value < 1e-3 and (ii) unique lines

cat prot_hits_human_killi.txt | grep -v '#' | perl -lane 'next if $F[10] > 1e-3; @human = split(/\|/,$F[1]); print "$F[0]\t$human[0]"' | sort -u > Parsed_human_to_killi_BLAST_1e-3.txt


#5) Parse the genome-associated gtf file, "GCA_014300015.1_MPIA_NFZ_2.0_genomic.gtf" to extract geneID/ProteinID conversion

 Use R script 3b_GTF_preprocessing.R to generate a correspondence table Parsed_GCA_014300015.1_MPIA_NFZ_2.0_Gene_to_Protein_ID_conversion.txt

#6) Download a correspondence table from ENSEMBL GeneID to Gene Symbol for humans to enable gene symbol mapping
   [Steps will be similar to 1]

	a) Select to "BioMart" from the menu at the top of the webpage
	b) CHOOSE DATABASE: Select "Ensembl Genes [version]"
	c) CHOOSE DATASET: Select "Human genes (GRCh38.p13)"
	d) Select "Attributes" on the side menu
	e) Select the "Features" option
	f) select "Gene stable ID" and "Gene name"
	g) Select "Results" from the top left corner menu
	h) Select "Unique results only" for tsv output
	j) Save file as "Ens104_GeneID_GeneSymbol_HUMAN_mart_export.txt".


##### Files to use in GO analysis:
	- Parsed_GCA_014300015.1_MPIA_NFZ_2.0_Gene_to_Protein_ID_conversion.txt
	- Parsed_human_to_killi_BLAST_1e-3.txt
	- Ens104_GeneID_GeneSymbol_HUMAN_mart_export.txt
