## Supplemental Code for "Dynamic regulation of gonadal transposon control across the lifespan of the naturally short-lived African turquoise killifish": 3b_GTF_preprocessing.R

```
setwd("/Users/berenice/Dropbox/Manuscripts_and_Publications/2022/2022_Bryan_PIWI_manuscript/CODE/3_Differential_Gene_Expression_Analysis")

# R version 4.1.2 (2021-11-01)

library(rtracklayer) # rtracklayer_1.54.0

# Parse GTF table to get gene/protein ID correspondence

# since killifish genomic data is not in annotation hub, need to import from gtf annotation file
gtf <- rtracklayer::import('/Users/berenice/Dropbox/Manuscripts_and_Publications/2022/2022_Bryan_PIWI_manuscript/CODE/FINAL_REPO/GCA_014300015.1_MPIA_NFZ_2.0_genomic.gtf')
gtf <- as.data.frame(gtf)

# clean up and format annotation data
gene_metadata <- unique(gtf[,c("gene_id","protein_id")])

# Keep only entries with protein_id
gene_metadata <- gene_metadata[!is.na(gene_metadata$protein_id),]

# export as text file
write.table(gene_metadata, file = "./Results/3_BLAST_and_annot_Parsing/Parsed_GCA_014300015.1_MPIA_NFZ_2.0_Gene_to_Protein_ID_conversion.txt", sep = "\t", quote = F, col.names = T, row.names = F)

##########################################################
sink(file = paste("./Results/3_BLAST_and_annot_Parsing/", Sys.Date(),"_gtf_parsing_session_Info.txt", sep =""))
sessionInfo()
sink()
```
