## Supplemental Code for "Dynamic regulation of gonadal transposon control across the lifespan of the naturally short-lived African turquoise killifish": 4a_Functions_for_GO_Enrich.R

```
################################################################
# DEBUG
# my.list       <- Ovary_Down_MAge
# my.conv.table <- conversion_table

convert_k2h <- function(my.list, my.conv.table) {
  
  # get the human homologs (column 2) corresponding to killi gene list ("gene_id")
  list.conv <- unique(my.conv.table[my.conv.table$gene_id %in% my.list,2])
  
  return(list.conv)
}
################################################################


################################################################
# DEBUG
# fg.list  <- Ovary_Down_MAge
# universe <- ovary.universe
# gsetColl <- gsc
# my.ontology <- "BP"

run_GO <- function(fg.list, universe, gsetColl, my.ontology, fdr){
  
  # specify GOstats parameters
  params <- GSEAGOHyperGParams(name              = paste0("GO_",my.ontology), 
                               geneSetCollection = gsetColl, 
                               geneIds           = fg.list$V1, 
                               universeGeneIds   = universe, 
                               ontology          = my.ontology,
                               pvalueCutoff      = 1, # 1 will get all terms, and then we can filter later
                               conditional       = F, # To consider GO DAG structure or not. 
                               testDirection     = "over") # Default hyper geometric test gives both over and under enriched terms. I am specifying the direction by "over". Fo depleted terms use under.
  
  # call hyperGTest to perform enrichment test
  Over <- hyperGTest(params)
  
  # calculate enrichment and add it to data frame.
  # Relative enrichment factor (E-value) for a GO term = (count/ DE )/(size/universe)
  enrichment = (summary(Over)[5]$Count / length(fg.list$V1)) / (summary(Over)[6]$Size / length(universe))
  
  # create a new data frame
  SummaryOver = data.frame(summary(Over), enrichment)
  
  # Filter the summary of OVER with size of the term, at least 2 genes for a go term
  # nonsensical to run enrichment if overlap less than that
  SummaryOver <- SummaryOver[SummaryOver$Count >= 2,]
  
  # adjust p value for multiple correction
  SummaryOver$padj <- p.adjust(SummaryOver$Pvalue, "BH")
  
  return(SummaryOver[SummaryOver$padj < fdr,])
}
################################################################


################################################################
#define function for plotting bubble data
# DEBUG
# my.color.vector.age = colorRampPalette(c("lavender","darkviolet"))(5)
# GO_res              = Ovary_Down_MAge_GO
# sex                 = "F"

bubble_plot <- function(GO_res, sex, cond, my.color.vector.age = colorRampPalette(c("lavender","darkviolet"))(5)){
  
  # generate -log10(FDR) 
  GO_res$minlog10fdr <- -log10(GO_res$padj)
  
  # get top 10 most enriched
  GO_res$sex  <- sex
  GO_res$Name <- paste0(GO_res[,grep("GO",colnames(GO_res))]," ",GO_res$Term)
  
  # top 10 most significant
  GO_res.filt <- GO_res %>%                                      
    arrange(padj) %>% 
    slice(1:10)
  
  # set up ranges
  my.max <- max(GO_res.filt$enrichment)
  my.values <- c(0,0.25*my.max,0.5*my.max,0.75*my.max,my.max)
  my.scaled <- rescale(my.values, to = c(0, 1))
  
  
  # create and return bubble plot
  bub_plot <- ggplot(GO_res.filt,aes(x = sex,y = reorder(Name, enrichment),colour = enrichment, size = minlog10fdr)) +
    theme_bw() + geom_point(shape = 16) + ggtitle(paste0("GO ", cond) )+
    scale_colour_gradientn(colours = my.color.vector.age, 
                           na.value = "grey50", guide = "colourbar", values = my.scaled, limits = c(0,ceiling(my.max))) +
    scale_size_continuous(limits = c(1, max(ceiling(GO_res.filt$minlog10fdr)) ))
    
    
  return(bub_plot)
}
################################################################
```
