## Supplemental Code for "Dynamic regulation of gonadal transposon control across the lifespan of the naturally short-lived African turquoise killifish": 4b_GO_Enrichment_Analysis.R

```
# setwd("/Users/bryanteefy/Dropbox/PIWI/CODE/3_Differential_Gene_Expression_Analysis")
setwd("/Users/berenice/Dropbox/Manuscripts_and_Publications/2022/2022_Bryan_PIWI_manuscript/CODE/3_Differential_Gene_Expression_Analysis")

# GO Analysis
# Original script written by Param Priya Singh
# Modified by Bryan Teefy & Bérénice Benayoun

# R version 4.1.2 (2021-11-01)

# load required libraries
library(GOstats)   # GOstats_2.60.0
library(biomaRt)   # biomaRt_2.50.3
library(GSEABase)  # GSEABase_1.56.0
library(scales)    # scales_1.2.0
library(ggplot2)   # ggplot2_3.3.6
library(dplyr)     # dplyr_1.0.9
library(ggplotify) # ggplotify_0.1.0
library(patchwork) # patchwork_1.1.1 
library(pheatmap)  # pheatmap_1.0.12

# load functions to run enrichment information
source('4a_Functions_for_GO_Enrich.R')

#################################################
# 1. prepare data for geneset analysis and set parameters

# import Killi CDS ID to human Ensembl Gene ID conversion table
blast_table <- read.table("./Input/Parsed_human_to_killi_BLAST_1e-3.txt", header = F, sep = "\t")
colnames(blast_table) <- c("killi_prot", "ensembl_gene_id")

# import human name to Ensembl ID conversion table
name_table <- read.csv("./Input/Ens104_GeneID_GeneSymbol_HUMAN_mart_export.txt", header = T, sep = "\t")
colnames(name_table) <- c("ensembl_gene_id", "Human_Gene_Symbol")

# import Gene ID to Killi CDS ID conversion table
prot_table <- read.csv("./Input/Parsed_GCA_014300015.1_MPIA_NFZ_2.0_Gene_to_Protein_ID_conversion.txt", header = T, sep = "\t")

# merge tables to create conversion table
conversion_table.1   <- merge(blast_table       , name_table, by    = "ensembl_gene_id")
conversion_table.2   <- merge(conversion_table.1, prot_table, by.x  = "killi_prot"     , by.y = "protein_id")

# Subset columns to retain only Killifish gene and Human Gene Symbol
# Since this annotation version can have more than 1 protein per gene, deduplicate
conversion_table     <- unique(conversion_table.2[,c("gene_id","Human_Gene_Symbol")])

# Output homology table for manuscript
write.table(unique(conversion_table.2[,c("killi_prot", "gene_id","ensembl_gene_id" , "Human_Gene_Symbol")]), file = paste0("./Results/", Sys.Date(),"_Killi_human_homology_table.txt"), sep = "\t", quote = F, row.names = F)

##################################################
# 2. Prepare gene set objects

#### get GO terms from ensembl 104 (for reproducibility since it was used for BLAST)
ensembl104 <- useEnsembl(biomart = 'genes', 
                         dataset = 'hsapiens_gene_ensembl',
                         version = 104)

# Obtain the GO terms associated with each human gene (including GO evidence domain)
h.genes2map <- unique(conversion_table.2$ensembl_gene_id)
length(h.genes2map) # 13344

# To avoid server/curl timeout, run in max 1000 gene slices
slices.st <- c(seq(1,length(h.genes2map),1000),length(h.genes2map))
GO_list.slices <- vector(mode = "list", length(slices.st)-1)

# this will run for ~15-30min
for (i in 1:length(GO_list.slices)) {
  
  GO_list.slices[[i]] <- getBM(filters    = "ensembl_gene_id", 
                               attributes = c("ensembl_gene_id","hgnc_symbol", "go_id", "namespace_1003"),   # namespace_1003 is GO domain [BP, CC, MF]
                               values     = h.genes2map[slices.st[i]:slices.st[i+1]], # always 1 overlap between slices, but repeated lines will be filtered out
                               uniqueRows = TRUE,
                               mart       = ensembl104)
}

# merge slices for downstream processing into a single object
GO_list <- GO_list.slices[[1]] # initialize
for (i in 2:length(GO_list.slices)) {
  GO_list <- unique(rbind(GO_list,GO_list.slices[[i]])) # merge and remove duplicate lines
}

# remove rows without a GO ID or a gene symbol
GO_list <- GO_list[GO_list$go_id != "",]
GO_list <- GO_list[GO_list$hgnc_symbol != "",]

save(GO_list, file = paste0("./Results/", Sys.Date(),"_BioMart_Ens104_HUMAN_GO_list_with_Domain.Rdata"))
# load("./Results/4_GO_enrichment/2022-07-13_BioMart_Ens104_HUMAN_GO_list_with_Domain.Rdata") # time save

# merge with the conversion table to get a GO table for killifish
GO_conv <- merge(GO_list[,-1], conversion_table, by.x = "hgnc_symbol",  by.y = "Human_Gene_Symbol")
GO_conv <- unique(GO_conv) # remove duplicated rows (if any)

# Parse into required 3 columns
goframeData <- data.frame("go_id"    = GO_conv$go_id, 
                          "Evidence" = rep("ISO", nrow(GO_conv)),
                          "gene_id"  = GO_conv$gene_id)

# put your data into a GOFrame object
goFrame <- GOFrame(goframeData, organism = "Killifish")
head(goframeData)
#         go_id Evidence      gene_id
# 1 GO:0003727      ISO G4P62_011648
# 2 GO:0016556      ISO G4P62_011648
# 3 GO:0005654      ISO G4P62_011648
# 4 GO:0016554      ISO G4P62_011648
# 5 GO:0003676      ISO G4P62_011648
# 6 GO:0005783      ISO G4P62_011648

# cast this object to a GOAllFrame object will tap into the GO.db package and populate this object with the implicated GO2All mappings for you
goAllFrame <- GOAllFrame(goFrame)

# generate geneSetCollection objects
gsc <- GeneSetCollection(goAllFrame, setType = GOCollection())

# FDR threshold for significance
fdr_thrs = 0.05

#################################################
# 3. Read in data from each groups from LRT gene analysis at FDR 1e-6 (and corresponding universe)

# Female Data
Ovary_Down_MAge <-  read.table("./Results/1_Ovary_DGE/2022-07-13_Ovary_LRT_Female_AGING_Group1_DGE_Genes_FDR1e-6.txt", skip = 1) # first line is "x"
Ovary_Up_MAge   <-  read.table("./Results/1_Ovary_DGE/2022-07-13_Ovary_LRT_Female_AGING_Group2_DGE_Genes_FDR1e-6.txt", skip = 1) # first line is "x"
Ovary_Down_Age  <-  read.table("./Results/1_Ovary_DGE/2022-07-13_Ovary_LRT_Female_AGING_Group3_DGE_Genes_FDR1e-6.txt", skip = 1) # first line is "x"
Ovary_Up_Age    <-  read.table("./Results/1_Ovary_DGE/2022-07-13_Ovary_LRT_Female_AGING_Group4_DGE_Genes_FDR1e-6.txt", skip = 1) # first line is "x"

# Male Data
Testes_Up_MAge   <-  read.table("./Results/2_Testis_DGE/2022-07-13_Testes_LRTMale_AGING_Group1_DGE_Genes_FDR1e-6.txt", skip = 1) # first line is "x"
Testes_Down_MAge <-  read.table("./Results/2_Testis_DGE/2022-07-13_Testes_LRTMale_AGING_Group2_DGE_Genes_FDR1e-6.txt", skip = 1) # first line is "x"
Testes_Down_Age  <-  read.table("./Results/2_Testis_DGE/2022-07-13_Testes_LRTMale_AGING_Group3_DGE_Genes_FDR1e-6.txt", skip = 1) # first line is "x"
# Testis up with Age has only 2 genes: nonsensical to run enrichment analysis
# Testes_Up_Age    <-  read.table("./Results/2_Testis_DGE/2022-07-13_Testes_LRTMale_AGING_Group4_DGE_Genes_FDR1e-6.txt", skip = 1) # first line is "x"

# get gene universe for ovaries and testes (all genes with LRT p-value in tissue)
ovary.universe   <- read.table("./Results/1_Ovary_DGE/2022-07-13_Ovary_LRT_Ovarian_LRT_DGE_Table_GeneONLY.txt", header = T, sep = "\t")$gene # 18660 genes
testis.universe  <- read.table("./Results/2_Testis_DGE/2022-07-13_Testes_LRT_Testicular_LRT_DGE_Table_GeneONLY.txt", header = T, sep = "\t")$gene # 19658 genes


#################################################
# 4. Run Functional enrichment analyses with GO (hypergeometric enrichment)

# Testis Up with Age has only 2 genes: nonsensical to run enrichment analysis

####### GO BP
# run GO on ovaries DE genes
Ovary_Down_MAge_GOBP <- run_GO(Ovary_Down_MAge  , ovary.universe , gsc, "BP", fdr_thrs) # 45 pathways
Ovary_Up_MAge_GOBP   <- run_GO(Ovary_Up_MAge    , ovary.universe , gsc, "BP", fdr_thrs) # 81 pathways
Ovary_Down_Age_GOBP  <- run_GO(Ovary_Down_Age   , ovary.universe , gsc, "BP", fdr_thrs) # 213 pathways
Ovary_Up_Age_GOBP    <- run_GO(Ovary_Up_Age     , ovary.universe , gsc, "BP", fdr_thrs) # 26 pathways

# run GO on testes  DE genes
Testes_Down_MAge_GOBP <- run_GO(Testes_Down_MAge, testis.universe, gsc, "BP", fdr_thrs) # 184 pathways
Testes_Up_MAge_GOBP   <- run_GO(Testes_Up_MAge  , testis.universe, gsc, "BP", fdr_thrs) # 51 pathways
Testes_Down_Age_GOBP  <- run_GO(Testes_Down_Age , testis.universe, gsc, "BP", fdr_thrs) # 177 pathways

# write and export GO results
write.table(Ovary_Down_MAge_GOBP, file = paste0("./Results/4_GO_enrichment/Individual_results/", Sys.Date(), "_GOstats_Ovary_Down_MAge_GOBP_FDR5_"  , nrow(Ovary_Down_MAge_GOBP ) , "_signif.txt"   ), sep = "\t" , row.names = F, col.names = T, quote = F)
write.table(Ovary_Up_MAge_GOBP  , file = paste0("./Results/4_GO_enrichment/Individual_results/", Sys.Date(), "_GOstats_Ovary_Up_MAge_GOBP_FDR5_"    , nrow(Ovary_Up_MAge_GOBP   ) , "_signif.txt"   ), sep = "\t" , row.names = F, col.names = T, quote = F)
write.table(Ovary_Down_Age_GOBP , file = paste0("./Results/4_GO_enrichment/Individual_results/", Sys.Date(), "_GOstats_Ovary_Down_Age_GOBP_FDR5_"   , nrow(Ovary_Down_Age_GOBP  ) , "_signif.txt"   ), sep = "\t" , row.names = F, col.names = T, quote = F)
write.table(Ovary_Up_Age_GOBP   , file = paste0("./Results/4_GO_enrichment/Individual_results/", Sys.Date(), "_GOstats_Ovary_Up_Age_GOBP_FDR5_"     , nrow(Ovary_Up_Age_GOBP    ) , "_signif.txt"   ), sep = "\t" , row.names = F, col.names = T, quote = F)

write.table(Testes_Down_MAge_GOBP, file = paste0("./Results/4_GO_enrichment/Individual_results/", Sys.Date(), "_GOstats_Testes_Down_MAge_GOBP_FDR5_", nrow(Testes_Down_MAge_GOBP ), "_signif.txt"  ), sep = "\t" , row.names = F, col.names = T, quote = F)
write.table(Testes_Up_MAge_GOBP  , file = paste0("./Results/4_GO_enrichment/Individual_results/", Sys.Date(), "_GOstats_Testes_Up_MAge_GOBP_FDR5_"  , nrow(Testes_Up_MAge_GOBP   ), "_signif.txt"  ), sep = "\t" , row.names = F, col.names = T, quote = F)
write.table(Testes_Down_Age_GOBP , file = paste0("./Results/4_GO_enrichment/Individual_results/", Sys.Date(), "_GOstats_Testes_Down_Age_GOBP_FDR5_" , nrow(Testes_Down_Age_GOBP  ), "_signif.txt"  ), sep = "\t" , row.names = F, col.names = T, quote = F)


####### GO MF
# run GO on ovaries DE genes
Ovary_Down_MAge_GOMF <- run_GO(Ovary_Down_MAge  , ovary.universe , gsc, "MF", fdr_thrs) # 2 pathways
Ovary_Up_MAge_GOMF   <- run_GO(Ovary_Up_MAge    , ovary.universe , gsc, "MF", fdr_thrs) # 43 pathways
Ovary_Down_Age_GOMF  <- run_GO(Ovary_Down_Age   , ovary.universe , gsc, "MF", fdr_thrs) # 29 pathways
Ovary_Up_Age_GOMF    <- run_GO(Ovary_Up_Age     , ovary.universe , gsc, "MF", fdr_thrs) # 10 pathways

# run GO on testes  DE genes
Testes_Down_MAge_GOMF <- run_GO(Testes_Down_MAge, testis.universe, gsc, "MF", fdr_thrs) # 17 pathways
Testes_Up_MAge_GOMF   <- run_GO(Testes_Up_MAge  , testis.universe, gsc, "MF", fdr_thrs) # 7 pathways
Testes_Down_Age_GOMF  <- run_GO(Testes_Down_Age , testis.universe, gsc, "MF", fdr_thrs) # 8 pathways

# write and export GO results
write.table(Ovary_Down_MAge_GOMF, file = paste0("./Results/4_GO_enrichment/Individual_results/", Sys.Date(), "_GOstats_Ovary_Down_MAge_GOMF_FDR5_"  , nrow(Ovary_Down_MAge_GOMF)  , "_signif.txt"   ), sep = "\t" , row.names = F, col.names = T, quote = F)
write.table(Ovary_Up_MAge_GOMF  , file = paste0("./Results/4_GO_enrichment/Individual_results/", Sys.Date(), "_GOstats_Ovary_Up_MAge_GOMF_FDR5_"    , nrow(Ovary_Up_MAge_GOMF  )  , "_signif.txt"   ), sep = "\t" , row.names = F, col.names = T, quote = F)
write.table(Ovary_Down_Age_GOMF , file = paste0("./Results/4_GO_enrichment/Individual_results/", Sys.Date(), "_GOstats_Ovary_Down_Age_GOMF_FDR5_"   , nrow(Ovary_Down_Age_GOMF )  , "_signif.txt"   ), sep = "\t" , row.names = F, col.names = T, quote = F)
write.table(Ovary_Up_Age_GOMF   , file = paste0("./Results/4_GO_enrichment/Individual_results/", Sys.Date(), "_GOstats_Ovary_Up_Age_GOMF_FDR5_"     , nrow(Ovary_Up_Age_GOMF   )  , "_signif.txt"   ), sep = "\t" , row.names = F, col.names = T, quote = F)

write.table(Testes_Down_MAge_GOMF, file = paste0("./Results/4_GO_enrichment/Individual_results/", Sys.Date(), "_GOstats_Testes_Down_MAge_GOMF_FDR5_", nrow(Testes_Down_MAge_GOMF ), "_signif.txt"  ), sep = "\t" , row.names = F, col.names = T, quote = F)
write.table(Testes_Up_MAge_GOMF  , file = paste0("./Results/4_GO_enrichment/Individual_results/", Sys.Date(), "_GOstats_Testes_Up_MAge_GOMF_FDR5_"  , nrow(Testes_Up_MAge_GOMF   ), "_signif.txt"  ), sep = "\t" , row.names = F, col.names = T, quote = F)
write.table(Testes_Down_Age_GOMF , file = paste0("./Results/4_GO_enrichment/Individual_results/", Sys.Date(), "_GOstats_Testes_Down_Age_GOMF_FDR5_" , nrow(Testes_Down_Age_GOMF  ), "_signif.txt"  ), sep = "\t" , row.names = F, col.names = T, quote = F)


####### GO CC
# run GO on ovaries DE genes
Ovary_Down_MAge_GOCC <- run_GO(Ovary_Down_MAge  , ovary.universe , gsc, "CC", fdr_thrs) # 7 pathways
Ovary_Up_MAge_GOCC   <- run_GO(Ovary_Up_MAge    , ovary.universe , gsc, "CC", fdr_thrs) # 35 pathways
Ovary_Down_Age_GOCC  <- run_GO(Ovary_Down_Age   , ovary.universe , gsc, "CC", fdr_thrs) # 45 pathways
Ovary_Up_Age_GOCC    <- run_GO(Ovary_Up_Age     , ovary.universe , gsc, "CC", fdr_thrs) #### 0 sig pathway

# run GO on testes  DE genes
Testes_Down_MAge_GOCC <- run_GO(Testes_Down_MAge, testis.universe, gsc, "CC", fdr_thrs) # 27 pathways
Testes_Up_MAge_GOCC   <- run_GO(Testes_Up_MAge  , testis.universe, gsc, "CC", fdr_thrs) # 52 pathways
Testes_Down_Age_GOCC  <- run_GO(Testes_Down_Age , testis.universe, gsc, "CC", fdr_thrs) # 20 pathways

# write and export GO results
write.table(Ovary_Down_MAge_GOCC, file = paste0("./Results/4_GO_enrichment/Individual_results/", Sys.Date(), "_GOstats_Ovary_Down_MAge_GOCC_FDR5_"  , nrow(Ovary_Down_MAge_GOCC)  , "_signif.txt"   ), sep = "\t" , row.names = F, col.names = T, quote = F)
write.table(Ovary_Up_MAge_GOCC  , file = paste0("./Results/4_GO_enrichment/Individual_results/", Sys.Date(), "_GOstats_Ovary_Up_MAge_GOCC_FDR5_"    , nrow(Ovary_Up_MAge_GOCC  )  , "_signif.txt"   ), sep = "\t" , row.names = F, col.names = T, quote = F)
write.table(Ovary_Down_Age_GOCC , file = paste0("./Results/4_GO_enrichment/Individual_results/", Sys.Date(), "_GOstats_Ovary_Down_Age_GOCC_FDR5_"   , nrow(Ovary_Down_Age_GOCC )  , "_signif.txt"   ), sep = "\t" , row.names = F, col.names = T, quote = F)
write.table(Ovary_Up_Age_GOCC   , file = paste0("./Results/4_GO_enrichment/Individual_results/", Sys.Date(), "_GOstats_Ovary_Up_Age_GOCC_FDR5_"     , nrow(Ovary_Up_Age_GOCC   )  , "_signif.txt"   ), sep = "\t" , row.names = F, col.names = T, quote = F)

write.table(Testes_Down_MAge_GOCC, file = paste0("./Results/4_GO_enrichment/Individual_results/", Sys.Date(), "_GOstats_Testes_Down_MAge_GOCC_FDR5_", nrow(Testes_Down_MAge_GOCC ), "_signif.txt"  ), sep = "\t" , row.names = F, col.names = T, quote = F)
write.table(Testes_Up_MAge_GOCC  , file = paste0("./Results/4_GO_enrichment/Individual_results/", Sys.Date(), "_GOstats_Testes_Up_MAge_GOCC_FDR5_"  , nrow(Testes_Up_MAge_GOCC   ), "_signif.txt"  ), sep = "\t" , row.names = F, col.names = T, quote = F)
write.table(Testes_Down_Age_GOCC , file = paste0("./Results/4_GO_enrichment/Individual_results/", Sys.Date(), "_GOstats_Testes_Down_Age_GOCC_FDR5_" , nrow(Testes_Down_Age_GOCC  ), "_signif.txt"  ), sep = "\t" , row.names = F, col.names = T, quote = F)


#################################################
# 5. create summary tables for supplementary table

# for simplicity, we are using 1 -> c, 2 -> b, 3 -> c and 4 -> d to export for supplementary tables

## BP
ovary.GOBP          <- rbind(Ovary_Down_MAge_GOBP,
                             Ovary_Up_MAge_GOBP  ,
                             Ovary_Down_Age_GOBP ,
                             Ovary_Up_Age_GOBP   )
ovary.GOBP$cluster  <- c(rep(1,nrow(Ovary_Down_MAge_GOBP)),
                         rep(2,nrow(Ovary_Up_MAge_GOBP  )),
                         rep(3,nrow(Ovary_Down_Age_GOBP )),
                         rep(4,nrow(Ovary_Up_Age_GOBP   )))

testis.GOBP         <- rbind(Testes_Down_MAge_GOBP ,
                             Testes_Up_MAge_GOBP   ,
                             Testes_Down_Age_GOBP  )
testis.GOBP$cluster <- c(rep(1,nrow(Testes_Down_MAge_GOBP )),
                         rep(2,nrow(Testes_Up_MAge_GOBP   )),
                         rep(3,nrow(Testes_Down_Age_GOBP  )))

## MF
ovary.GOMF          <- rbind(Ovary_Down_MAge_GOMF,
                             Ovary_Up_MAge_GOMF  ,
                             Ovary_Down_Age_GOMF ,
                             Ovary_Up_Age_GOMF   )
ovary.GOMF$cluster  <- c(rep(1,nrow(Ovary_Down_MAge_GOMF)),
                         rep(2,nrow(Ovary_Up_MAge_GOMF  )),
                         rep(3,nrow(Ovary_Down_Age_GOMF )),
                         rep(4,nrow(Ovary_Up_Age_GOMF   )))

testis.GOMF         <- rbind(Testes_Down_MAge_GOMF ,
                             Testes_Up_MAge_GOMF   ,
                             Testes_Down_Age_GOMF  )
testis.GOMF$cluster <- c(rep(1,nrow(Testes_Down_MAge_GOMF )),
                         rep(2,nrow(Testes_Up_MAge_GOMF   )),
                         rep(3,nrow(Testes_Down_Age_GOMF  )))

## CC
ovary.GOCC          <- rbind(Ovary_Down_MAge_GOCC,
                             Ovary_Up_MAge_GOCC  ,
                             Ovary_Down_Age_GOCC ) # Up age did not have any significant CCs, omit
ovary.GOCC$cluster  <- c(rep(1,nrow(Ovary_Down_MAge_GOCC)),
                         rep(2,nrow(Ovary_Up_MAge_GOCC  )),
                         rep(3,nrow(Ovary_Down_Age_GOCC )))

testis.GOCC         <- rbind(Testes_Down_MAge_GOCC ,
                             Testes_Up_MAge_GOCC   ,
                             Testes_Down_Age_GOCC  )
testis.GOCC$cluster <- c(rep(1,nrow(Testes_Down_MAge_GOCC )),
                         rep(2,nrow(Testes_Up_MAge_GOCC   )),
                         rep(3,nrow(Testes_Down_Age_GOCC  ))) # Up age did not have any significant CCs, omit


# write and export all significant GO results tables for supplementary information
write.table(ovary.GOBP , file = paste0("./Results/4_GO_enrichment/", Sys.Date(), "_GOstats_Ovary_Summary_Table_GOBP_FDR5.txt"   ), sep = "\t" , row.names = F, col.names = T, quote = F)
write.table(testis.GOBP, file = paste0("./Results/4_GO_enrichment/", Sys.Date(), "_GOstats_Testes_Summary_Table_GOBP_FDR5.txt"  ), sep = "\t" , row.names = F, col.names = T, quote = F)

write.table(ovary.GOMF , file = paste0("./Results/4_GO_enrichment/", Sys.Date(), "_GOstats_Ovary_Summary_Table_GOMF_FDR5.txt"   ), sep = "\t" , row.names = F, col.names = T, quote = F)
write.table(testis.GOMF, file = paste0("./Results/4_GO_enrichment/", Sys.Date(), "_GOstats_Testes_Summary_Table_GOMF_FDR5.txt"  ), sep = "\t" , row.names = F, col.names = T, quote = F)

write.table(ovary.GOCC , file = paste0("./Results/4_GO_enrichment/", Sys.Date(), "_GOstats_Ovary_Summary_Table_GOCC_FDR5.txt"   ), sep = "\t" , row.names = F, col.names = T, quote = F)
write.table(testis.GOCC, file = paste0("./Results/4_GO_enrichment/", Sys.Date(), "_GOstats_Testes_Summary_Table_GOCC_FDR5.txt"  ), sep = "\t" , row.names = F, col.names = T, quote = F)


#################################################
# 5. Create Bubble Plots of GO Analysis

##################################################################
######################         GO BP        ######################
# run bubble processing females
Ovary_Down_MAge_Bubble_Plot <- bubble_plot(Ovary_Down_MAge_GOBP, "F", "Ovary_Down_MAge")
Ovary_Up_MAge_Bubble_Plot   <- bubble_plot(Ovary_Up_MAge_GOBP  , "F", "Ovary_Up_MAge")
Ovary_Down_Age_Bubble_Plot  <- bubble_plot(Ovary_Down_Age_GOBP , "F", "Ovary_Down_Age")
Ovary_Up_Age_Bubble_Plot    <- bubble_plot(Ovary_Up_Age_GOBP   , "F", "Ovary_Up_Age")

#group female bubbleplots
female_plots <- as.ggplot(Ovary_Down_MAge_Bubble_Plot) +
                as.ggplot(Ovary_Up_MAge_Bubble_Plot) +
                as.ggplot(Ovary_Down_Age_Bubble_Plot) +
                as.ggplot(Ovary_Up_Age_Bubble_Plot) +
                guide_area()+
                plot_layout(ncol = 2, guides = "collect") +
                plot_annotation(tag_levels = "A", title = "Ovary GO BP Bubble Plots",
                   theme = theme(plot.title = element_text(size = 26)))

my.bubbleplot.out <- paste0("./Results/4_GO_enrichment/", Sys.Date(),"_AGING_Bubbleplots_females_GOBP_FDR5.pdf")
pdf(my.bubbleplot.out, onefile = F, width =  16, height = 15)
female_plots
dev.off()


# run bubble processing males  (up age - too few genes)
Testes_Down_MAge_Bubble_Plot <- bubble_plot(Testes_Down_MAge_GOBP, "M", "Testes_Down_MAge")
Testes_Up_MAge_Bubble_Plot   <- bubble_plot(Testes_Up_MAge_GOBP  , "M", "Testes_Up_MAge")
Testes_Down_Age_Bubble_Plot  <- bubble_plot(Testes_Down_Age_GOBP , "M", "Testes_Down_Age")

#group male bubbleplots
male_plots <- as.ggplot(Testes_Down_MAge_Bubble_Plot) +
              as.ggplot(Testes_Up_MAge_Bubble_Plot) +
              as.ggplot(Testes_Down_Age_Bubble_Plot) +
              guide_area()+
              plot_layout(ncol = 2, guides = "collect") +
              plot_annotation(tag_levels = "A", title = "Testis GOBP Bubble Plots",
                  theme = theme(plot.title = element_text(size = 26)))

my.bubbleplot.out <- paste0("./Results/4_GO_enrichment/", Sys.Date(),"_AGING_Bubbleplots_males_GOBP_FDR5.pdf")
pdf(my.bubbleplot.out, onefile = F, width =  15, height = 15)
male_plots
dev.off()


##################################################################
######################         GO MF        ######################
# run bubble processing females
Ovary_Down_MAge_Bubble_Plot <- bubble_plot(Ovary_Down_MAge_GOMF, "F", "Ovary_Down_MAge")
Ovary_Up_MAge_Bubble_Plot   <- bubble_plot(Ovary_Up_MAge_GOMF  , "F", "Ovary_Up_MAge")
Ovary_Down_Age_Bubble_Plot  <- bubble_plot(Ovary_Down_Age_GOMF , "F", "Ovary_Down_Age")
Ovary_Up_Age_Bubble_Plot    <- bubble_plot(Ovary_Up_Age_GOMF   , "F", "Ovary_Up_Age")

#group female bubbleplots
female_plots <- as.ggplot(Ovary_Down_MAge_Bubble_Plot) +
  as.ggplot(Ovary_Up_MAge_Bubble_Plot) +
  as.ggplot(Ovary_Down_Age_Bubble_Plot) +
  as.ggplot(Ovary_Up_Age_Bubble_Plot) +
  guide_area()+
  plot_layout(ncol = 2, guides = "collect") +
  plot_annotation(tag_levels = "A", title = "Ovary GO MF Bubble Plots",
                  theme = theme(plot.title = element_text(size = 26)))

my.bubbleplot.out <- paste0("./Results/4_GO_enrichment/", Sys.Date(),"_AGING_Bubbleplots_females_GOMF_FDR5.pdf")
pdf(my.bubbleplot.out, onefile = F, width =  16, height = 15)
female_plots
dev.off()


# run bubble processing males  (up age - too few genes)
Testes_Down_MAge_Bubble_Plot <- bubble_plot(Testes_Down_MAge_GOMF, "M", "Testes_Down_MAge")
Testes_Up_MAge_Bubble_Plot   <- bubble_plot(Testes_Up_MAge_GOMF  , "M", "Testes_Up_MAge")
Testes_Down_Age_Bubble_Plot  <- bubble_plot(Testes_Down_Age_GOMF , "M", "Testes_Down_Age")

#group male bubbleplots
male_plots <- as.ggplot(Testes_Down_MAge_Bubble_Plot) +
  as.ggplot(Testes_Up_MAge_Bubble_Plot) +
  as.ggplot(Testes_Down_Age_Bubble_Plot) +
  guide_area()+
  plot_layout(ncol = 2, guides = "collect") +
  plot_annotation(tag_levels = "A", title = "Testis GOMF Bubble Plots",
                  theme = theme(plot.title = element_text(size = 26)))

my.bubbleplot.out <- paste0("./Results/4_GO_enrichment/", Sys.Date(),"_AGING_Bubbleplots_males_GOMF_FDR5.pdf")
pdf(my.bubbleplot.out, onefile = F, width =  15, height = 15)
male_plots
dev.off()


##################################################################
######################         GO CC        ######################
# run bubble processing females
Ovary_Down_MAge_Bubble_Plot <- bubble_plot(Ovary_Down_MAge_GOCC, "F", "Ovary_Down_MAge")
Ovary_Up_MAge_Bubble_Plot   <- bubble_plot(Ovary_Up_MAge_GOCC  , "F", "Ovary_Up_MAge")
Ovary_Down_Age_Bubble_Plot  <- bubble_plot(Ovary_Down_Age_GOCC , "F", "Ovary_Down_Age")
# no ovary uo with ag pathways

#group female bubbleplots
female_plots <- as.ggplot(Ovary_Down_MAge_Bubble_Plot) +
  as.ggplot(Ovary_Up_MAge_Bubble_Plot) +
  as.ggplot(Ovary_Down_Age_Bubble_Plot) +
  guide_area()+
  plot_layout(ncol = 2, guides = "collect") +
  plot_annotation(tag_levels = "A", title = "Ovary GO CC Bubble Plots",
                  theme = theme(plot.title = element_text(size = 26)))

my.bubbleplot.out <- paste0("./Results/4_GO_enrichment/", Sys.Date(),"_AGING_Bubbleplots_females_GOCC_FDR5.pdf")
pdf(my.bubbleplot.out, onefile = F, width =  16, height = 15)
female_plots
dev.off()


# run bubble processing males  (up age - too few genes)
Testes_Down_MAge_Bubble_Plot <- bubble_plot(Testes_Down_MAge_GOCC, "M", "Testes_Down_MAge")
Testes_Up_MAge_Bubble_Plot   <- bubble_plot(Testes_Up_MAge_GOCC  , "M", "Testes_Up_MAge")
Testes_Down_Age_Bubble_Plot  <- bubble_plot(Testes_Down_Age_GOCC , "M", "Testes_Down_Age")

#group male bubbleplots
male_plots <- as.ggplot(Testes_Down_MAge_Bubble_Plot) +
  as.ggplot(Testes_Up_MAge_Bubble_Plot) +
  as.ggplot(Testes_Down_Age_Bubble_Plot) +
  guide_area()+
  plot_layout(ncol = 2, guides = "collect") +
  plot_annotation(tag_levels = "A", title = "Testis GO CC Bubble Plots",
                  theme = theme(plot.title = element_text(size = 26)))

my.bubbleplot.out <- paste0("./Results/4_GO_enrichment/", Sys.Date(),"_AGING_Bubbleplots_males_GOCC_FDR5.pdf")
pdf(my.bubbleplot.out, onefile = F, width =  15, height = 15)
male_plots
dev.off()


####################################################################################
# 5. Generate Heatmaps from "piRNA metabolic process" Genes (GO term "GO:0034587")

# read in female and male normalized gene count matrices from 1_Ovary_Gene_DGE.R and 2_Testes_Gene_DGE.R
ovary_norm_counts  <- read.table("./Results/1_Ovary_DGE/2022-07-13_Ovary_LRT_Female_Normalized_DEseq2_GeneTE_Count_Matrix.txt")
testis_norm_counts <- read.table("./Results/2_Testis_DGE/2022-07-13_Testes_LRT_Male_Normalized_DEseq2_GeneTE_Count_Matrix.txt")

# grab gene IDs from the gsc object for "piRNA metabolic process" and merge to get human IDs
piRNA_met_pro.genes       <- as.vector(gsc[["GO:0034587"]]@geneIds)
piRNA_met_pro             <- conversion_table[conversion_table$gene_id %in% piRNA_met_pro.genes, ]
piRNA_met_pro$concat_name <- paste(piRNA_met_pro$gene_id, piRNA_met_pro$Human_Gene_Symbol,sep=" | ")

# extract normalized counts for "piRNA metabolic process" genes in each tissue
fem_pi  <- as.data.frame(ovary_norm_counts[piRNA_met_pro$gene_id,])
male_pi <- as.data.frame(testis_norm_counts[piRNA_met_pro$gene_id,])

# normalize median values from young samples in each sex for heatmap plotting
fem_pi.norm    <- fem_pi/apply(fem_pi[,c(1:5)] , 1, median)
male_pi.norm   <- male_pi/apply(male_pi[,c(1:4)], 1, median)

# merge and set rownames for easier merges
piRNApath.norm <- cbind(fem_pi.norm, male_pi.norm)
rownames(piRNApath.norm) <- piRNA_met_pro$concat_name

# get median logFC between youth and mid-age (already logged, so it's a substraction in log space)
f.sort <- sort(apply(fem_pi[,6:10],1,median)-apply(fem_pi[,1:5],1,median), index.return = T)

#output heatmaps
pdf(paste("./Results/4_GO_enrichment/",  Sys.Date(), "_Heatmap_of_gene_expression_piRNA_metab_process_Young_median_Norm.pdf"), onefile = F, width = 20, height = 10)
pheatmap(piRNApath.norm[f.sort$ix,],
         cluster_cols = F,
         cluster_rows = F,
         colorRampPalette(rev(c("#CC3333","#FF9999","#FFCCCC","white","#CCCCFF","#9999FF","#333399")))(50),
         show_rownames = T, scale="none",
         border = NA, cellheight = 15,
         main = paste("piRNA metabolic process Females/Males"), cellwidth = 15)
dev.off()


####################################
# extract normalized counts for "piRNA metabolic process" genes in each tissue for supplementary table
piRNApath.exp <- cbind(fem_pi, male_pi)
rownames(piRNApath.exp) <- piRNA_met_pro$concat_name

write.table(piRNApath.exp, file = paste0("./Results/4_GO_enrichment/", Sys.Date(), "_piRNA_metabolic_process_expression_table.txt"  ), sep = "\t" , row.names = T, col.names = T, quote = F)


##########################################################
sink(file = paste("./Results/4_GO_enrichment/", Sys.Date(),"_GO_analysis_session_Info.txt", sep =""))
sessionInfo()
sink()
```
