## Supplemental Code for "Dynamic regulation of gonadal transposon control across the lifespan of the naturally short-lived African turquoise killifish": 5_PCA_Plot_Generation_v2.R

```
# set working directory to directory containing mRNA count matrix
# setwd("/Users/bryanteefy/Dropbox/PIWI/CODE/3_Differential_Gene_Expression_Analysis")
setwd("/Users/berenice/Dropbox/Manuscripts_and_Publications/2022/2022_Bryan_PIWI_manuscript/CODE/3_Differential_Gene_Expression_Analysis")

# PCA Plot Generation for Genes, TEs, and piRNA mapping to TEs

# R version 4.1.2 (2021-11-01)

#load libraries
library(DESeq2) # DESeq2_1.34.0

##############################################################################
# 1. Get Gene/TE Count Matrix and process with DESeq2 VST normalization

########## A. Load and preprocess TEtranscript counts
# load raw count matrix
my.gonad <- read.table("./Input/Gene_TE_Counts.cntTable", header = T)

#rename
colnames(my.gonad) <- c("GeneName","MF1","MF2","MF3","MF4","MF5","MM1","MM2","MM3","MM5","OF1","OF3","OF4","OF5","OM1","OM3","OM4","OM5","YF1","YF2","YF3","YF4","YF5","YM1","YM2","YM3","YM5")

#reorganize by sex and age
my.gonad <- my.gonad[,c("GeneName",
                        "YM1",
                        "YM2",
                        "YM3",
                        "YM5",
                        "MM1",
                        "MM2",
                        "MM3",
                        "MM5",
                        "OM1",
                        "OM3",
                        "OM4",
                        "OM5",
                        "YF1",
                        "YF2",
                        "YF3",
                        "YF4",
                        "YF5",
                        "MF1",
                        "MF2",
                        "MF3",
                        "MF4",
                        "MF5",
                        "OF1",
                        "OF3",
                        "OF4",
                        "OF5"
)]

my.Sex  <- c(rep("M",12),rep("F",14))
my.Age  <- c(rep(5,4),rep(10,4),rep(15,4),rep(5,5),rep(10,5),rep(15,4))

# format for processing
rownames(my.gonad) <- my.gonad$GeneName

# get the genes with no reads out
my.good <- which(apply(my.gonad[,-1]>0, 1, sum) >= 6) # see deseq2 vignette, need to remove too low genes
my.filtered.matrix <- my.gonad[my.good,-1] # 22458  transcripts

# Prerequisite: round the counts to nearest integer
# (TEtranscripts will create fractional counts from multimapping reads; integers required for DEseq2)
rounded_matrix <- round(my.filtered.matrix)

########## B. Get VST Normalized Expression Matrix
# design matrix
dataDesign = data.frame( row.names = colnames(rounded_matrix), 
                         age = my.Age,
                         sex = my.Sex)

# get matrix using age as a modeling covariate
dds <- DESeqDataSetFromMatrix(countData = rounded_matrix,
                              colData = dataDesign,
                              design = ~ age + sex)

# run DESeq normalizations and export results
dds.deseq <- DESeq(dds)

# normalize expression values
tissue.cts <- getVarianceStabilizedData(dds.deseq)

# define colors 
my.colors                                     <- rep("deepskyblue1",26)
my.colors[grep("MM",colnames(tissue.cts))]    <- "deepskyblue3"
my.colors[grep("OM",colnames(tissue.cts))]    <- "deepskyblue4"
my.colors[grep("YF",colnames(tissue.cts))]    <- "deeppink1"
my.colors[grep("MF",colnames(tissue.cts))]    <- "deeppink3"
my.colors[grep("OF",colnames(tissue.cts))]    <- "deeppink4"


# define shapes
my.pch                                     <- rep(19,26)
my.pch[grep("MM",colnames(tissue.cts))]    <- 17
my.pch[grep("OM",colnames(tissue.cts))]    <- 15
my.pch[grep("YF",colnames(tissue.cts))]    <- 19
my.pch[grep("MF",colnames(tissue.cts))]    <- 17
my.pch[grep("OF",colnames(tissue.cts))]    <- 15

##############################################################################
# 2. Define PCA and plotting functions [valid for Genes, TEs and piRNA cluster]

# PCA analysis
# define PCA function
make_PCA <- function(count.mat) {
  my.pos.var <- apply(count.mat,1,var) > 0
  my.pca <- prcomp(t(count.mat[my.pos.var,]),scale = TRUE)
  x <- my.pca$x[,1]
  y <- my.pca$x[,2]
  my.summary <- summary(my.pca)
  return(list(x, y, my.summary))
}

# define plotting function
plot_function <- function(x,y,z, pt.shp) {
  plot(x[[1]], x[[2]],
       cex=3, pch = pt.shp, col = y,
       xlab = paste('PC1 (', round(100*x[[3]]$importance[,1][2],1),"%)", sep=""),
       ylab = paste('PC2 (', round(100*x[[3]]$importance[,2][2],1),"%)", sep=""),
       xlim = z,
       ylim = z,
       cex.lab = 1.5,
       cex.axis = 1.5) 
}

##############################################################################
# 3. Make PCA Plots for Genes

# set prefix
my.outprefix <- paste(Sys.Date(),"Genes_Gonad",sep="_")

## retain only genes for genic PCA
#both sexes
tissue.cts.genes <- tissue.cts[!grepl("NotFur", rownames(tissue.cts)), ]

#females
tissue.cts.genes.F <- tissue.cts.genes[,grepl("F", colnames(tissue.cts.genes)) ]

#males
tissue.cts.genes.M <- tissue.cts.genes[,!grepl("F", colnames(tissue.cts.genes)) ]

# both sexes
# plot and output
my.pca.out <- paste0("./Results/5_PCA_plotting/", my.outprefix,"_Gene_PCA_plot.pdf")
pdf(my.pca.out)
plot_function(make_PCA(tissue.cts.genes), my.colors, c(-150,150), my.pch)
dev.off()

#### females
#plot and output
my.pca.out <- paste0("./Results/5_PCA_plotting/", my.outprefix,"_Ovary_Gene_PCA_plot.pdf")
pdf(my.pca.out)
plot_function(make_PCA(tissue.cts.genes.F), my.colors[13:26], c(-150,150), my.pch[13:26])
dev.off()

#### males
#plot and output
my.pca.out <- paste0("./Results/5_PCA_plotting/", my.outprefix,"_Testes_Gene_PCA_plot.pdf")
pdf(my.pca.out)
plot_function(make_PCA(tissue.cts.genes.M), my.colors[1:12], c(-250,250), my.pch[1:12])
dev.off()

##############################################################################
# 4. Make PCA Plots for TEs only

# set prefix
my.outprefix <- paste(Sys.Date(),"TE_only_Gonad",sep="_")

## retain only TEs
#both sexes
tissue.cts.TEs <- tissue.cts[grepl("NotFur", rownames(tissue.cts)), ]

#females
tissue.cts.TEs.F <- tissue.cts.TEs[,grepl("F", colnames(tissue.cts.TEs)) ]

#males
tissue.cts.TEs.M <- tissue.cts.TEs[,!grepl("F", colnames(tissue.cts.TEs)) ]

# both sexes
#plot and output
my.pca.out <- paste0("./Results/5_PCA_plotting/", my.outprefix,"_TE_PCA_plot.pdf")
pdf(my.pca.out)
plot_function(make_PCA(tissue.cts.TEs), my.colors, c(-50,50), my.pch)
dev.off()

# females
#plot and output
my.pca.out <- paste0("./Results/5_PCA_plotting/", my.outprefix,"_Ovary_TE_PCA_plot.pdf")
pdf(my.pca.out)
plot_function(make_PCA(tissue.cts.TEs.F), my.colors[13:26], c(-75,75), my.pch[13:26])
dev.off()

# males
#plot and output
my.pca.out <- paste0("./Results/5_PCA_plotting/", my.outprefix,"_Testes_TE_PCA_plot.pdf")
pdf(my.pca.out)
plot_function(make_PCA(tissue.cts.TEs.M), my.colors[1:12], c(-100,100), my.pch[1:12])
dev.off()

##############################################################################
#5.  Repeat with piRNAs mapping to TEs #

#load count matrix (bowtie mapping, counting over TEs, piRNA clusters, rRNA and genes with fractional counts by FeatureCounts)
my.gonad <- read.table("./Input/piRNA_count_matrix.txt", header = T)

#preprocess piRNA count matrix to remove rRNA counts and remove non-count data
my.gonad <- my.gonad[,c(1,7:32)]
my.gonad <- my.gonad[!grepl("rRNA", my.gonad$Geneid), ] 

#remove excess from colnames
colnames(my.gonad) <- gsub("[.][.][.]", "", colnames(my.gonad))
colnames(my.gonad) <- gsub("[.]bam", "", colnames(my.gonad))

my.Sex  <- c(rep("M",12),rep("F",14))
my.Age  <- c(rep(5,4),rep(10,4),rep(15,4),rep(5,5),rep(10,5),rep(15,4))

# format for processing
rownames(my.gonad) <- my.gonad$Geneid

# get the genes with no reads out
my.good <- which(apply(my.gonad[,-1]>0, 1, sum) >= 6) # see deseq2 vignette, need to remove too low genes
my.filtered.matrix <- my.gonad[my.good,-1] # 19959  transcripts/features

#Prerequisite: round the counts to nearest integer (DEseq2 needs integers)
rounded_matrix <- round(my.filtered.matrix)

########## B. Get VST Normalized Expression Matrix
# design matrix
dataDesign = data.frame( row.names = colnames(rounded_matrix), 
                         age       = my.Age,
                         sex       = my.Sex)

# get matrix using age as a modeling covariate
dds <- DESeqDataSetFromMatrix(countData = rounded_matrix,
                              colData   = dataDesign,
                              design    = ~ age + sex)

# run DESeq normalizations and export results
dds.deseq <- DESeq(dds)

# normalize expression values
tissue.cts <- getVarianceStabilizedData(dds.deseq)

#keep only TEs features expression for analysis
tissue.cts <- tissue.cts[grepl("NotFur", rownames(tissue.cts)),]

################################################################
# 6. Plot and Export piRNA Mapping to TE PCA plots 

#females
tissue.cts.F <- tissue.cts[,grepl("F", colnames(tissue.cts)) ]

#males
tissue.cts.M <- tissue.cts[,!grepl("F", colnames(tissue.cts)) ]

# both sexes
#plot and output
my.pca.out <- paste0("./Results/5_PCA_plotting/",  my.outprefix,"_piRNA_PCA_plot.pdf")
pdf(my.pca.out)
plot_function(make_PCA(tissue.cts), my.colors, c(-75,75), my.pch)
dev.off()

# females
#plot and output
my.pca.out <- paste0("./Results/5_PCA_plotting/",  my.outprefix,"_Ovary_piRNA_PCA_plot.pdf")
pdf(my.pca.out)
plot_function(make_PCA(tissue.cts.F), my.colors[13:26], c(-50,50), my.pch[13:26])
dev.off()

# males
#plot and output
my.pca.out <- paste0("./Results/5_PCA_plotting/",  my.outprefix,"_Testes_piRNA_PCA_plot.pdf")
pdf(my.pca.out)
plot_function(make_PCA(tissue.cts.M), my.colors[1:12], c(-100,100), my.pch[1:12])
dev.off()

#######################
sink(file = paste("./Results/5_PCA_plotting/", Sys.Date(),"_PCA_code_session_Info.txt", sep =""))
sessionInfo()
sink()
```
