## Supplemental Code for "Dynamic regulation of gonadal transposon control across the lifespan of the naturally short-lived African turquoise killifish": README_Differential_Gene_Expression_Analysis.txt

########################################################
   README -- 3_Differential_Gene_Expression_Analysis
########################################################

# All scripts ran with R version 4.1.2 (2021-11-01)

*** All R package versions indicated in cognate scripts ***
	
	1. Ovarian differential gene expression using likelihood ratio testing (LRT) with DESeq2 
		- 1_Ovary_Gene_DGE.R
		
	2. Testicular differential gene expression using LRT with DESeq2
		- 2_Testes_Gene_DGE.R
		
	3. How to generate Human/Killifish conversion tables using local BLAST, parse protein/gene conversions and human gene annotations
	 	- 3a_BLAST_preprocessing.txt
	 	- 3b_GTF_preprocessing.R
	 	
	4. Runs Hypergeometric GO analysis, also outputs "piRNA metabolic process" expression tables
		- 4_GO_Enrichment_Analysis.R: main script to run GO overrepresentation analysis
		- 4_Functions_for_GO_Enrich.R: functions required to run 4_GO_Enrichment_Analysis.R

	5. PCA Plot Generation for Genes, TEs, and piRNA clusters (DEseq2 normalization and PCA plotting)
		- 5_PCA_Plot_Generation.R
	
