## Supplemental Code for "Dynamic regulation of gonadal transposon control across the lifespan of the naturally short-lived African turquoise killifish": 2_Testes_TE_DGE.R

```
# set working directory to directory containing mRNA count matrix
 setwd("/Users/bryanteefy/Dropbox/PIWI/CODE/4_TE_analysis")
#setwd("/Users/berenice/Dropbox/Manuscripts_and_Publications/2022/2022_Bryan_PIWI_manuscript/CODE/4_TE_analysis")

# Male Differential TE Expression Analysis (LRT)

# R version 4.1.2 (2021-11-01)

# load libraries              
library(DESeq2)               # DESeq2_1.34.0
library('bitops')             # bitops_1.0-7
library(RColorBrewer)         # RColorBrewer_1.1-3
library(pheatmap)             # pheatmap_1.0.12
library(DEGreport)            # DEGreport_1.30.3 
library(tibble)               # tibble_3.1.7
library(dplyr)                # dplyr_1.0.9 
library(ggpubr)               # ggpubr_0.4.0 
library(ggplotify)            # ggplotify_0.1.0 
library(SummarizedExperiment) # SummarizedExperiment_1.24.0

##############################################################################################################################
# load count matrix
my.gonad <- read.table("./Input/Gene_TE_Counts.cntTable", header = T)

# reorganize by sex and age
my.gonad <- my.gonad[,c("GeneName",
                        "YM1",
                        "YM2",
                        "YM3",
                        "YM5",
                        "MM1",
                        "MM2",
                        "MM3",
                        "MM5",
                        "OM1",
                        "OM3",
                        "OM4",
                        "OM5",
                        "YF1",
                        "YF2",
                        "YF3",
                        "YF4",
                        "YF5",
                        "MF1",
                        "MF2",
                        "MF3",
                        "MF4",
                        "MF5",
                        "OF1",
                        "OF3",
                        "OF4",
                        "OF5"
)]

#generate age variables for females
my.Age  <- c(rep("Young",4),rep("Middle",4),rep("Old",4))
rownames(my.gonad) <- my.gonad$GeneName

#retain only female data 
my.gonad <- my.gonad[,c(1:13)]

# get the genes with no reads out
my.good <- which(apply(my.gonad[,-1]>0, 1, sum) >= 6)
my.filtered.matrix <- my.gonad[my.good,-1]

# Prerequisite: round the counts to nearest integer 
# (TEtranscripts will create fractional counts from multimapping reads; integers required for DEseq2)
rounded_matrix <- round(my.filtered.matrix)

#set prefix
my.outprefix <- paste(Sys.Date(),"Testes_TE_analysis_LRT",sep="_")

#set colors
colors <- c("deepskyblue", "deepskyblue3", "deepskyblue4")

################################################################################################################
# 2. DESeq2 LRT DGE on cleaned data

# design matrix
dataDesign = data.frame( row.names = colnames(rounded_matrix), 
                         age = my.Age)

# get matrix using age as a modeling covariate
dds <- DESeqDataSetFromMatrix(countData = rounded_matrix,
                              colData = dataDesign,
                              design = ~ age)

# run DESeq normalizations and export results with LRT
dds_lrt <- DESeq(dds, test="LRT", reduced = ~ 1)

# Extract results
res_LRT <- results(dds_lrt)

# Create a tibble for LRT results
res_LRT_tb <- res_LRT %>%
  data.frame() %>%
  rownames_to_column(var = "gene") %>% 
  as_tibble()

# write output of LRT DGE Analysis
write.table(res_LRT_tb, file = paste0("./Results/2_Testis_TE_DGE/", my.outprefix,"_Testicular_LRT_DGE_Table_ALL.txt"), sep = "\t" , row.names = F, quote=F)

# Subset to return genes with padj < 1e-6
# LRT is very lenient, so a stringent cutoff is required
# see: https://hbctraining.github.io/DGE_workshop_salmon/lessons/08_DGE_LRT.html
sigLRT_genes <- res_LRT_tb %>% filter(padj < 1e-6)

# Get number of significant genes + TEs
nrow(sigLRT_genes) # 628  genes + TEs

# subset/sort results for faster cluster finding
clustering_sig_genes <- sigLRT_genes %>% arrange(padj)

# get VST-normalized expression values
tissue.cts <- getVarianceStabilizedData(dds_lrt)

# write out normalized count matrix for use in GO analyses
write.table(tissue.cts, file = paste0("./Results/2_Testis_TE_DGE/", my.outprefix,"_Female_Normalized_DEseq2_GeneTE_Count_Matrix.txt"), sep = "\t" , row.names = T, quote = F)

# Extract significant genes
sig_genes <- tissue.cts[clustering_sig_genes$gene, ]

### Try subsetting after clustering? Using the TEs only to cluster leads to errors
# retain TEs only; no genes
#### FishTEDB TEs all have "NotFur" prefix in name, which helps filter them
sig_genes_TE <- sig_genes[grepl("NotFur", rownames(sig_genes)), ] # 25 TEs

########## Because all male significant TEs are downregulated with age, degpattern throws an error
# for the purpose of plotting the jitter box plot;
#####       - we cluster on all genes + TE
#####       - we extract the TE rows (all belong to cluster 1)
#####       - we make a "mock" object for plotting containing just the TEs so that degPlotCluster works

# Use the `degPatterns` function from the 'DEGreport' package to show gene clusters across sample groups
#### yields warning "geom_path: Each group consists of only one observation. Do you need to adjust the group aesthetic?"
clusters <- degPatterns(sig_genes, metadata = dataDesign, time = "age", col=NULL, minc = 1)
clusters.norm.TEs <- clusters$normalized[grepl("NotFur", clusters$normalize$genes), ] # 25 TEs
summary(clusters.norm.TEs) ### all in cluster 1

# update age variable for plotting
clusters.norm.TEs$age <- clusters.norm.TEs$merge
clusters.norm.TEs$age <- factor(clusters.norm.TEs$age, levels=c("one_groupYoung", "one_groupMiddle", "one_groupOld"))

my.clusters.out <- paste0("./Results/2_Testis_TE_DGE/", my.outprefix,"_AGING_TEs_boxplot_grouped_clusters_FDR1e-6.pdf")
pdf(my.clusters.out, onefile = F)
degPlotCluster(clusters.norm.TEs, "age", lines=TRUE, min_genes = 1, col = "age") + theme_bw() + scale_color_manual(values=colors[c(1:3)])
dev.off()

# create merged table of LRT/clustering results
# non significant genes will have 'NA' after outer join operation
res.TEs <- data.frame(res_LRT_tb)[grepl("NotFur", res_LRT_tb$gene), ]

# All significant all belong to cluster 1
res.TEs$cluster <- NA
res.TEs$cluster[res.TEs$gene %in% rownames(sig_genes_TE)] <- 1

write.table(res.TEs, file = paste0("./Results/2_Testis_TE_DGE/", my.outprefix,"_DESeq2_LRT_TE_DGE_Table_ALL_with_DGEPatterns_Clusters_FDR1e-6.txt"), sep = "\t" , row.names = F, quote = F)


######################################################################################
# 3. Generate Heatmaps color coded by TE Family

# format for heatmap function
group1 <- data.frame("genes" = rownames(sig_genes_TE), cluster = rep(1,nrow(sig_genes_TE)))

# Create function that 1. corrects modified TE names 2. Specifies TE family 3. generates heatmap
# x <- group1
heatmap_TE <- function (x) {
  gene_names <- x$genes
  gene_names <- gsub("\\.", ":", gene_names)
  test2 <- as.data.frame(gene_names)
  matrix <- as.data.frame(tissue.cts)
  matrix$gene_names <- rownames(matrix)
  # matrix$gene_names <- gsub("-", ":", matrix$gene_names) [names were not jumbled]
  matrix2 <- merge(matrix, test2, by = "gene_names")
  matrix3 <- matrix2[,c(2:ncol(matrix2))]
  num <- nrow(matrix3)
  new_names <- gsub(".+:", "", gene_names)
  # merge unclear and Unknown families
  new_names <- gsub("unclear", "Unknown", new_names)
  NNs <- as.character(factor(new_names))
  rownames(matrix3) <- paste0("row_", seq(nrow(matrix3)))
  annotation_row = data.frame(
    Family = factor(NNs)
  )
  rownames(annotation_row) = rownames(matrix3)
  NNz <- order(NNs)
  order_of_TEs <- paste0("row_", NNz)
  # create colors for each group
  annoCol<-list(Family=c(DNA="firebrick1", LINE="darkturquoise", LTR="cornflowerblue", SINE="blue", Unknown="gold"))
  hm <- pheatmap(matrix3[order_of_TEs,],
                 cluster_cols = F,
                 cluster_rows = T,
                 colorRampPalette(rev(c("#CC3333","#FF9999","#FFCCCC","white","#CCCCFF","#9999FF","#333399")))(50),
                 show_rownames = F, scale="row",
                 border = NA,cellheight = 1,
                 main = paste("TEs DE with Age", length(order_of_TEs), "TEs"), cellwidth = 15,
                 annotation_row = annotation_row,
                 annotation_colors = annoCol)
  return(hm)
}

my.heatmap.out <- paste0("./Results/2_Testis_TE_DGE/", my.outprefix,"_AGING_TE_Heatmap_FDR1e-6.pdf")
pdf(my.heatmap.out, onefile = F, height = 10, width = 10)
heatmap_TE(group1)
dev.off()


######################################################################################
# 4. Expression Correlation Heatmap Generation

#restrict analysis to only TEs, not genes
TE_exp <- tissue.cts[grepl("NotFur", rownames(tissue.cts)), ]

#obtain a correlation matrix of the gene expression data
correlation <- as.matrix(cor(TE_exp, method = "spearman"))

#arrange data
age <- data.frame("AgeGroup" = c(rep("YM",4),rep("MM",4),rep("OM",4) ))
rownames(age) = colnames(correlation)
age_cols = list(AgeGroup = c(YM = "deepskyblue", MM = "deepskyblue3", OM = "deepskyblue4"))

#plot and export
my.heatmap.out <- paste0("./Results/2_Testis_TE_DGE/", my.outprefix,"_AGING_TEs_Exp_Correlation_Heatmap.pdf")
pdf(my.heatmap.out, onefile = F, height = 10, width = 10)
pheatmap(rbind(correlation,rep(0.9,nrow(correlation))),
         cluster_cols = F,
         cluster_rows = F,
         colorRampPalette(rev(c("#CC3333","#FF9999","#FFCCCC","white")))(20),
         show_rownames = T, scale="none",
         main = "Testes TE Expression Correlation Matrix", 
         cellwidth = 40, cellheight = 40, display_numbers = T, number_color = "white", fontsize_number = 12,
         annotation_col = age,
         annotation_colors = age_cols)
dev.off()
dev.off()

#######################
sink(file = paste("./Results/2_Testis_TE_DGE/", my.outprefix,"_session_Info.txt", sep =""))
sessionInfo()
sink()
```
