## Supplemental Code for "Dynamic regulation of gonadal transposon control across the lifespan of the naturally short-lived African turquoise killifish": 3_Fractional_TE_Analysis.R

```
# set working directory to directory containing mRNA count matrix
 setwd("/Users/bryanteefy/Dropbox/PIWI/CODE/4_TE_analysis")
#setwd("/Users/berenice/Dropbox/Manuscripts_and_Publications/2022/2022_Bryan_PIWI_manuscript/CODE/4_TE_analysis")

# Fractional TE Representation in Gonadal Transcriptomes

# R version 4.1.2 (2021-11-01)

#load libraries
library(reshape2)   # reshape2_1.4.4
library(beeswarm)   # beeswarm_0.4.0

##############################################################################################################################
# load count matrix
my.gonad <- read.table("./Input/Gene_TE_Counts.cntTable", header = T)

# set prefix
my.outprefix <- paste(Sys.Date(),"TE_percentages",sep="_")

colnames(my.gonad) <- c("GeneName","MF1","MF2","MF3","MF4","MF5","MM1","MM2","MM3","MM5","OF1","OF3","OF4","OF5","OM1","OM3","OM4","OM5","YF1","YF2","YF3","YF4","YF5","YM1","YM2","YM3","YM5")

my.gonad <- my.gonad[,c("GeneName",
                        "YM1",
                        "YM2",
                        "YM3",
                        "YM5",
                        "MM1",
                        "MM2",
                        "MM3",
                        "MM5",
                        "OM1",
                        "OM3",
                        "OM4",
                        "OM5",
                        "YF1",
                        "YF2",
                        "YF3",
                        "YF4",
                        "YF5",
                        "MF1",
                        "MF2",
                        "MF3",
                        "MF4",
                        "MF5",
                        "OF1",
                        "OF3",
                        "OF4",
                        "OF5"
)]

rownames(my.gonad) <- my.gonad$GeneName

#generate count matrix
matrix <- my.gonad[,c(2:27)]

# select TEs and sum the counts
TEs.age <- matrix[grepl("NotFur", rownames(matrix)), ]
TE_sums <- colSums(TEs.age)

# select genes and sum the counts
genes.age <- matrix[!grepl("NotFur", rownames(matrix)), ]
genes_sums <- colSums(genes.age)

#Generate Fractional TE proportions as a percentage
TE_percentages <- (TE_sums/(TE_sums + genes_sums))*100

#rearrange scores for plotting
aggregate_te_scores <- melt(TE_percentages)
aggregate_te_scores$name <- rownames(aggregate_te_scores)
aggregate_te_scores$name <- gsub("\\d", "", aggregate_te_scores$name)
colnames(aggregate_te_scores) <-  c("te_perc", "group")

#order for plotting
aggregate_te_scores$group <- factor(aggregate_te_scores$group , levels=c("YF", "MF", "OF", "YM", "MM", "OM"))

#get specific p-values for plotting
fem_only <- subset(aggregate_te_scores, grepl("F",aggregate_te_scores$group))
m_only   <- subset(aggregate_te_scores, !grepl("F",aggregate_te_scores$group))


####### Tests
f.ym <- wilcox.test (aggregate_te_scores$te_perc[aggregate_te_scores$group == "YF"],aggregate_te_scores$te_perc[aggregate_te_scores$group == "MF"])
f.mo <- wilcox.test (aggregate_te_scores$te_perc[aggregate_te_scores$group == "MF"],aggregate_te_scores$te_perc[aggregate_te_scores$group == "OF"])
f.yo <- wilcox.test (aggregate_te_scores$te_perc[aggregate_te_scores$group == "YF"],aggregate_te_scores$te_perc[aggregate_te_scores$group == "OF"])

m.ym <- wilcox.test (aggregate_te_scores$te_perc[aggregate_te_scores$group == "YM"],aggregate_te_scores$te_perc[aggregate_te_scores$group == "MM"])
m.mo <- wilcox.test (aggregate_te_scores$te_perc[aggregate_te_scores$group == "MM"],aggregate_te_scores$te_perc[aggregate_te_scores$group == "OM"])
m.yo <- wilcox.test (aggregate_te_scores$te_perc[aggregate_te_scores$group == "YM"],aggregate_te_scores$te_perc[aggregate_te_scores$group == "OM"])


my.boxplot.out <- paste0("./Results/3_Fractional_analysis/", my.outprefix,"_AGING_SEX_TE_Percentages.pdf")
pdf(my.boxplot.out, height = 5, width = 6)
boxplot(te_perc ~ group, data = aggregate_te_scores, 
        col = c("deeppink", "deeppink3", "deeppink4","deepskyblue", "deepskyblue3", "deepskyblue4"),
        ylim = c(0,15), las = 2, ylab = "Gonadal TE fraction (%)")
beeswarm::beeswarm(te_perc ~ group, data = aggregate_te_scores, add = T, pch = 16)
text(1.5, 12, signif(f.ym$p.value,2) )
text(2.5, 12, signif(f.mo$p.value,2) )
text(2  , 14, signif(f.yo$p.value,2) )

text(4.5, 12, signif(m.ym$p.value,2) )
text(5.5, 12, signif(m.mo$p.value,2) )
text(5  , 14, signif(m.yo$p.value,2) )
dev.off()

#######################
sink(file = paste("./Results/3_Fractional_analysis/", my.outprefix,"_session_Info.txt", sep =""))
sessionInfo()
sink()
```
