## Supplemental Code for "Dynamic regulation of gonadal transposon control across the lifespan of the naturally short-lived African turquoise killifish": README_TE_Analysis.txt

########################################################
   				README -- 4_TE_Analysis
########################################################

# All scripts ran with R version 4.1.2 (2021-11-01)

*** All R package versions indicated in cognate scripts ***
	
	1. Ovarian differential TE expression using LRT with DESeq2
		- 1_Ovary_TE_DGE.R
		
	2. Testicular differential TE expression using LRT with DESeq2
		- 2_Testes_TE_DGE.R
		
	3. Plotting TE counts as a fraction of transcriptomes
		- 3_Fractional_TE_Analysis.R
		
	
