## Supplemental Code for "Dynamic regulation of gonadal transposon control across the lifespan of the naturally short-lived African turquoise killifish": 1_Ovary_piRNA_DGE.R

```
# set working directory to directory containing piRNA count matrix
# setwd("/Users/bryanteefy/Dropbox/PIWI/CODE/5_piRNA_Analysis")
setwd("/Users/berenice/Dropbox/Manuscripts_and_Publications/2022/2022_Bryan_PIWI_manuscript/CODE/5_piRNA_Analysis")

#Map Female piRNAs to TEs and run LRT DGE

# R version 4.1.2 (2021-11-01)

#load libraries
library(DESeq2)                  # DESeq2_1.34.0
library('bitops')                # bitops_1.0-7
library(RColorBrewer)            # RColorBrewer_1.1-3
library(pheatmap)                # pheatmap_1.0.12
library(DEGreport)               # DEGreport_1.30.3 
library(tibble)                  # tibble_3.1.7
library(dplyr)                   # dplyr_1.0.9 
library(ggplot2)                 # ggplot2_3.3.6
library(ggpubr)                  # ggpubr_0.4.0 
library(ggplotify)               # ggplotify_0.1.0 
library(SummarizedExperiment)    # SummarizedExperiment_1.24.0

#load count matrix (bowtie mapping, counting over TEs, piRNA clusters, rRNA and genes with fractional counts by FeatureCounts)
my.gonad <- read.table("./Input/piRNA_count_matrix.txt", header = T)

#preprocess piRNA count matrix to remove rRNA counts and remove non-count data
my.gonad <- my.gonad[,c(1,7:32)]
my.gonad <- my.gonad[!grepl("rRNA", my.gonad$Geneid), ] 

#remove excess characters from colnames
colnames(my.gonad) <- gsub("[.][.][.]", "", colnames(my.gonad))
colnames(my.gonad) <- gsub("[.]bam", "", colnames(my.gonad))

#generate age variables for females
my.Age  <- c(rep("Young",5),rep("Middle",5),rep("Old",4))
rownames(my.gonad) <- my.gonad$Geneid

#retain only female data (select only columns with female samples)
my.gonad <- my.gonad[,c(1,14:27)]

# get the genes with no reads out as they create issues with DEseq2
my.good <- which(apply(my.gonad[,-1]>0, 1, sum) >= 6)
my.filtered.matrix <- my.gonad[my.good,-1]

#Prerequisite: round the counts to nearest integer (DEseq2 needs integers)
rounded_matrix <- round(my.filtered.matrix)

#set prefix
my.outprefix <- paste(Sys.Date(),"Ovary_LRT_piRNA",sep="_")

#set colors
colors <- c("deeppink", "deeppink3", "deeppink4")

################################################################################################################
# 2. DESeq2 LRT DGE on cleaned data

# design matrix
dataDesign = data.frame( row.names = colnames(rounded_matrix), 
                         age       = my.Age)

# get matrix using age as a modeling covariate
dds <- DESeqDataSetFromMatrix(countData = rounded_matrix,
                              colData   = dataDesign,
                              design    = ~ age)

# run DESeq normalizations and export results with LRT
dds_lrt <- DESeq(dds, test="LRT", reduced = ~ 1)

# Extract results
res_LRT <- results(dds_lrt)

# Create a tibble for LRT results
res_LRT_tb <- res_LRT %>%
  data.frame() %>%
  rownames_to_column(var="gene") %>% 
  as_tibble()

# create output with only TEs
#### FishTEDB TEs all have "NotFur" prefix in name, which helps filter them out
#### We keep the genes this far to make sure all normalizations take their abundance into account
#### Only TEs are analyzed from this point forward
res_LRT_tb.TEs <- res_LRT_tb[grepl("NotFur", res_LRT_tb$gene), ]

# write output of LRT DGE Analysis to text file
write.table(res_LRT_tb    , file = paste0("./Results/1_Ovary_piRNA_DGE/", my.outprefix,"_Ovarian_LRT_DGE_piRNA_Table_ALL.txt"), sep = "\t" , row.names = F, quote = F)
write.table(res_LRT_tb.TEs, file = paste0("./Results/1_Ovary_piRNA_DGE/", my.outprefix,"_Ovarian_LRT_DGE_Table_piRNATEONLY.txt"), sep = "\t" , row.names = F, quote = F)

# Subset to return genes/TEs with padj < 1E-6
sigLRT_genes <- res_LRT_tb %>% 
  filter(padj < 0.000001)

# Get number of significant genes/TEs
nrow(sigLRT_genes) # 723

# Subset results for faster cluster finding
clustering_sig_genes <- sigLRT_genes %>%
  arrange(padj)

# normalize expression values
tissue.cts <- getVarianceStabilizedData(dds_lrt)

# write out normalized count matrix for use in GO analyses
write.table(tissue.cts, file = paste0("./Results/1_Ovary_piRNA_DGE/", my.outprefix,"_Female_Normalized_DEseq2_piRNATEGenes_Count_Matrix.txt"), sep = "\t" , row.names = T, quote = F)

# Extract significant genes
sig_genes <- tissue.cts[clustering_sig_genes$gene, ]

#retain TEs only (remove all other entries)
sig_genes <- sig_genes[grepl("NotFur", rownames(sig_genes)), ]

# Use the `degPatterns` function from the 'DEGreport' package to show gene clusters across sample groups
clusters <- degPatterns(sig_genes, metadata = dataDesign, time = "age", col=NULL, minc = 1)
clusters$normalized$age <- clusters$normalized$merge
clusters$normalized$age <- factor(clusters$normalized$age, levels=c("one_groupYoung", "one_groupMiddle", "one_groupOld"))

my.clusters.out <- paste0("./Results/1_Ovary_piRNA_DGE/", my.outprefix,"AGING_grouped_clusters_FDR5.pdf")
pdf(my.clusters.out, onefile = F, width = 20)
degPlotCluster(clusters$normalized, "age", lines=TRUE, min_genes = 1, col = "age") + theme_bw() + scale_color_manual(values=colors[c(1:3)])
dev.off()

#put clusters into a dataframe
df_cluster <- clusters$df

#Extract the Group 1 genes
cluster_groups <- clusters$df
group1 <- clusters$df %>%
  filter(cluster == 1)

#Extract the Group 2 genes
group2 <- clusters$df %>%
  filter(cluster == 2)

#Extract the Group 3 genes
group3 <- clusters$df %>%
  filter(cluster == 3)

# write out files
write.table(group1[,1], file = paste0("./Results/1_Ovary_piRNA_DGE/", my.outprefix,"_Female_AGING_Group1_DGE_piRNA_TEs_FDR1e-6.txt"), sep = "\t" , row.names = F, quote = F)
write.table(group2[,1], file = paste0("./Results/1_Ovary_piRNA_DGE/", my.outprefix,"_Female_AGING_Group2_DGE_piRNA_TEs_FDR1e-6.txt"), sep = "\t" , row.names = F, quote = F)
write.table(group3[,1], file = paste0("./Results/1_Ovary_piRNA_DGE/", my.outprefix,"_Female_AGING_Group3_DGE_piRNA_TEs_FDR1e-6.txt"), sep = "\t" , row.names = F, quote = F)

# create merged table of LRT/clustering results
# non significant genes will have 'NA' after outer join operation
df_cluster$genes <- gsub(".", ":", df_cluster$genes, fixed = T) 
res_LRT.clust   <- merge(data.frame(res_LRT_tb.TEs), df_cluster, by.x = "gene", by.y = "genes", all.x = T)
write.table(res_LRT.clust, file = paste0("./Results/1_Ovary_piRNA_DGE/", my.outprefix,"_DESeq2_LRT_DGE_Table_ALL_piRNA_TEs_with_DGEPatterns_Clusters_FDR1e-6.txt"), sep = "\t" , row.names = F, quote = F)

######################################################################################
#3. Generate Heatmaps color coded by TE Family

#Create function that 1. corrects modified TE names 2. Specifies TE family 3. generates heatmap
heatmap_TE <- function (x) {
  gene_names <- x$genes
  gene_names <- gsub("\\.", ":", gene_names)
  test2 <- as.data.frame(gene_names)
  matrix <- as.data.frame(tissue.cts)
  matrix$gene_names <- rownames(matrix)
  matrix$gene_names <- gsub("-", ":", matrix$gene_names)
  matrix2 <- merge(matrix, test2, by = "gene_names")
  matrix3 <- matrix2[,c(2:ncol(matrix2))]
  num <- nrow(matrix3)
  new_names <- gsub(".+:", "", gene_names)
  #merge unclear and Unknown families
  new_names <- gsub("unclear", "Unknown", new_names)
  NNs <- as.character(factor(new_names))
  rownames(matrix3) <- paste0("row_", seq(nrow(matrix3)))
  annotation_row = data.frame(
    Family = factor(NNs)
  )
  rownames(annotation_row) = rownames(matrix3)
  NNz <- order(NNs)
  order_of_TEs <- paste0("row_", NNz)
  # create colors for each group
  annoCol<-list(Family=c(DNA="firebrick1", LINE="darkturquoise", LTR="cornflowerblue", SINE="blue", Unknown="gold"))
  hm <- pheatmap(matrix3[order_of_TEs,],
                 cluster_cols = F,
                 cluster_rows = T,
                 colorRampPalette(rev(c("#CC3333","#FF9999","#FFCCCC","white","#CCCCFF","#9999FF","#333399")))(50),
                 show_rownames = F, scale="row",
                 border = NA,cellheight = 1,
                 main = paste(length(order_of_TEs), "TEs"), cellwidth = 15,
                 annotation_row = annotation_row,
                 annotation_colors = annoCol)
  return(hm)
}

female_plots <- ggarrange(as.ggplot(heatmap_TE(group1)),
                          as.ggplot(heatmap_TE(group2)),
                          as.ggplot(heatmap_TE(group3)))

my.heatmap.out <- paste0("./Results/1_Ovary_piRNA_DGE/", my.outprefix,"AGING_Heatmap_FDR5.pdf")
pdf(my.heatmap.out, onefile = F, height = 20, width = 20)
female_plots
dev.off()

######################################################################################
# 4. Expression Correlation Heatmap (QC)

# restrict analysis to only genes, not TEs
piRNA_exp <- tissue.cts[grepl("NotFur", rownames(tissue.cts)), ]

# obtain a correlation matrix of the gene expression data
correlation <- as.matrix(cor(piRNA_exp, method = "spearman"))

# arrange data
age <- data.frame("AgeGroup" = c(rep("YF",5),rep("MF",5),rep("OF",4) ))
rownames(age) = colnames(correlation)
age_cols = list(AgeGroup = c(YF = "deeppink", MF = "deeppink3", OF = "deeppink4"))

# plot and export
my.heatmap.out <- paste0("./Results/1_Ovary_piRNA_DGE/", my.outprefix,"_AGING_Gene_Exp_Correlation_Heatmap.pdf")
pdf(my.heatmap.out)
pheatmap(rbind(correlation,rep(0.9,nrow(correlation))),
         cluster_cols = F,
         cluster_rows = F,
         colorRampPalette(rev(c("#CC3333","#FF9999","#FFCCCC","white")))(20),
         show_rownames = T, scale="none",
         main = "Ovary piRNA Expression Correlation Matrix", 
         cellwidth = 20, cellheight = 20, display_numbers = T, number_color = "white", fontsize_number = 6,
         annotation_col = age,
         annotation_colors = age_cols)
dev.off()

#######################
sink(file = paste("./Results/1_Ovary_piRNA_DGE/", my.outprefix,"_session_Info.txt", sep =""))
sessionInfo()
sink()
```
