## Supplemental Code for "Dynamic regulation of gonadal transposon control across the lifespan of the naturally short-lived African turquoise killifish": 4_piRNA_cognate_TE_scatterplots_v2.R

```
# set working directory to directory containing piRNA count matrix
# setwd("/Users/bryanteefy/Dropbox/PIWI/CODE/5_piRNA_Analysis")
setwd("/Users/berenice/Dropbox/Manuscripts_and_Publications/2022/2022_Bryan_PIWI_manuscript/CODE/5_piRNA_Analysis")

#Plot Correlation between piRNAs and cognate TEs

#load libraries
library(DESeq2) # DESeq2_1.34.0
library(ggpubr) # ggpubr_0.4.0 

#########################################
#Generate normalized piRNA count matrix

#set prefix
my.outprefix <- paste(Sys.Date(),"TE_scatterplots",sep="_")

#load piRNA count matrix and preprocess
piRNA_counts <- read.table("./Input/piRNA_count_matrix.txt", header = T)
piRNA_counts <- piRNA_counts[,c(1,7:32)]
piRNA_counts <- piRNA_counts[!grepl("rRNA", piRNA_counts$Geneid), ] 
colnames(piRNA_counts) <- gsub("[.][.][.]", "", colnames(piRNA_counts))
colnames(piRNA_counts) <- gsub("[.]bam", "", colnames(piRNA_counts))

#load mRNA count matrix and preprocess
# load count matrix
mRNA_counts <- read.table("./Input/Gene_TE_Counts.cntTable", header = T)
mRNA_counts <- mRNA_counts[,c("GeneName",
                              "YM1","YM2","YM3","YM5",
                              "MM1","MM2","MM3","MM5",
                              "OM1","OM3","OM4","OM5",
                              "YF1","YF2","YF3","YF4","YF5",
                              "MF1","MF2","MF3","MF4","MF5",
                              "OF1","OF3","OF4","OF5")]

#Define biological variables
my.Sex  <- c(rep("M",12),rep("F",14))
my.Age  <- c(rep(5,4),rep(10,4),rep(15,4),rep(5,5),rep(10,5),rep(15,4))
groups <- c("YM","MM","OM","YF","MF","OF")
cols = c("deepskyblue","deepskyblue3","deepskyblue4","deeppink","deeppink3","deeppink4")

#define count normalization function and group median obtainment
obtain_medians <- function(x, agevar = my.Age, sexvar = my.Sex) {
  # format for processing
  rownames(x) <- x[,1]
  
  # get the genes with no reads out
  my.good <- which(apply(x[,-1]>0, 1, sum) >= 6) # see deseq2 vignette, need to remove too low genes
  my.filtered.matrix <- x[my.good,-1] 
  
  # Prerequisite: round the counts to nearest integer
  rounded_matrix <- round(my.filtered.matrix)
  
  # design matrix
  dataDesign = data.frame( row.names = colnames(rounded_matrix), 
                           age       = agevar,
                           sex       = sexvar)
  
  # get matrix using age as a modeling covariate
  dds <- DESeqDataSetFromMatrix(countData = rounded_matrix,
                                colData = dataDesign,
                                design = ~ age + sex)
  
  # run DESeq normalizations and export results
  dds.deseq <- DESeq(dds)
  
  # normalize expression values
  tissue.cts <- getVarianceStabilizedData(dds.deseq)
  
  #retain only TE sequences
  tissue.cts <- as.data.frame(tissue.cts[grepl("NotFur", rownames(tissue.cts)),])
  
  #obtain TE IDs
  TEs <- as.data.frame(rownames(tissue.cts))
  colnames(TEs) <- "ID"
  
  #define for loop to take median values per group
  for( i in 1: length(groups)) {
    a <- tissue.cts[ ,grepl(groups[i],names(tissue.cts))]
    b <- as.data.frame(apply(a, 1, median))
    TEs[,(i+1)] <- b
    colnames(TEs)[(i+1)] <- paste0(groups[i], "_median")
  }
  return(TEs)
}

#get median values, update names, then merge both dataframes
piRNA_medians            <- obtain_medians(piRNA_counts)
colnames(piRNA_medians) <- gsub("median", "piRNA_median", colnames(piRNA_medians))

mRNA_medians <- obtain_medians(mRNA_counts)

all_median_values <- merge(mRNA_medians, piRNA_medians, by = "ID")

#define scatterplot function
scatter <- function(a, b, c, d) {
  ggscatter(all_median_values, x = a, y = b,
            color = cols[d], size = 1, # Points color, shape and size
            add.params = list(color = "black", fill = NA), # Customize reg. line
            conf.int = TRUE, # Add confidence interval
            cor.coef = TRUE, # Add correlation coefficient. see ?stat_cor
            cor.coeff.args = list(method = "spearman", label.x = 6.5, label.sep = "\n"),  ### Spearman
            main = c, xlab = "Normalized piRNA Counts", ylab = "Normalized TE Counts")
}

#execute scatterplot function
YM <- scatter("YM_piRNA_median", "YM_median", "Young Testes"       , 1)
MM <- scatter("MM_piRNA_median", "MM_median", "Middle-Aged Testes" , 2)
OM <- scatter("OM_piRNA_median", "OM_median", "Old Testes"         , 3)
YF <- scatter("YF_piRNA_median", "YF_median", "Young Ovaries"      , 4)
MF <- scatter("MF_piRNA_median", "MF_median", "Middle-Aged Ovaries", 5)
OF <- scatter("OF_piRNA_median", "OF_median", "Old Ovaries"        , 6)

#plot and export
my.scatter.out <- paste0("./Results/4_piRNA_cognate_TE_scatterplots/", my.outprefix,"_piRNA_TE_cognate_scatterplot_SPEARMAN.pdf")
pdf(my.scatter.out, height = 10, width = 15)
ggarrange(YF,MF,OF,YM,MM,OM)
dev.off()

#######################
sink(file = paste("./Results/4_piRNA_cognate_TE_scatterplots/", my.outprefix,"_session_Info.txt", sep =""))
sessionInfo()
sink()
```
