## Supplemental Code for "Dynamic regulation of gonadal transposon control across the lifespan of the naturally short-lived African turquoise killifish": 5_length_and_nuc_dist_scripts_v2.R

```
# set working directory to directory containing piRNA count matrix
# setwd("/Users/bryanteefy/Dropbox/PIWI/CODE/5_piRNA_Analysis")
setwd("/Users/berenice/Dropbox/Manuscripts_and_Publications/2022/2022_Bryan_PIWI_manuscript/CODE/5_piRNA_Analysis")

# R version 4.1.2 (2021-11-01)

#length and nucleotide distribution plot generation

#load libraries
library(Biostrings) # Biostrings_2.62.0 
require(ggplot2)    # ggplot2_3.3.6
require(ggseqlogo)  # ggseqlogo_0.1
library(reshape2)   # reshape2_1.4.4

####################################
#1. Generate length distributions

#set outprefix
my.outprefix <- paste(Sys.Date(),"piRNA",sep="_")

#load list of file names
L = list.files(path = "./Input/piRNA_length_data/female_lengths", pattern=".txt", recursive = T)

names <- gsub("_piRNA_length.txt", "", L)

DFs = lapply(L, function(x) {
  data <- read.table(paste0("./Input/piRNA_length_data/female_lengths/", x), comment.char = "")
  colnames(data) <- c("freq", "length")
  return(data)
})

combo.f = Reduce(function(...) merge(..., by = "length", all=T), DFs)
colnames(combo.f) <- c("length", names)
combo.f$sums.f <- rowSums(combo.f[,c(2:15)])

female_length <- ggplot(data=combo.f, aes(x=length, y=sums.f, group=1)) +
  geom_line()+
  geom_point() + ggtitle("Seq Dist")

#save female logo plot
my.pca.out <- paste0("./Results/5_length_and_nuc_dist_scripts/", my.outprefix,"female_length_plot.pdf")
pdf(my.pca.out)
female_length
dev.off()

#repeat with males

#load list of file names
L = list.files(path = "./Input/piRNA_length_data/male_lengths", pattern=".txt", recursive = T)

names <- gsub("_piRNA_length.txt", "", L)

DFs = lapply(L, function(x) {
  data <- read.table(paste0("./Input/piRNA_length_data/male_lengths/", x), comment.char = "")
  colnames(data) <- c("freq", "length")
  return(data)
})

combo.m = Reduce(function(...) merge(..., by = "length", all=T), DFs)
colnames(combo.m) <- c("length", names)
combo.m$sums.m <- rowSums(combo.m[,c(2:13)])

male_length <- ggplot(data=combo.m, aes(x=length, y=sums.m, group=1)) +
  geom_line()+
  geom_point() + ggtitle("Seq Dist")

#save male length plot
my.pca.out <- paste0("./Results/5_length_and_nuc_dist_scripts/", my.outprefix,"male_length_plot.pdf")
pdf(my.pca.out)
male_length
dev.off()

#combine both samples
all <- merge(combo.f[,c(1,16)], combo.m[,c(1,14)], by = "length")

#set up a per million scaling factor for comparability of library sizes across sexes
# normalize to 1000 for readability
all$Female <- all$sums.f/((sum(all$sums.f))/1e3)
all$Male   <- all$sums.m/((sum(all$sums.m))/1e3)
all        <- all[,c(1,4,5)]

#melt to adapt for plotting
data <- melt(all, id = "length")
colnames(data)[c(2,3)] <- c("Sex","reads_per_1e3")

#plot dual length plot
my.plot.out <- paste0("./Results/5_length_and_nuc_dist_scripts/", my.outprefix, "_dual_length_plot_by_1e3.pdf")
pdf(my.plot.out)
ggplot(data=data, aes(x=length, y=reads_per_1e3, group=Sex)) +
  geom_line(aes(color=Sex)) +
  geom_point(aes(color=Sex))+ theme_bw() +
  scale_colour_manual(name= "Sex", values = c("deeppink", "deepskyblue"))+
  ggtitle("piRNA Size Distributions")  +
  theme(plot.title = element_text(hjust = 0.5)) +
  xlab("Length (nt)") + ylab("Reads per 1e3")
dev.off()

##############################################
#2. Nucleotide Distribution Plots

#load trimmed fasta sequences without headers
female_sequences <- read.table("./Input/total_fasta_females_replaced.fasta")

#make position weight matrix
pfm_fm <- consensusMatrix(female_sequences$V1)

#make logo plot
fem_plot <- ggseqlogo(pfm_fm, method = 'bit', seq_type = "rna") + ggtitle("Females")

#remove large modified fasta file
rm(female_sequences)  

#save female logo plot
my.logo.out <- paste0("./Results/5_length_and_nuc_dist_scripts/", my.outprefix,"female_logo_plot.pdf")
pdf(my.logo.out, height = 2.5, width = 5)
fem_plot
dev.off()


#load trimmed fasta sequences without headers
male_sequences <- read.table("./Input/total_fasta_males_replaced.fasta")

#make position weight matrix
pfm_m <- consensusMatrix(male_sequences$V1)

#make logo plot
male_plot <- ggseqlogo(pfm_m, method = 'bit', seq_type = "rna") + ggtitle("Males")

#remove large modified fasta file
rm(male_sequences)  

#save male logo plot
my.logo.out <- paste0("./Results/5_length_and_nuc_dist_scripts/", my.outprefix,"male_logo_plot.pdf")
pdf(my.logo.out, height = 2.5, width = 5)
male_plot
dev.off()

#######################
sink(file = paste("./Results/5_length_and_nuc_dist_scripts/", my.outprefix,"_session_Info.txt", sep =""))
sessionInfo()
sink()
```
