## Supplemental Code for "Dynamic regulation of gonadal transposon control across the lifespan of the naturally short-lived African turquoise killifish": 6_ppr_mbr.R

```
# set working directory to directory containing piRNA count matrix
# setwd("/Users/bryanteefy/Dropbox/PIWI/CODE/5_piRNA_Analysis")
setwd("/Users/berenice/Dropbox/Manuscripts_and_Publications/2022/2022_Bryan_PIWI_manuscript/CODE/5_piRNA_Analysis")

# R version 4.1.2 (2021-11-01)

#Grouped ping-pong analysis using the output of Ping-Pong Meter

#this is the script to generate all of the ppm-mrb plots
#this script gives us the ppr-mbr score which is the ping-pong instances per million bootstapped reads
#the input is the number of instances of a given overlap per million read pairs

#############################################################

#set prefix
my.outprefix <- paste(Sys.Date(),"ppr_mbr_analysis",sep="_")

#load list of file names
L = list.files(path = "./Input/ppr_mbr", pattern=".txt", recursive = T)
names <- gsub("_ppm.txt", "", L)

#this function keeps only the instances of 10 bp overlaps per million read pairs
DFs = lapply(L, function(x) {
  data <- read.table(paste0("./Input/ppr_mbr/", x), skip = 2, nrows = 100)
  data <- data[,c(1,38)]
  colnames(data) <- c("rep", "score")
  score <- median(data$score)
  return(score)
})

#run for loop to get median ppm per replicate
med_ppm <- data.frame()
for(i in 1:length(L)) {
  total <- cbind(names[i], DFs[[i]])
  med_ppm <- rbind(med_ppm, total)
}
colnames(med_ppm) <- c("group", "score")
med_ppm$score <- as.numeric(med_ppm$score)

#divide ppr-mbr score by 10e5 for plotting
med_ppm$score <- med_ppm$score/100000

#remove rep ID to plot by group ID only
med_ppm$group <- gsub('[[:digit:]]+', '', med_ppm$group)

#order for plotting
med_ppm$group <- factor(med_ppm$group , levels=c("YF", "MF", "OF", "YM", "MM", "OM"))

#get specific p-values for plotting
fem_only <- subset(med_ppm, grepl("F",med_ppm$group))
m_only   <- subset(med_ppm, !grepl("F",med_ppm$group))

####### Tests
f.ym <- wilcox.test (fem_only$score[fem_only$group == "YF"],fem_only$score[fem_only$group == "MF"])
f.mo <- wilcox.test (fem_only$score[fem_only$group == "MF"],fem_only$score[fem_only$group == "OF"])
f.yo <- wilcox.test (fem_only$score[fem_only$group == "YF"],fem_only$score[fem_only$group == "OF"])

m.ym <- wilcox.test (med_ppm$score[med_ppm$group == "YM"],med_ppm$score[med_ppm$group == "MM"])
m.mo <- wilcox.test (med_ppm$score[med_ppm$group == "MM"],med_ppm$score[med_ppm$group == "OM"])
m.yo <- wilcox.test (med_ppm$score[med_ppm$group == "YM"],med_ppm$score[med_ppm$group == "OM"])

my.boxplot.out <- paste0("./Results/6_ppr_mbr/", my.outprefix,"_AGING_SEX_TE_Percentages.pdf")
pdf(my.boxplot.out, height = 5, width = 5)
boxplot(score ~ group, data = med_ppm, 
        col = c("deeppink", "deeppink3", "deeppink4","deepskyblue", "deepskyblue3", "deepskyblue4"),
        ylim = c(0,6), las = 2, ylab = "Median x10e5 ppr-mbr (100 bootstraps")
beeswarm::beeswarm(score ~ group, data = med_ppm, add = T, pch = 16)
text(1.5, 5.5, signif(f.ym$p.value,2) )
text(2.5, 5.5, signif(f.mo$p.value,2) )
text(2  , 6, signif(f.yo$p.value,2) )

text(4.5, 5.5, signif(m.ym$p.value,2) )
text(5.5, 5.5, signif(m.mo$p.value,2) )
text(5  , 6, signif(m.yo$p.value,2) )
dev.off()

#######################
sink(file = paste("./Results/6_ppr_mbr/", my.outprefix,"_session_Info.txt", sep =""))
sessionInfo()
sink()
```
