## Supplemental Code for "Dynamic regulation of gonadal transposon control across the lifespan of the naturally short-lived African turquoise killifish": 7_locus_level_z10_analysis_v2.R

```
# set working directory to directory containing z10 data
# setwd("/Users/bryanteefy/Dropbox/PIWI/CODE/5_piRNA_Analysis")
setwd("/Users/berenice/Dropbox/Manuscripts_and_Publications/2022/2022_Bryan_PIWI_manuscript/CODE/5_piRNA_Analysis")

# R version 4.1.2 (2021-11-01)

#get z10 scores

#load libraries
library(purrr)         # purrr_0.3.4
library(DESeq2)        # DESeq2_1.34.0
library('bitops')      # bitops_1.0-7
library(pheatmap)      # pheatmap_1.0.12
library(DEGreport)     # DEGreport_1.30.3 
library(tibble)        # tibble_3.1.7
library(dplyr)         # dplyr_1.0.9
library(ggplotify)     # ggplotify_0.1.0
library(ggpubr)        # ggpubr_0.4.0

#set prefix
my.outprefix <- paste(Sys.Date(),"z10_analysis",sep="_")

colors <- c("deeppink", "deeppink3", "deeppink4","deepskyblue", "deepskyblue3", "deepskyblue4")

#load list of file names
L = list.files(path = "./Input/locus_level_z_scores", pattern=".txt", recursive = T)

#take basename
names <- gsub("_TEs_only_histos_reduced.txt", "", L)

#establish function to call locus level z-10 scores
DFs = lapply(L, function(x) {
  data <- read.table(paste0("./Input/locus_level_z_scores/", x), comment.char = "")
  colnames(data) <- c("chr", "pos", "freq")
  data <- data[!data$pos > 20,] #anything above 20 is noisy
  rownames(data) <- 1:nrow(data) #rename rows sequentially
  d = data.frame(a=rep(0, length(unique(data$chr))), z=rep(0,length(unique(data$chr)))) #initialize dataframe
  #define z10 function
  z_10_score <- function(w) {
    z <- (((w[,3][w[,2]==10 & w[,1]==a]) - mean(w[,3][w[,2]!=10 & w[,1]==a]))/(sd(w[,3][w[,2]!=10 & w[,1]==a])))
    return(z) # take the z10 score. i.e. the mean of overlap position 10 - mean of all other positions over the standard deviation of all other positions
  }
  for( i in seq(from = 1, to = length((unique(data$chr))), by = 1)) {
    y <- distinct(data[1])
    a <- y[i,]
    z <- z_10_score(data)
    d[i, ] = c(a, z)
  } # run the z10 function for each unique transposon 
  z_scores <- d # save output as z-scores
  z_scores$z <- as.numeric(z_scores$z) # make z-score column numerical
  colnames(z_scores) <- c("ID", "Z10_score") #rename columns 
  return(z_scores)
})
#There were 26 warnings (use warnings() to see them) DISREGARD, does not affect output

#merge all outputs into a single dataframe
z_10s = Reduce(function(...) merge(..., by = "ID", all=T), DFs) #There were 23 warnings (use warnings() to see them) DISREGARD, does not affect output
colnames(z_10s) <- c("ID", names)

#reorganize by age and sex, remove infinites and NAs
z10_scores <- z_10s
rownames(z10_scores) <- z10_scores$ID
z10_scores <- z10_scores[,c(2:27)]
z10_scores <- z10_scores[is.finite(rowSums(z10_scores)),]
z10_scores <- z10_scores[apply(is.na(z10_scores), 1, sum) == 0,]
z10_scores <- z10_scores[,c(23:26, 6:9, 14:17, 18:22, 1:5, 10:13)]

#import Z10 scores and take only ovary and testis data alone
fem_only <- z10_scores[,c(13:26)]
male_only <- z10_scores[,c(1:12)]

#define function to run anova over z10 matrices
anova_testing <- function(x) {
  
  #transpose and organize z10 scores for anova testing by age group
  z_matrix <- as.data.frame(t(x))
  z_matrix$group <- rownames(z_matrix)
  z_matrix <- z_matrix[,c(ncol(z_matrix),1:(ncol(z_matrix) -1))]
  z_matrix$group <- gsub("\\d", "", z_matrix$group)
  
  #run anova testing on z10 matrix. generate a list of anova tests for each TE transcript
  fit_aov <- function(col) {
    aov(col ~ group, data = z_matrix)
  }
  
  anovas <- map(z_matrix[, 2:ncol(z_matrix)], fit_aov)
  
  #extract p-values, adjust, and bind to TE name
  pvals <- sapply(anovas, function(x) summary(x)[[1]][["Pr(>F)"]][[1]])
  padj <- p.adjust(pvals, method = "BH", n = length(pvals))
  signif_table <- as.data.frame(colnames(z_matrix[,-1]))
  signif_table <- cbind(signif_table, padj)
  colnames(signif_table) <- c("TE", "padj")
  
  #retain only significant (p-value < 0.05) TEs
  sig <- subset(signif_table, signif_table$padj < 0.05)
  
  #extract significant TEs from the z10 score matrix
  sig_final <- x %>% filter(row.names(x) %in% sig$TE)
  
  #return
  return(sig_final)
}

# keep signifcant results by sex
fem_sig  <- anova_testing(fem_only)  # 109
male_sig <- anova_testing(male_only) # 83

##############################
#2. Run "LRT-like" analysis on significant female Z-scores

#make data design table
my.Age.fem  <- c(rep("Young",5),rep("Middle",5),rep("Old",4))
dataDesign.fem = data.frame( row.names = colnames(fem_sig), 
                             age = my.Age.fem)

# Use the `degPatterns` function from the 'DEGreport' package to show TE clusters across sample groups
clusters <- degPatterns(fem_sig, metadata = dataDesign.fem, time = "age", col=NULL, minc = 1)
clusters$normalized$age <- clusters$normalized$merge
clusters$normalized$age <- factor(clusters$normalized$age, levels=c("one_groupYoung", "one_groupMiddle", "one_groupOld"))

my.clusters.out <- paste0("./Results/7_locus_level_z10_analysis/", my.outprefix,"z10_grouped_clusters_FDR5_female.pdf")
pdf(my.clusters.out, onefile = F)
degPlotCluster(clusters$normalized, "age", lines=TRUE, min_genes = 1, col = "age") + theme_bw() + scale_color_manual(values=colors[c(1:3)])
dev.off()

#separate into groups
group1.f <- clusters$df %>%
  filter(cluster == 1)

group2.f <- clusters$df %>%
  filter(cluster == 2)

group3.f <- clusters$df %>%
  filter(cluster == 3)

group4.f <- clusters$df %>%
  filter(cluster == 4)

#rename TEs due to changes created by degPatterns
rename <- function(x) {
  x$genes <- gsub("\\.\\.", "::", x$genes)
  x$genes <- gsub("\\.", "-", x$genes)
  return(x)
}

group1.f <- rename(group1.f)
group2.f <- rename(group2.f)
group3.f <- rename(group3.f)
group4.f <- rename(group4.f)

# write out files
write.table(rbind(group1.f,group2.f,group3.f,group4.f), file = paste0("./Results/7_locus_level_z10_analysis/", my.outprefix,"_Female_AGING_ANOVA_Z10_piRNA_TEs.FDR5_withGroups.txt"), sep = "\t" , row.names = F, quote = F)


###############################################
#repeat with males

#make data design table
my.Age.m  <- c(rep("Young",4),rep("Middle",4),rep("Old",4))
dataDesign.m = data.frame( row.names = colnames(male_sig), 
                           age       = my.Age.m)

# Use the `degPatterns` function from the 'DEGreport' package to show TE clusters across sample groups
clusters <- degPatterns(male_sig, metadata = dataDesign.m, time = "age", col=NULL, minc = 1)
clusters$normalized$age <- clusters$normalized$merge
clusters$normalized$age <- factor(clusters$normalized$age, levels=c("one_groupYoung", "one_groupMiddle", "one_groupOld"))

my.clusters.out <- paste0("./Results/7_locus_level_z10_analysis/", my.outprefix,"z10_grouped_clusters_FDR5_male.pdf")
pdf(my.clusters.out, onefile = F)
degPlotCluster(clusters$normalized, "age", lines=TRUE, min_genes = 1, col = "age") + theme_bw() + scale_color_manual(values=colors[c(4:6)])
dev.off()

#separate into groups
group1.m <- clusters$df %>%
  filter(cluster == 1)

group2.m <- clusters$df %>%
  filter(cluster == 2)

#rename
group1.m <- rename(group1.m)
group2.m <- rename(group2.m)

# write out files
write.table(rbind(group1.m,group2.m), file = paste0("./Results/7_locus_level_z10_analysis/", my.outprefix,"_Male_AGING_ANOVA_Z10_piRNA_TEs.FDR5_withGroups.txt"), sep = "\t" , row.names = F, quote = F)


###############################################
#3. Plot all Z10 Data as Heatmaps

#Define pre-processing function to adapt names and color by TE family (females)
plot_hm <- function (x,y) {
  gene_names <- x[,1]
  GN <- as.data.frame(gene_names)
  matrix <- as.data.frame(y)
  matrix$gene_names <- rownames(matrix)
  matrix2 <- merge(matrix, GN, by = "gene_names")
  matrix3 <- matrix2[,c(2:ncol(matrix2))]
  num <- nrow(matrix3)
  new_names <- gsub("unclear", "Unknown", gene_names)
  new_names <- gsub("\\:\\:.+", "", new_names)
  new_names <- gsub("?.+_", "", new_names)
  NNs <- as.character(factor(new_names))
  rownames(matrix3) <- paste0("row_", seq(nrow(matrix3)))
  annotation_row = data.frame(
    Family = factor(NNs)
  )
  rownames(annotation_row) = rownames(matrix3)
  NNz <- order(NNs)
  TE_order <- paste0("row_", NNz)
  # create colors for each group
  annoCol<-list(Family=c(DNA="firebrick1", LINE="darkturquoise", LTR="cornflowerblue", SINE="blue", Unknown="gold"))
  hm <- pheatmap(matrix3[TE_order,],
                 cluster_cols = F,
                 cluster_rows = T,
                 colorRampPalette(rev(c("#CC3333","#FF9999","#FFCCCC","white","#CCCCFF","#9999FF","#333399")))(50),
                 show_rownames = F, scale="row",
                 border = NA,cellheight = 1,
                 main = paste("Genes Up with Age", length(TE_order), "Genes"), cellwidth = 15,
                 annotation_row = annotation_row,
                 annotation_colors = annoCol)
  return(hm)
}

my.heatmap.out <- paste0("./Results/7_locus_level_z10_analysis/", my.outprefix,"Female_z10_heatmap.pdf")
pdf(my.heatmap.out, onefile = F, height = 10, width = 10)
ggarrange(as.ggplot(plot_hm(group1.f, fem_sig)),
          as.ggplot(plot_hm(group2.f, fem_sig)),
          as.ggplot(plot_hm(group3.f, fem_sig)),
          as.ggplot(plot_hm(group4.f, fem_sig)))
dev.off()

my.heatmap.out <- paste0("./Results/7_locus_level_z10_analysis/", my.outprefix,"Male_z10_heatmap.pdf")
pdf(my.heatmap.out, onefile = F, height = 10, width = 10)
ggarrange(as.ggplot(plot_hm(group1.m, male_sig)),
          as.ggplot(plot_hm(group2.m, male_sig)))
dev.off()

###############################################
#4. Plot all Z10 Data as Grouped Boxplots

#take median values for each replicate
replicate_medians <- as.data.frame(apply(z10_scores, 2, median))
replicate_medians$group <- rownames(replicate_medians)
colnames(replicate_medians) <- c("z_score", "group")

#define groups
groups <- c("YM","MM","OM","YF","MF","OF")

#remove rep ID to plot by group ID only
replicate_medians$group <- gsub('\\d', '', replicate_medians$group)

#order for plotting
replicate_medians$group <- factor(replicate_medians$group , levels=c("YF", "MF", "OF", "YM", "MM", "OM"))

#get specific p-values for plotting
fem_only <- subset(replicate_medians, grepl("F",replicate_medians$group))
m_only   <- subset(replicate_medians, !grepl("F",replicate_medians$group))

####### Tests
f.ym <- wilcox.test (replicate_medians$z_score[replicate_medians$group == "YF"],replicate_medians$z_score[replicate_medians$group == "MF"])
f.mo <- wilcox.test (replicate_medians$z_score[replicate_medians$group == "MF"],replicate_medians$z_score[replicate_medians$group == "OF"])
f.yo <- wilcox.test (replicate_medians$z_score[replicate_medians$group == "YF"],replicate_medians$z_score[replicate_medians$group == "OF"])

m.ym <- wilcox.test (replicate_medians$z_score[replicate_medians$group == "YM"],replicate_medians$z_score[replicate_medians$group == "MM"])
m.mo <- wilcox.test (replicate_medians$z_score[replicate_medians$group == "MM"],replicate_medians$z_score[replicate_medians$group == "OM"])
m.yo <- wilcox.test (replicate_medians$z_score[replicate_medians$group == "YM"],replicate_medians$z_score[replicate_medians$group == "OM"])

my.boxplot.out <- paste0("./Results/7_locus_level_z10_analysis/", my.outprefix,"_AGING_SEX_TE_z10_boxplots.pdf")
pdf(my.boxplot.out, height = 5, width = 5)
boxplot(z_score ~ group, data = replicate_medians, 
        col = c("deeppink", "deeppink3", "deeppink4","deepskyblue", "deepskyblue3", "deepskyblue4"),
        ylim = c(0,18), las = 2, ylab = "Z10 Scores")
beeswarm::beeswarm(z_score ~ group, data = replicate_medians, add = T, pch = 16)
text(1.5, 15, signif(f.ym$p.value,2) )
text(2.5, 15, signif(f.mo$p.value,2) )
text(2  , 16, signif(f.yo$p.value,2) )

text(4.5, 15, signif(m.ym$p.value,2) )
text(5.5, 15, signif(m.mo$p.value,2) )
text(5  , 16, signif(m.yo$p.value,2) )
dev.off()

#######################
sink(file = paste("./Results/7_locus_level_z10_analysis/", my.outprefix,"_session_Info.txt", sep =""))
sessionInfo()
sink()
```
