## Supplemental Code for "Dynamic regulation of gonadal transposon control across the lifespan of the naturally short-lived African turquoise killifish": 8_histogram_plots.R

```
# set working directory to directory containing z10 data
# setwd("/Users/bryanteefy/Dropbox/PIWI/CODE/5_piRNA_Analysis")
setwd("/Users/berenice/Dropbox/Manuscripts_and_Publications/2022/2022_Bryan_PIWI_manuscript/CODE/5_piRNA_Analysis")

# R version 4.1.2 (2021-11-01)

#Histograms of z-scores
#Warning: This script is time-consuming/memory intensive to run
# 
#load libraries
library(purrr)        # purrr_0.3.4  
library('bitops')     # bitops_1.0-7   
library(DEGreport)    # DEGreport_1.30.3 
library(tibble)       # tibble_3.1.7
library(dplyr)        # dplyr_1.0.9         
library(ggplot2)      # ggplot2_3.3.6 
library(ggplotify)    # ggplotify_0.1.0 
library(ggpubr)       # ggpubr_0.4.0 

#set output prefix
my.outprefix <- paste(Sys.Date(),"z10_analysis",sep="_")

#load list of file names
L = list.files(path = "./Input/locus_level_z_scores", pattern=".txt", recursive = T)

#store names for later assignment
names <- gsub("_TEs_only_histos_reduced.txt", "", L)

#initialize empty dataframes and lists
df1 <-as.data.frame(matrix(nrow=20,ncol=1))
lsZ<-list()

#Define and run function to generate Z-scores for every overlap length from 1-20 nts
DFs = lapply(L, function(x) {
  for(j in 1:20){
    data <- read.table(paste0("./Input/locus_level_z_scores/", x), comment.char = "")
    colnames(data) <- c("chr", "pos", "freq")
    data <- data[!data$pos > 20,] #anything above 20 is noisy
    rownames(data) <- 1:nrow(data) #rename rows sequentially
    d = data.frame(a=rep(0, length(unique(data$chr))), z=rep(0,length(unique(data$chr)))) #initialize dataframe
    #define z10 function
    z_j_score <- function(w) {
      z <- (((w[,3][w[,2]==j & w[,1]==a]) - mean(w[,3][w[,2]!=j & w[,1]==a]))/(sd(w[,3][w[,2]!=j & w[,1]==a])))
      return(z) # take the zj score. i.e. the mean of overlap position j - mean of all other positions over the standard deviation of all other positions
    }
    for( i in seq(from = 1, to = length((unique(data$chr))), by = 1)) {
      y <- distinct(data[1])
      a <- y[i,]
      z <- z_j_score(data)
      d[i, ] = c(a, z)
    } # run the z10 function for each unique transposon 
    z_scores <- d
    z_scores$z <- as.numeric(z_scores$z)
    colnames(z_scores) <- c("ID", "Zj_score")
    lsZ[[j]] <- z_scores #save as list element for each round
  }
  lsZ <- lapply(lsZ,na.omit) #remove any rows with NAs
  for(k in 1:20){
    df1[k,] <- median(lsZ[[k]]$Zj_score) #take the median z[pos] score and put it into a dataframe
  }
  return(df1) #return median z[pos] score by replicate
})

# There were 50 or more warnings (use warnings() to see the first 50) # NA induced by coercion, DISREGARD

#set names
for(i in seq(from = 1, to = length(DFs))) {
  colnames(DFs[[i]]) <- names[i]
}

#sort
MFs <- cbind(DFs[[1]] ,DFs[[2]] ,DFs[[3]] ,DFs[[4]],DFs[[5]])
MMs <- cbind(DFs[[6]] ,DFs[[7]] ,DFs[[8]] ,DFs[[9]] )
OFs <- cbind(DFs[[10]],DFs[[11]],DFs[[12]],DFs[[13]])
OMs <- cbind(DFs[[14]],DFs[[15]],DFs[[16]],DFs[[17]])
YFs <- cbind(DFs[[18]],DFs[[19]],DFs[[20]],DFs[[21]],DFs[[22]])
YMs <- cbind(DFs[[23]],DFs[[24]],DFs[[25]],DFs[[26]])

#define plotting preparation function by taking the median Z-score value for each groip
plot_ready <- function(x){
  x$Median <- apply(x, 1, median)
  x$Overlap <- as.numeric(rownames(x))
  y <- x[,c("Median", "Overlap")]
  return(y)
}

#generate histograms

#define colors
colors <- c("deeppink", "deeppink3", "deeppink4", "deepskyblue", "deepskyblue3", "deepskyblue4")

#define plotting function
plot_hist <- function(a,b,c) {
  ggplot(a, aes(x=Overlap, y=Median)) + 
    geom_bar(stat = "identity", fill = colors[b]) +
    ggtitle(c) + xlab("piRNA 5' Overlap value")+ylab("Z-Score Median Value") +
    ylim(-1,12) + xlim(0,21) +
    theme(text = element_text(size=20),
          axis.text.x = element_text(angle=0, hjust=0.5), 
          plot.title = element_text(color="black", size=20, hjust = 0.5))+ theme_bw()
}

#plot and export
my.hist.out <- paste0("./Results/8_histogram_plots/", my.outprefix,"z10_histograms.pdf")
pdf(my.hist.out, onefile = F, height = 15, width = 20)
ggarrange(plot_hist(plot_ready(YFs), 1, "Young Ovaries Z-Scores"),
          plot_hist(plot_ready(MFs), 2, "Middle-Aged Ovaries Z-Scores"),
          plot_hist(plot_ready(OFs), 3, "Old Ovaries Z-Scores"),
          plot_hist(plot_ready(YMs), 4, "Young Testes Z-Scores"),
          plot_hist(plot_ready(MMs), 5, "Middle-Aged Testes Z-Scores"),
          plot_hist(plot_ready(OMs), 6, "Old Testes Z-Scores"))
dev.off()

#######################
sink(file = paste0("./Results/8_histogram_plots/", my.outprefix,"_session_Info.txt", sep =""))
sessionInfo()
sink()
```
