## Supplemental Code for "Dynamic regulation of gonadal transposon control across the lifespan of the naturally short-lived African turquoise killifish": 9_compare_TE_piRNAs.R

```
# setwd("/Users/bryanteefy/Dropbox/PIWI/CODE/5_piRNA_Analysis")
setwd("/Users/berenice/Dropbox/Manuscripts_and_Publications/2022/2022_Bryan_PIWI_manuscript/CODE/5_piRNA_Analysis")

# R version 4.1.2 (2021-11-01)

# compare DE TEs and DE TE-targeting piRNAs

# load libraries
library(Vennerable)        # Vennerable_3.1.0.9000

####################################
#1. Read DE results

#set outprefix
my.outprefix <- paste(Sys.Date(),"piRNA_TE_comparison_ovary",sep="_")

# load results of LRT TE analysis (ovary)
ovary.TE.LRT  <- read.table("../4_TE_Analysis/Results/1_Ovary_TE_DGE/2022-07-13_Ovary_TE_analysis_LRT_DESeq2_LRT_TE_DGE_Table_ALL_with_DGEPatterns_Clusters_FDR1e-6.txt", header = T)

# load results of LRT piRNA analysis (ovary)
ovary.pi.LRT  <- read.table("./Results/1_Ovary_piRNA_DGE/2022-07-27_Ovary_LRT_piRNA_DESeq2_LRT_DGE_Table_ALL_piRNA_TEs_with_DGEPatterns_Clusters_FDR1e-6.txt", header = T)

# compare overlap of TEs up at mid-age [pattern b, group 1 in output] with piRNA down at mid-age [pattern a, group 1 in output]
ovary.TE.LRT.up_Mage  <- ovary.TE.LRT[which(ovary.TE.LRT$cluster == 1),]
ovary.pi.LRT.dwn_Mage <- ovary.pi.LRT[which(ovary.pi.LRT$cluster == 1),]


##### Venn Diagrams using Vennerable
my.ov.DE.MidAge <- list("TE_up_MidAge"         = ovary.TE.LRT.up_Mage$gene   ,
                        "piRNA_down_MidAge"    = ovary.pi.LRT.dwn_Mage$gene  )
my.Venn <- Venn(my.ov.DE.MidAge)

# Note: the universe/background will be TEs detected in both analysis
my.uni <- intersect(ovary.TE.LRT$gene, ovary.pi.LRT$gene)


# without scaling
pdf(paste0("./Results/9_DE_piRNA_TE_comparison_Venn/",my.outprefix,"_Venn.pdf"))
plot(my.Venn, doWeights=F, show = list(FaceText = "weight", SetLabels = TRUE, Faces = FALSE))
dev.off()

# test enrichment (overlap greater than by chance)
fisher.test(matrix(c(36, 
                     163,
                     272,
                     length(my.uni)-36-163-272),
                   2,2), 
            alternative = 'greater') # test if overlap is larger than expected
# Fisher's Exact Test for Count Data
# data:  matrix(c(36, 163, 272, length(my.uni) - 36 - 163 - 272), 2, 2)
# p-value = 0.009075
# alternative hypothesis: true odds ratio is greater than 1
# 95 percent confidence interval:
#  1.162773      Inf
# sample estimates:
# odds ratio 
#   1.644688 

# get the consistently regulated TEs
my.consistent <- intersect(ovary.TE.LRT.up_Mage$gene, ovary.pi.LRT.dwn_Mage$gene)

# extract TE family 
new_names <- gsub(".+:", "", my.consistent)

# make unclear merge with unknown
new_names[new_names == "unclear"] <- "Unknown"

# output pie chart of composition
inter.tab <- signif(table(new_names)/length(new_names)*100,2)
names(inter.tab) <- paste( names(inter.tab) , inter.tab, "%")

pdf(paste0("./Results/9_DE_piRNA_TE_comparison_Venn/",my.outprefix,"_pie_chart_of_intersection.pdf"))
pie(inter.tab, col = c("firebrick1", "darkturquoise", "cornflowerblue", "gold"))
dev.off()


######################
sink(file = paste("./Results/9_DE_piRNA_TE_comparison_Venn/", my.outprefix,"_session_Info.txt", sep =""))
sessionInfo()
sink()
```
