## Supplemental Code for "Dynamic regulation of gonadal transposon control across the lifespan of the naturally short-lived African turquoise killifish": README_5_piRNA_Analysis.txt

########################################################
            README -- 5_piRNA_Analysis
########################################################

# All scripts ran with R version 4.1.2 (2021-11-01)

*** All R package versions indicated in cognate scripts ***
	
	1. Ovarian differential piRNA expression over TEs using LRT with DESeq2
		- 1_Ovary_piRNA_DGE.R
		
	2. Testicular differential piRNA expression over TEs using LRT with DESeq2
		- 2_Testes_piRNA_DGE.R	
		
	3. Generates stacked barplots showing piRNA cluster and genomic TE composition by family 
		- 3_TE_repeat_content_in_clusters.R
		
	4. Plot correlation between piRNAs and cognate TEs 
		- 4_piRNA_cognate_TE_scatterplots.R
		
	5. Generate plots of piRNA length and nucleotide distribution
		- 5_length_and_nuc_dist_scripts.R
		
	6. Analysis of ping-pong via PPmeter output 
		- 6_ppm_mbr.R
		
	7. TE consensus level Z10-score generation and ANOVA-based DGE 
		- 7_locus_level_z10_analysis.R
		
	8. Generate Z-score Histograms
		- 8_histogram_plots.R
	
	9. Compare Ovarian Differentially Expressed TEs and piRNAs
		- 9_compare_TE_piRNAs.R
