## Supplemental Code for "Dynamic regulation of gonadal transposon control across the lifespan of the naturally short-lived African turquoise killifish": 1_GRZ_lifespan_v2.R

```
#set working directory
 setwd("/Users/bryanteefy/Dropbox/PIWI/CODE/6_Animal_Analysis")
#setwd("/Users/berenice/Dropbox/Manuscripts_and_Publications/2022/2022_Bryan_PIWI_manuscript/CODE/6_Animal_Analysis/")

#Generate Fecundity and Lifespan Plots

# R version 4.1.2 (2021-11-01)

#load libraries
library(beeswarm)  # beeswarm_0.4.0
library(survival)  # survival_3.3-1
library(ranger)    # ranger_0.14.1
library(ggplot2)   # ggplot2_3.3.6
library(dplyr)     # dplyr_1.0.9  
library(ggfortify) # ggfortify_0.4.14

################################
# 1. Import required data

#Import Killfish lifespan data
span <-read.table("./Input/Killifish_Lifespans.txt", header = T)

#########################
# 2. Plot Lifespan

#obtain median lifespan
Fem_med_life <- median(span[span$sex == 'Female', 'time']) # 26.55 weeks
Male_med_life <- median(span[span$sex == 'Male', 'time']) # 19.4 weeks

#fit using the survival package
km_trt_fit <- survfit(Surv(time, status) ~ sex, data=span)

#running lg rank test
surv.test <- survdiff(Surv(time, status) ~ sex, span)

surv.test
#survdiff(formula = Surv(time, status) ~ sex, data = span)
#
#N Observed Expected (O-E)^2/E (O-E)^2/V
#sex=Female 24       24     35.1      3.51      9.61
#sex=Male   37       37     25.9      4.76      9.61
#
#Chisq= 9.6  on 1 degrees of freedom, p= 0.002

#make into a dataframe
modf <- fortify(km_trt_fit)

#change "stata" back to "sex"
colnames(modf)[9] <- "sex"

#remove excess data
modf <- modf[,c(1,5,9)]

#format for plotting
modf[58,] <- c(0, 1, "Female")
modf[59,] <- c(0, 1, "Male")
modf$time <- as.numeric(modf$time)
modf$surv <- as.numeric(modf$surv)
modf$sex  <- as.character(modf$sex)

#plot and export
my.curve.out <- paste0("./Results/", Sys.Date(),"_GRZ_survival_curve.pdf")
pdf(my.curve.out, onefile = F, width = 10)
ggplot(modf, aes(x = time, y = surv, group=sex)) + 
  geom_point(aes(color=sex)) +
  geom_line(aes(color=sex)) +
  ggtitle("GRZ Survival Curve") + xlab("Time (Weeks)")+ylab("Fraction Surviving") +
  scale_color_manual(values=c("deeppink", "deepskyblue")) +theme_bw() +
  xlim(0, 50)+
  theme(text = element_text(size=20),
        axis.text.x = element_text(angle=0, hjust=0.5), 
        plot.title = element_text(color="black", size=20, hjust = 0.5))
dev.off()


#######################
sink(file = paste0("./Results/", Sys.Date(),"_session_Info.txt", sep =""))
sessionInfo()
sink()
```
