## Supplemental Code for "Dynamic regulation of gonadal transposon control across the lifespan of the naturally short-lived African turquoise killifish": 2_Oocyte_Diameter_v2.R

```
#set working directory
setwd("/Users/bryanteefy/Dropbox/PIWI/CODE/6_Animal_Analysis")

#Generate Oocyte Diameter Plots

# R version 4.2.2 (2022-10-31)

#load libraries
library(ggplot2)
library(reshape2)
library(ggpubr)

#make into function
input_tables <- c("./Input/Katelyn_Final_Oocyte_Diameter.txt", "./Input/Alan_Final_Oocyte_Diameter.txt", "./Input/Berenice_Final_Oocyte_Diameter.txt", "./Input/Bryan_Final_Oocyte_Diameter.txt")

run_plot <- function (x,y) {
  
  #import quantification of oocyte sizes
  oo.d <- read.table(input_tables[x], header = T, fill = T)
  oo.d.m <- melt(oo.d)
  oo.d.m <- na.omit(oo.d.m)
  colnames(oo.d.m) <- c("sample", "oocyte_diameter")
  
  #rearrange scores for plotting
  oo.d.m$sample <- gsub("\\d", "", oo.d.m$sample)
  
  #order for plotting
  oo.d.m$sample <- factor(oo.d.m$sample , levels=c("YF", "MF", "OF"))
  
  #run Kolmogorov-smirnov test between young and middle-aged, and middle-aged and old samples
  YvM.pval <- (ks.test(subset(oo.d.m, oo.d.m$sample == "YF")[,2], subset(oo.d.m, oo.d.m$sample == "MF")[,2]))$p.value
  MvO.pval <- (ks.test(subset(oo.d.m, oo.d.m$sample == "MF")[,2], subset(oo.d.m, oo.d.m$sample == "OF")[,2]))$p.value
  
  #plot oocyte diameter data by group with KS test appended onto plots
  plot <- ggplot(oo.d.m, aes(x=sample, y=oocyte_diameter)) + 
    geom_boxplot()+ 
    ggtitle(y) + theme_bw() + 
    geom_text(x=1.5, y=800, label=YvM.pval, family = "Helvetica") +
    geom_text(x=2.5, y=800, label=MvO.pval, family = "Helvetica")
  
  return(plot)
  
}

Katelyn <- run_plot(1, "Observer 1")
Alan <- run_plot(2, "Observer 2")
Berenice <- run_plot(3, "Observer 3")
Bryan <- run_plot(4, "Observer 4")

my.plot <- paste0("./Results/2_oocyte_diameter/", Sys.Date(),"_GRZ_oocyte_diameter_KS_test.pdf")
pdf(my.plot)
ggarrange(Katelyn, Alan, Berenice, Bryan) 
dev.off()
```
