## Supplemental Code for "Dynamic regulation of gonadal transposon control across the lifespan of the naturally short-lived African turquoise killifish": README_Animal_Analysis_v2.txt

########################################################
              README -- 6_Animal_Analysis
########################################################

# Scripts ran with R version 4.1.2 (2021-11-01)

*** All R package versions indicated in cognate scripts ***

	1. Generates lifespan curves for both sexes
	   Runs statistical analysis for each as well
		- 1_GRZ_lifespan_v2.R
	
	2. Generates oocyte diameter distributions and comparisons:
		- 2_Oocyte_Diameter_v2.R
