## Supplemental Code for "Dynamic regulation of gonadal transposon control across the lifespan of the naturally short-lived African turquoise killifish": 1_Run_BLAST_zebra_killi_v2.txt

#1) Download  Danio rerio Protein Sequence (FASTA) from Ensembl
# Sequences were downloaded October 31, 2022, corresponding to Ensembl release 108 from

# http://uswest.ensembl.org/info/data/ftp/index.html/
# 
# To download human protein sequences, perform the following steps:
# 
# 	a) Select to "BioMart" from the menu at the top of the webpage
# 	b) CHOOSE DATABASE: Select "Ensembl Genes [version]"
# 			-Note: this publication used Ensembl release 108
# 	c) CHOOSE DATASET: Select "Zebrafish genes (GRCz11)" (filters: protein coding)
# 	d) Select "Attributes" on the side menu
# 	e) Select the "Sequences" option
# 	f) Under the newly populated "SEQUENCES" menu, ensure that "Peptide" is selected
# 	g) Go to header, make sure only are selected "Protein stable ID version, Gene name"
# 	h) Select "Results" from the top left corner menu
# 	i) Ensure that protein sequences are present in FASTA format 
# 		ex. >ENSG00000001617|ENSG00000001617.12|ENST00000002829|ENST00000002829.8
# 			MLVAGLLLWASLL...*
# 	j) Export file as a "Compressed file (.gz)" in FASTA format
# 	k) Unzip file locally and save this file as "Zebra_Ens108_prot.fasta".
	
# Parse Gene Names
cat Zebra_Ens108_prot.fasta | grep '>' |  perl -lane '$_ =~ s/>//g; $_ =~ s/\|/\t/g; print $_' > Zebrafish_Gene_Name_ENSDAR_table_Ens108.txt


#2) Generate a local BLAST database of the fish Protein Sequences [BLAST 2.10.0+]
# makeblastdb -in GCA_014300015.1_MPIA_NFZ_2.0_protein.faa -dbtype prot -out Killi_prot_db 

#3) BLAST Zebrafish Protein Sequences against the Killifish Protein Database
blastp -db Killi_prot_db -query Zebra_Ens108_prot.fasta                  -evalue 1e-5 -outfmt 6 -max_target_seqs 1 -subject_besthit > 2022-11-01_best_hits_zebra_to_killi_eval_1e-5.txt

# extract best killifish hit for each zebrafish protein
cat 2022-11-01_best_hits_zebra_to_killi_eval_1e-5.txt | sort -k1,1 -k12,12nr -k11,11n | sort -u -k1,1 > 2022-11-01_best_hits_zebra_to_killi_eval_1e-5.UNIQUE.txt
