## Supplemental Code for "Dynamic regulation of gonadal transposon control across the lifespan of the naturally short-lived African turquoise killifish": 2_Parse_killifish_zebrafish_Blast_results_v2.R

```
setwd('/Users/berenice/Dropbox/Manuscripts_and_Publications/2022/2022_Bryan_PIWI_manuscript_SUBMITTED/CODE/7_Ovary_deconvolution_test')
options(stringsAsFactors = F)

# Gene ID to Gene Name conversion tables
killi.gnames <- read.csv("./Input/Parsed_GCA_014300015.1_MPIA_NFZ_2.0_Gene_to_Protein_ID_conversion.txt", header = T, sep = "\t")

# Read best BLAST results
z2k <- read.csv("./Input/2022-11-01_best_hits_zebra_to_killi_eval_1e-5.UNIQUE.txt", header = F, sep = "\t")
z2k$Danrer_GeneName <- unlist(lapply(strsplit(z2k$V1,"|", fixed = T),'[',2))

# save unique gene pairs
z2k.cl <- unique(z2k[,c("Danrer_GeneName","V2")]) # 28186 hits
colnames(z2k.cl)[2] <- "Killi_Best"

# Add in killi gene accession
z2k.cl.ann     <- merge(z2k.cl, killi.gnames, by.x = "Killi_Best" , by.y = "protein_id")

save(z2k.cl.ann, file = paste0(Sys.Date(), "_homology_table_killifish_zebrafish.RData" ))
```
