## Supplemental Code for "Dynamic regulation of gonadal transposon control across the lifespan of the naturally short-lived African turquoise killifish": 3_Deconvolute_ovarian_transcriptome_Collapse_v2.R

```
setwd('/Users/berenice/Dropbox/Manuscripts_and_Publications/2022/2022_Bryan_PIWI_manuscript_SUBMITTED/CODE/7_Ovary_deconvolution_test')
options(stringsAsFactors = F)

# Load necessary libraries
library('GenomicFeatures') # GenomicFeatures_1.46.5
library('Seurat')          # Seurat_4.2.0
library('granulator')      # granulator_1.2.0
library('pheatmap')        # pheatmap_1.0.12
library('beeswarm')        # beeswarm_0.4.0     

# load accessory functions
source('3_FUNCTIONS_Deconvolution.R')

########################################################################
# 1. Generate tpm tables for bulk ovary RNAseq 

# TPM normalization is required by Granulator package
# http://bioconductor.org/packages/release/bioc/vignettes/granulator/inst/doc/granulator.html

# Extract transcript length information using GenomicFeatures package
# https://rdrr.io/bioc/GenomicFeatures/man/makeTxDbFromGFF.html
killi.txdb           <- makeTxDbFromGFF(file="./Input/GCA_014300015.1_MPIA_NFZ_2.0_genomic.gtf")
killi.trscpt.lengths <- transcriptLengths(killi.txdb, with.cds_len=T,with.utr5_len=T, with.utr3_len=T)

# Retain only longest transcript for each gene
killi.tx.lg.flt <- aggregate(killi.trscpt.lengths[,c("nexon","tx_len")], by = list(killi.trscpt.lengths$gene_id), FUN = "max")
colnames(killi.tx.lg.flt)[1] <- "gene_id"

# load bulk count matrix
my.gonad <- read.table("./Input/Gene_TE_Counts.cntTable", header = T)

# reorganize by sex and age
my.gonad <- my.gonad[,c("GeneName",
                        "YM1",
                        "YM2",
                        "YM3",
                        "YM5",
                        "MM1",
                        "MM2",
                        "MM3",
                        "MM5",
                        "OM1",
                        "OM3",
                        "OM4",
                        "OM5",
                        "YF1",
                        "YF2",
                        "YF3",
                        "YF4",
                        "YF5",
                        "MF1",
                        "MF2",
                        "MF3",
                        "MF4",
                        "MF5",
                        "OF1",
                        "OF3",
                        "OF4",
                        "OF5"
)]


# generate age variables for females
my.Age             <- c(rep("Young",5),rep("Middle",5),rep("Old",4))

# retain only female data 
my.gonad <- my.gonad[,c(1,14:27)]

# Generate a merged matrix with counts and transcript length
my.gonad.lgth           <- merge(killi.tx.lg.flt[,c("gene_id","tx_len")], my.gonad, by.x = "gene_id", by.y = "GeneName")
rownames(my.gonad.lgth) <- my.gonad.lgth$gene_id

# Generate tpm values
my.ov.tpm <- tpm3(my.gonad.lgth[,-c(1:2)], my.gonad.lgth$tx_len)

write.table(my.ov.tpm, file = paste0(Sys.Date(),"_Killifish_ovary_TPM.txt"), sep = "\t", quote = F)

########################################################################
# 2. Load and process scRNA seq on zebrafish ovary
# https://singlecell.broadinstitute.org/single_cell/study/SCP928/40dpf-ovary-all-cells

######## A. Read in the Seurat and metadata  ########
# Create Seurat object using expression data and published cell annotation
my.meta.data.all              <- read.csv("./Input/Ovary_scRNAseq/zx124_40com_ssportal_meta.txt"    , "\t", header = T)
my.meta.data.all              <- my.meta.data.all[-1,] # remove line with redundant header info
rownames(my.meta.data.all)    <- my.meta.data.all$NAME
colnames(my.meta.data.all)[5] <- "Comp_annot"

# Load Seurat object
load('./Input/Ovary_scRNAseq/zx124_40com_final_orig.robj')
zx124_40com_final_orig
# An object of class Seurat 
# 43536 features across 25089 samples within 2 assays 
# Active assay: SCT (21766 features, 3000 variable features)
# 1 other assay present: RNA
# 2 dimensional reductions calculated: pca, umap

# Add metadata
my.zebra.ov  <- AddMetaData(zx124_40com_final_orig, my.meta.data.all, col.name = NULL) # from Figure 1 annotation

# Examine distribution
table(my.zebra.ov@meta.data$Comp_annot)

# Create higher level (less subclassses) annotation
my.zebra.ov@meta.data$Clean <- NA
my.zebra.ov@meta.data$Clean[my.zebra.ov@meta.data$Comp_annot %in% c("GSC+GC_Pro") ] <- "GSPCs"
my.zebra.ov@meta.data$Clean[my.zebra.ov@meta.data$Comp_annot %in% c("Early_Meio","Late_Meio","Meio") ] <- "Meiotic"
my.zebra.ov@meta.data$Clean[my.zebra.ov@meta.data$Comp_annot %in% c("Early_OO_1","Early_OO_2","Early_OO_3") ] <- "Oocytes"
my.zebra.ov@meta.data$Clean[my.zebra.ov@meta.data$Comp_annot %in% c("Follicle_1","Follicle_2","Follicle_lhx9") ] <- "Follicle"
my.zebra.ov@meta.data$Clean[my.zebra.ov@meta.data$Comp_annot %in% c("Macrophage") ]  <- "Macrophage"
my.zebra.ov@meta.data$Clean[my.zebra.ov@meta.data$Comp_annot %in% c("Neutrophils") ] <- "Neutrophils"
my.zebra.ov@meta.data$Clean[my.zebra.ov@meta.data$Comp_annot %in% c("NK-like") ]     <- "NK-like"
my.zebra.ov@meta.data$Clean[my.zebra.ov@meta.data$Comp_annot %in% c("Theca") ]       <- "Theca"
my.zebra.ov@meta.data$Clean[my.zebra.ov@meta.data$Comp_annot %in% c("Vasculature") ] <- "Vasculature"
my.zebra.ov@meta.data$Clean[grep("Stromal_",my.zebra.ov@meta.data$Comp_annot) ]      <- "Stromal"
my.zebra.ov@meta.data$Clean[grep("Unknown_",my.zebra.ov@meta.data$Comp_annot) ]      <- "Unknown"

table(my.zebra.ov@meta.data$Comp_annot,my.zebra.ov@meta.data$Clean)
table(my.zebra.ov@meta.data$Clean)
# Follicle       GSPCs  Macrophage     Meiotic Neutrophils     NK-like     Oocytes     Stromal       Theca 
# 4346        3332         556        4315         380        1549        3041        5525         891 
# Unknown Vasculature 
# 268         886 

### subset to remove "unknown"
Idents(object = my.zebra.ov) <- "Clean"
my.zebra.ov.cl <- subset(my.zebra.ov, idents = "Unknown", invert = TRUE)

# Make global reference pseudobulks
zebra.ov.comp.pb.v2   <- data.frame(AggregateExpression(my.zebra.ov.cl   , group.by = "Clean", slot = "counts", assay = "RNA")$RNA)

# Generate tpm values for pure pseudobulk
zebra.ov.comp.pb.v2.tpm <- tpmUMI(zebra.ov.comp.pb.v2)


######## B. Make ovary pseudobulks to test deconvolution accuracy  ########

# Get "True" Percentages
global.props <- table(my.zebra.ov.cl@meta.data$Clean)/length(my.zebra.ov.cl@meta.data$Clean[!is.na(my.zebra.ov.cl@meta.data$Clean)])

#### simulate mixtures
set.seed(123456789)

# Create full pseudobulk for each cell type
zebra.ov.pure.pb  <- data.frame(AggregateExpression(my.zebra.ov.cl, group.by = "Clean", slot = "counts", assay = "RNA")$RNA)

# Make up random "shuffles" of ground truth percentages (first 1 is ground truth)
my.pb.freqs <- data.frame("PB_1" = as.numeric(global.props),
                          "PB_2" = sample(as.numeric(global.props)),
                          "PB_3" = sample(as.numeric(global.props)),
                          "PB_4" = sample(as.numeric(global.props)),
                          "PB_5" = sample(as.numeric(global.props)))
rownames(my.pb.freqs) <- names(global.props)

# Make ovarian bulks using "real" proportions for a weighted mean
my.ov.sim.1 <- get_wt_mean(zebra.ov.pure.pb, my.pb.freqs$PB_1)
my.ov.sim.2 <- get_wt_mean(zebra.ov.pure.pb, my.pb.freqs$PB_2)
my.ov.sim.3 <- get_wt_mean(zebra.ov.pure.pb, my.pb.freqs$PB_3)
my.ov.sim.4 <- get_wt_mean(zebra.ov.pure.pb, my.pb.freqs$PB_4)
my.ov.sim.5 <- get_wt_mean(zebra.ov.pure.pb, my.pb.freqs$PB_5)


# Make dataframe
Zebra.ov.PBmix <- round(data.frame('PB_1'= my.ov.sim.1, 
                                   'PB_2'= my.ov.sim.2, 
                                   'PB_3'= my.ov.sim.3,
                                   'PB_4'= my.ov.sim.4,
                                   'PB_5'= my.ov.sim.5))
rownames(Zebra.ov.PBmix) <- rownames(zebra.ov.pure.pb)

# Generate tpm values for fake bulks
Zebra.ov.PBmix.tpm <- tpmUMI(Zebra.ov.PBmix)


########################################################################
# 3. Killify zebrafish ovary pseudobulk

# load orthology table
load('./Input/2022-11-02_homology_table_killifish_zebrafish.RData')
# z2k.cl.ann

# "killify" zebrafish pure cells and mixes
k.zebra.pure.ov.v2.tmp <- merge(unique(z2k.cl.ann[,2:3]), zebra.ov.comp.pb.v2.tpm, by.x = "Danrer_GeneName", by.y = "row.names")
k.zebra.ov.mix.tmp  <- merge(unique(z2k.cl.ann[,2:3]), Zebra.ov.PBmix.tpm  , by.x = "Danrer_GeneName", by.y = "row.names")

# For some genes, there are 2 zebrafish genes but only one killi homolog
# aggregate the tpms

k.zebra.pure.ov.v2 <- aggregate(k.zebra.pure.ov.v2.tmp[,-c(1:2)], by = list(k.zebra.pure.ov.v2.tmp$gene_id), sum)
k.zebra.ov.mix     <- aggregate(k.zebra.ov.mix.tmp [,-c(1:2)]   , by = list(k.zebra.ov.mix.tmp$gene_id ), sum)

rownames(k.zebra.pure.ov.v2) <- k.zebra.pure.ov.v2$Group.1
rownames(k.zebra.ov.mix ) <- k.zebra.ov.mix$Group.1

k.zebra.pure.ov.v2 <-  k.zebra.pure.ov.v2[,-1]
k.zebra.ov.mix     <-  k.zebra.ov.mix [,-1]

# save R objects
save(my.ov.tpm, k.zebra.pure.ov.v2, k.zebra.ov.mix, file = paste0(Sys.Date(),"_TPM_normalized_matrices_forGranulator.RData") )
save(my.pb.freqs, file = paste0(Sys.Date(),"_Proportions_for_PBmix.RData") )


#
# write.table(my.ov.tpm[rownames(k.zebra.pure.ov),], file = paste0(Sys.Date(),"_Killifish_ovary_TPM_zebgenesonly.txt"), sep = "\t", quote = F)


########################################################################
# 4. Try deconvolution using granulator
# http://bioconductor.org/packages/release/bioc/vignettes/granulator/inst/doc/granulator.html#bulk-pbmcs-rna-seq

# load('2022-11-02_TPM_normalized_matrices.RData')
# load('2022-11-02_Proportions_for_PBmix.RData')

# plot signature matrix similarity using spearman correlation
my.cors <- cor(k.zebra.pure.ov.v2, method ="spearman")

pdf(paste0(Sys.Date(),"_Heatmap_cell_type_ovary_correlation_spearman_collapse.pdf"))
pheatmap(my.cors)
dev.off()

# piwil1 - KAF7199819.1/G4P62_016517
pdf(paste0(Sys.Date(),"_Heatmap_piwil1_tpm_Ovary_PB.pdf"))
pheatmap(k.zebra.pure.ov.v2["G4P62_016517",], cluster_rows = F, cellheight = 20)
dev.off()


#########################################################
########   A. Benchmark on artificial PB mixes   ########

# Deconvolution of pseudo bulk RNA-seq data from zebrafish ovary to determine performance
PBmix.decon <- deconvolute(m = as.matrix(k.zebra.ov.mix), sigMatrix = as.matrix(k.zebra.pure.ov.v2))

# We can plot the estimated cell type proportions with the function plot_proportions().
# Notice that while the sum of cell types proportions cannot exceed 100%, for some methods part of the bulk RNA-seq signal remains unassigned.
# plot cell type proportions for svr model on ABIS_S0 reference profile
plot_proportions(deconvoluted = PBmix.decon, method = 'svr')

PBmix.bench <- benchmark(deconvoluted = PBmix.decon, ground_truth = as.matrix(t(my.pb.freqs)) )

# print metrics
PBmix.bench$rank
#    signature  method mean_pcc mean_ccc mean_adj.r2 mean_rmse
# 1      sig1     svr   0.9005   0.0124      0.7533    0.0240
# 2      sig1    nnls   0.8932   0.0182      0.7367    0.0283
# 3      sig1     ols   0.8932   0.0182      0.7367    0.0283
# 4      sig1   qprog   0.8932   0.0182      0.7367    0.0283
# 5      sig1 qprogwc   0.8932   0.0182      0.7367    0.0283
# 6      sig1     rls   0.8932   0.0182      0.7367    0.0283
# 7      sig1 dtangle   0.8816   0.0084      0.7144    0.0186

pdf(paste0(Sys.Date(),"_dothcart_granulator_algorithms_performance_on_PBmix_benchmarking.pdf"), height = 4, width = 4)
dotchart(PBmix.bench$rank$mean_pcc, labels = PBmix.bench$rank$method, xlim = c(0.5,1), pch = 16, xlab = "Mean PCC")
dev.off()

write.table(PBmix.bench$rank, file = paste0(Sys.Date(),"_granulator_deconvolution_algorithms_performance_on_PBmix.txt"), sep = "\t", quote = F, row.names = F)

# plot pearson correlation between predictions and true proportions
pdf(paste0(Sys.Date(),"_Deconvolution_algorithms_performance_on_PBmix_PCC_metric.pdf"), height = 5, width = 6)
plot_benchmark(benchmarked = PBmix.bench, metric = 'pcc')
dev.off()

# Extract regression/correlation plots for top 2 methods
pdf(paste0(Sys.Date(),"_realvspred_cell_type_ovary_correlation_SVR.pdf"), height = 5, width = 8)
plot_regress(benchmarked = PBmix.bench, method = 'svr')
dev.off()

pdf(paste0(Sys.Date(),"_realvspred_cell_type_ovary_correlation_NNLS.pdf"), height = 5, width = 8)
plot_regress(benchmarked = PBmix.bench, method = 'nnls')
dev.off()


##############################################################
########   B. Run deconvolution on killi ovary data   ########

# Deconvolution of pseudo bulk RNA-seq data using top 2 algorithms from benchmark
killi.decon <- deconvolute(m = as.matrix(my.ov.tpm), sigMatrix = as.matrix(k.zebra.pure.ov.v2), c("nnls","svr"))
### yields negative cell proportions
# will try CIBERSORT

nnls.res <- killi.decon$proportions$nnls_sig1
nnls.res$Age <- factor(c(rep("Middle",5),rep("Old",4),rep("Young",5)), levels = c("Young","Middle","Old") )

svr.res <- killi.decon$proportions$svr_sig1
svr.res$Age <- factor(c(rep("Middle",5),rep("Old",4),rep("Young",5)), levels = c("Young","Middle","Old") )

# GSPCs and Meiotic OOcytes have the highest piwil1 expression 
# Is the decreased expression due to decreased expression?
nnls.res$PiwiHi <-  nnls.res$GSPCs + nnls.res$Meiotic
svr.res$PiwiHi  <-  svr.res$GSPCs  + svr.res$Meiotic

wilcox.test(nnls.res$PiwiHi[nnls.res$Age == "Young"] , nnls.res$PiwiHi[nnls.res$Age == "Middle"]) # p-value = 0.08969
wilcox.test(nnls.res$PiwiHi[nnls.res$Age == "Middle"], nnls.res$PiwiHi[nnls.res$Age == "Old"]   ) # p-value = 0.01945

wilcox.test(svr.res$PiwiHi[svr.res$Age == "Young"]  , svr.res$PiwiHi[svr.res$Age == "Middle"])   # p-value = 0.07491
wilcox.test(svr.res$PiwiHi[svr.res$Age == "Middle"] , svr.res$PiwiHi[svr.res$Age == "Old"]   )   # p-value = 0.02684

pdf(paste0(Sys.Date(),"_killi_ovary_piwiHi_cells_NNLS.pdf"), height = 4, width = 5)
beeswarm(Meiotic + GSPCs ~ Age, data = nnls.res, pch = 16, ylim = c(0,60), main = "NNLS deconvolution", pwcol = c(rep("deeppink3",5),rep("deeppink4",4),rep("deeppink",5)) )
text(1.5, 55, "ns")
text(2.5, 55, "*")
dev.off()

pdf(paste0(Sys.Date(),"_killi_ovary_piwiHi_cells_SVR.pdf"), height = 4, width = 5)
beeswarm(Meiotic + GSPCs ~ Age, data = svr.res , pch = 16, ylim = c(0,60), main = "SVR deconvolution", pwcol = c(rep("deeppink3",5),rep("deeppink4",4),rep("deeppink",5)) )
text(1.5, 55, "ns")
text(2.5, 55, "*")
dev.off()

#######################
sink(file = paste0(Sys.Date(),"_Granulator_prep_session_Info.txt", sep =""))
sessionInfo()
sink()
```
