## Supplemental Code for "Dynamic regulation of gonadal transposon control across the lifespan of the naturally short-lived African turquoise killifish": 3_FUNCTIONS_Deconvolution.R

```
# deconvolution analysis accessory functions

################################################
# Function to generate tpm values
# https://support.bioconductor.org/p/91218/
tpm3 <- function(counts,len) {
  x <- counts/len
  return(t(t(x)*1e6/colSums(x)))
}


################################################
# Function to make weighted means of expression
get_wt_mean <- function(exp.matrix,my.weights){
  my.result <- rep(0,dim(exp.matrix)[1])
  for (i in 1: dim(exp.matrix)[1]) {
    my.result[i] <- weighted.mean(exp.matrix[i,], my.weights)
  }
  return(my.result)
}


################################################
# Function to generate "fake" tpm values from UMI 
# (length correction of 100 based on 10xGenomics UMI read)
tpmUMI <- function(counts) {
  x <- counts/100
  return(t(t(x)*1e6/colSums(x)))
}
```
