## Supplemental Code for "Dynamic regulation of gonadal transposon control across the lifespan of the naturally short-lived African turquoise killifish": README_Deconvolution.txt

########################################################
              README -- 7_Ovary_deconvolution_test
########################################################

# Scripts ran with R version 4.1.2 (2021-11-01)

*** All R package versions indicated in cognate scripts ***

	1. Identify best BLAST hit of Zebrafish/Killifish proteins
		- 1_Run_BLAST_zebra_killi_v2.txt

	2. Generate Zebrafish/Killifish conversion tables, parse protein/gene conversions across species
		- 2_Parse_killifish_zebrafish_Blast_results_v2.R
	
	3. Run transcriptome deconvolution using Granulator
		Use Zebrafish scRNAseq ovarian dataset (https://singlecell.broadinstitute.org/single_cell/study/SCP928/40dpf-ovary-all-cells)
		- 3_Deconvolute_ovarian_transcriptome_Collapse_v2.R
		- 3_FUNCTIONS_Deconvolution.R
	
