## Supplemental Code for "Dynamic regulation of gonadal transposon control across the lifespan of the naturally short-lived African turquoise killifish": Complete_Repository_README.txt

# Killifish_reproductive_aging_resource

## README ## 

Code for the manuscript:
Bryan B. Teefy, Ari Adler, Alan Xu, Katelyn Hsu, Param Priya Singh and Bérénice A. Benayoun
"Dynamic regulation of gonadal homeostasis across the lifespan of the naturally short-lived African turquoise killifish"
(biorXiv preprint at: https://www.biorxiv.org/content/10.1101/2022.09.02.506427v1)

The code is broadly arranged by the RNA species being analyzed:

	# 1_Gene_and_TE_Preprocessing                    : see README_Gene_and_TE_Preprocessing.txt
	# 2_piRNA_preprocessing				 : see README_piRNA_Preprocessing.txt
	# 3_Differential_Gene_Expression_Analysis	 : see README_DGE_Analysis.txt
	# 4_TE_Analysis       				 : see README_TE_Analysis.txt
	# 5_piRNA_Analysis				 : see README_piRNA_Analysis.txt
	# 6_Animal_Analysis      			 : see README_Animal_Analysis_v2.txt
	# 7_Ovary_deconvolution_test                     : see README_Deconvolution.txt
