## Supplemental Fig. S1 for "Dynamic regulation of gonadal transposon control across the lifespan of the naturally short-lived African turquoise killifish"

Figure S1

A

| Datasets generated in this study | Ovaries | Testes | Total | |
|---|---|---|---|---|
| mRNA-seq | 5,5,4 (14) | 4,4,4 (12) | 26 | 52 datasets |
| small RNA-seq | 5,5,4 (14) | 4,4,4 (12) | 26 | |

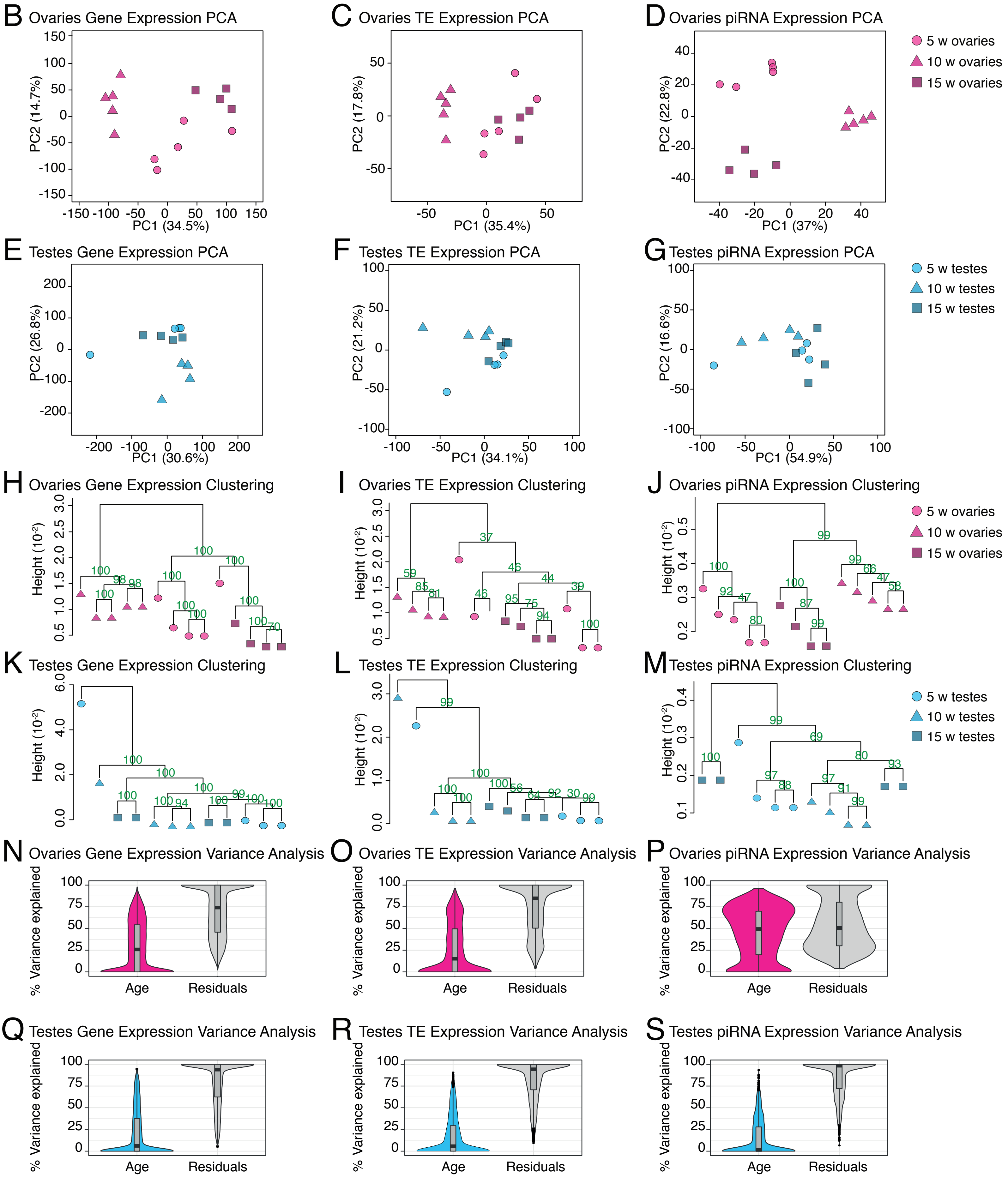
