## Supplementary figures and images for "Dynamic regulation of gonadal transposon control across the lifespan of the naturally short-lived African turquoise killifish"

### Supplemental Fig. S2

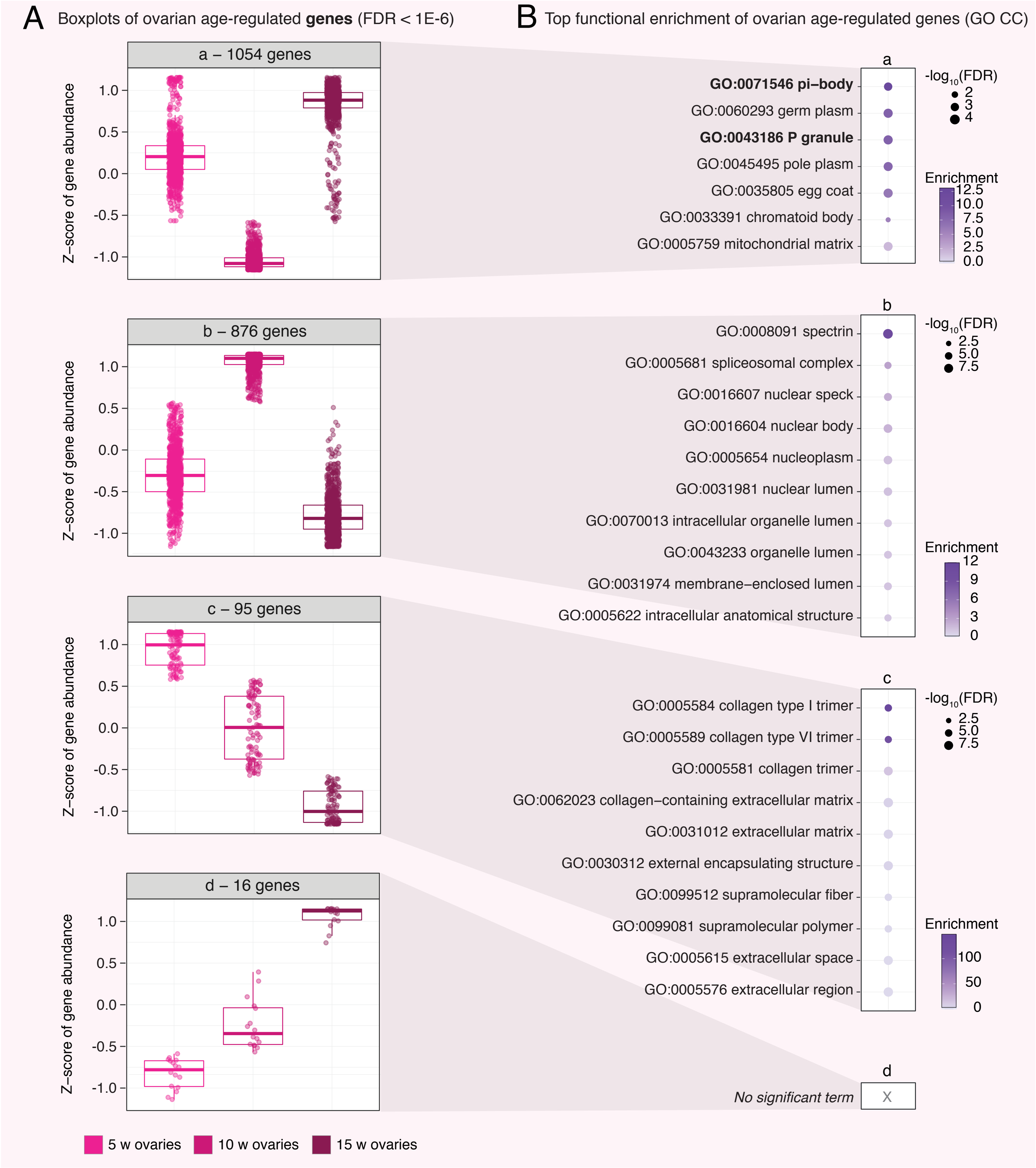

### Supplemental Fig. S3

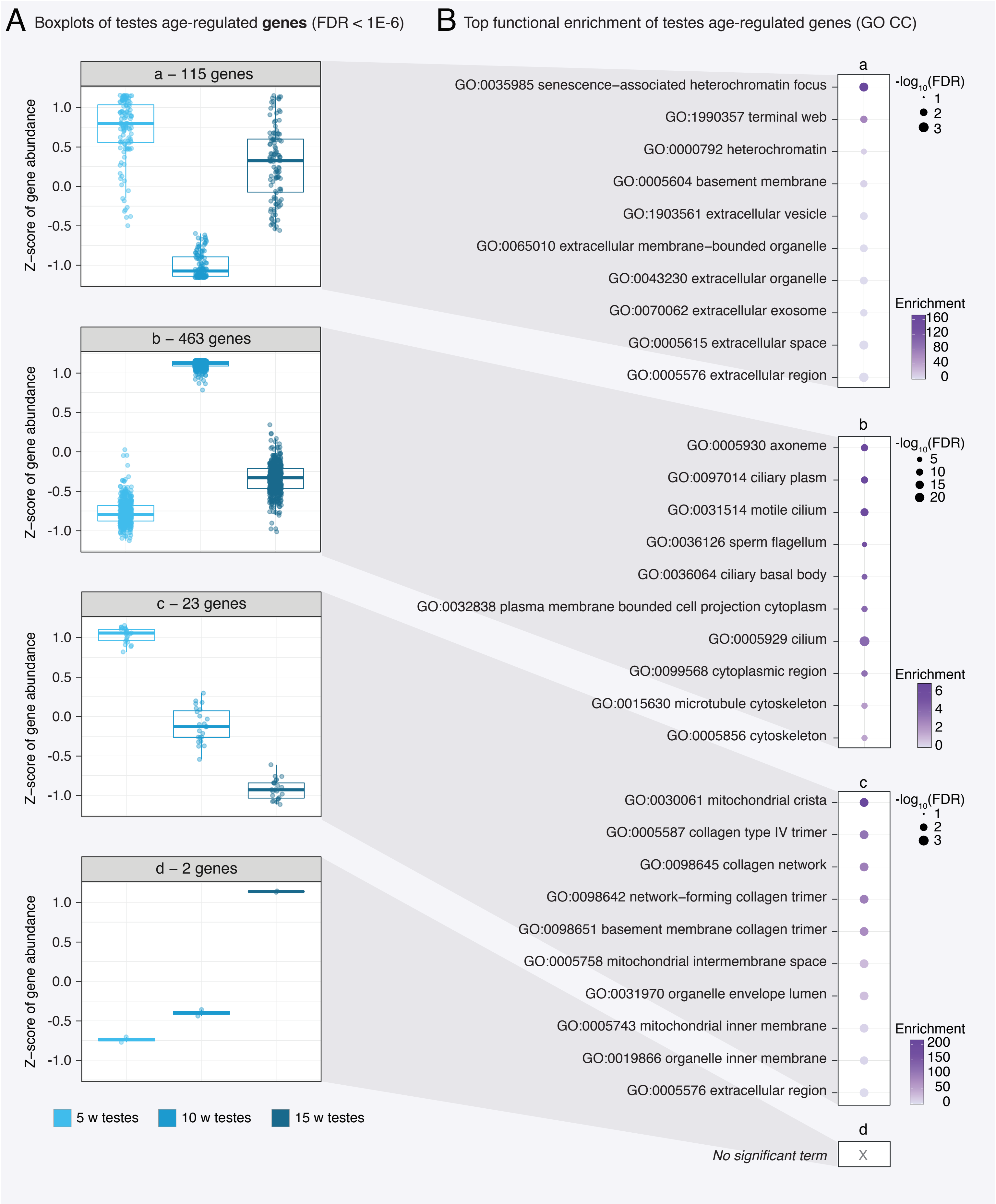

### Supplemental Fig. S4

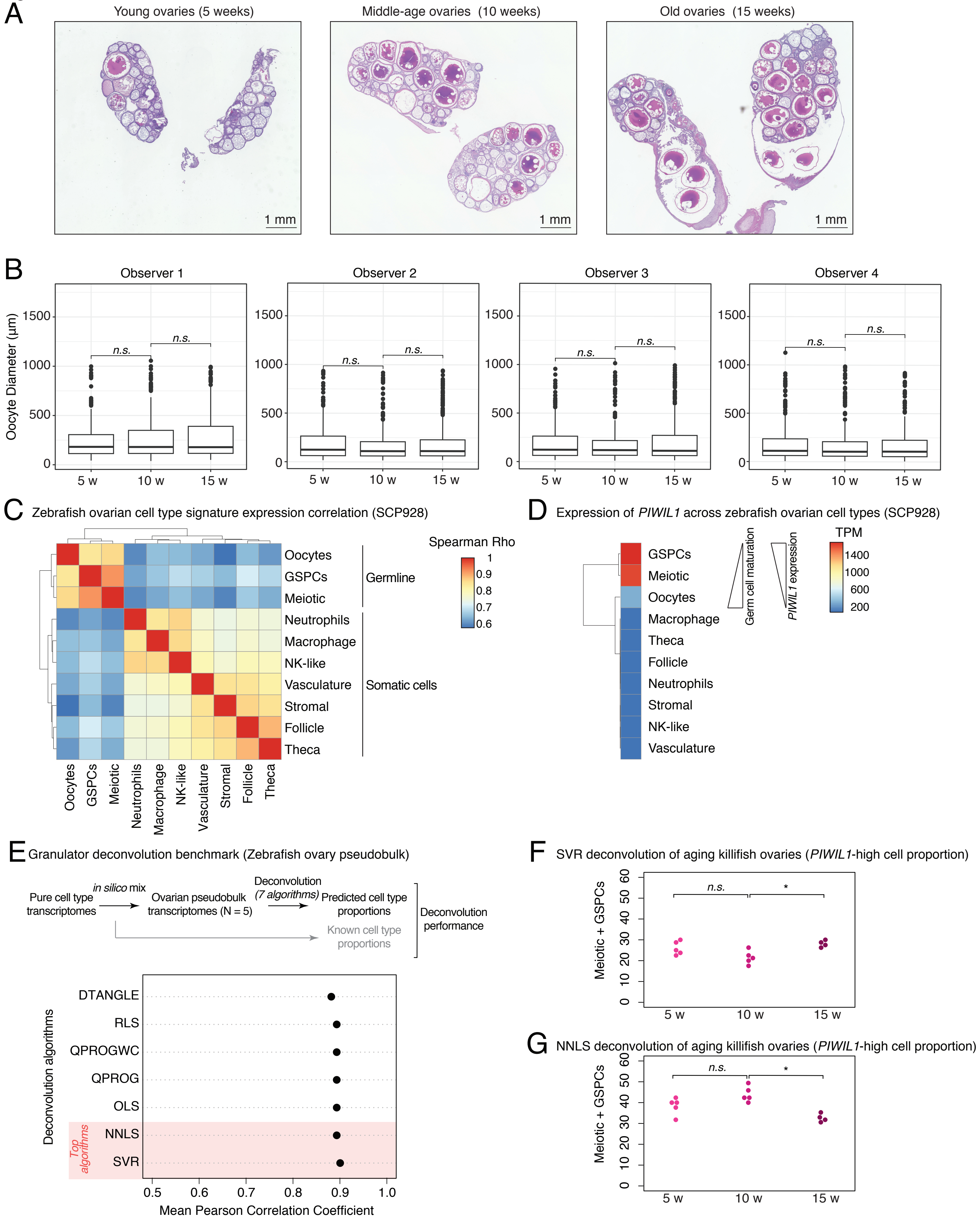

### Supplemental Fig. S5

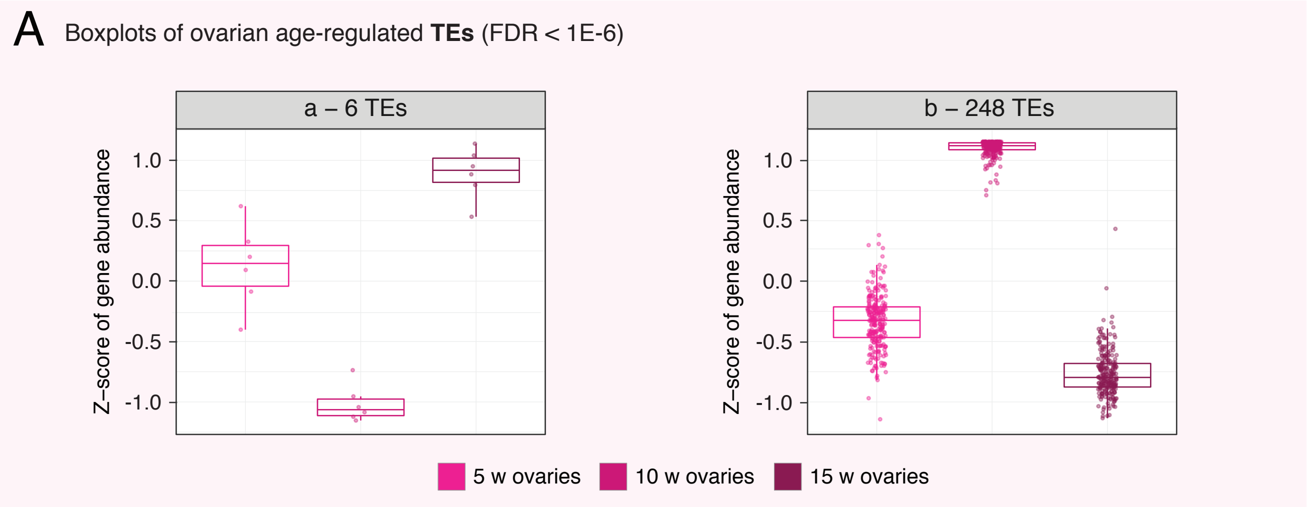

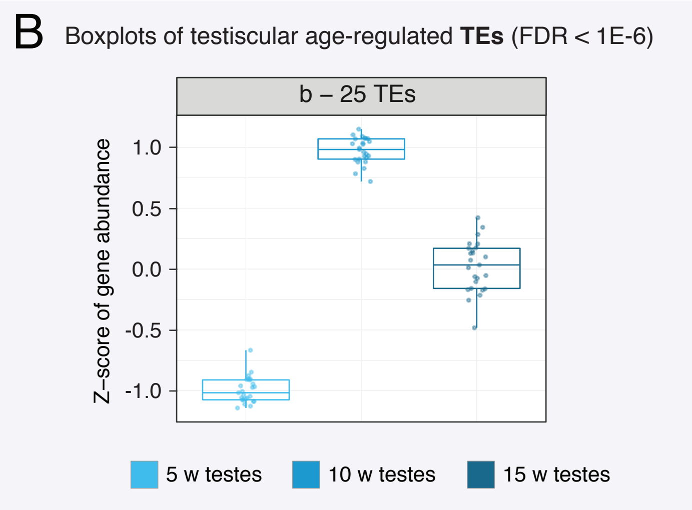

### Supplemental Fig. S6

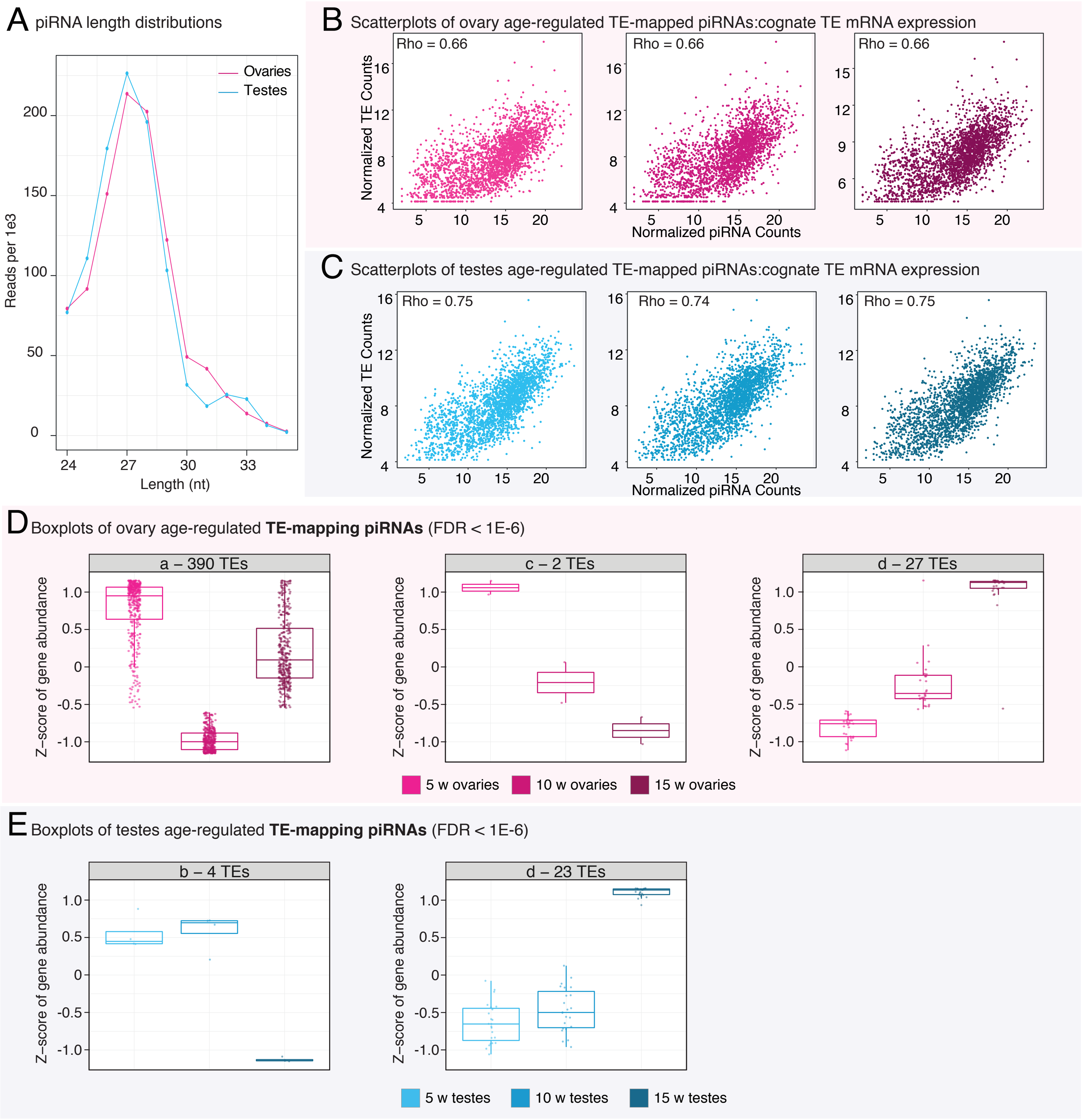

### Supplemental Fig. S7

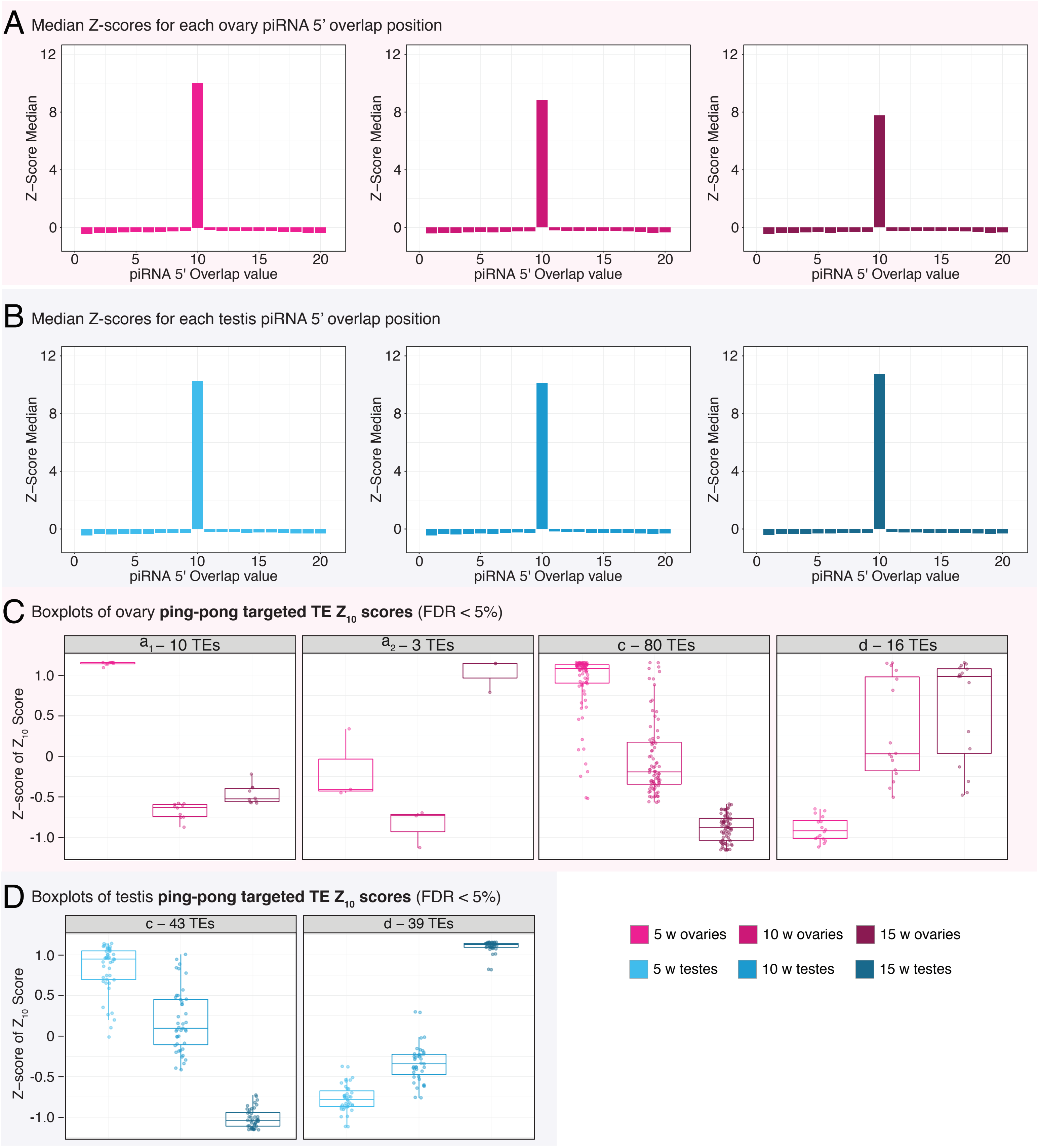
